# Genomic variations and epigenomic landscape of the Medaka Inbred Kiyosu-Karlsruhe (MIKK) panel

**DOI:** 10.1101/2021.05.17.444424

**Authors:** Adrien Leger, Ian Brettell, Jack Monahan, Carl Barton, Nadeshda Wolf, Natalja Kusminski, Cathrin Herder, Narendar Aadepu, Clara Becker, Jakob Gierten, Omar T. Hammouda, Eva Hasel, Colin Lischik, Katharina Lust, Risa Suzuki, Tinatini Tavhelidse, Thomas Thumberger, Erika Tsingos, Philip Watson, Bettina Welz, Kiyoshi Naruse, Felix Loosli, Joachim Wittbrodt, Ewan Birney, Tomas Fitzgerald

## Abstract

The teleost medaka (*Oryzias latipes*) is a well-established vertebrate model system, with a long history of genetic research, and multiple high-quality reference genomes available for several inbred strains (*HdrR*, *HNI* and *HSOK*). Medaka has a high tolerance to inbreeding from the wild, thus allowing one to establish inbred lines from wild founder individuals. We have exploited this feature to create an inbred panel resource: the Medaka Inbred Kiyosu-Karlsruhe (MIKK) panel. This panel of 80 near-isogenic inbred lines contains a large amount of genetic variation inherited from the original wild population. We used Oxford Nanopore Technologies (ONT) long read data to further investigate the genomic and epigenomic landscapes of a subset of the MIKK panel. Nanopore sequencing allowed us to identify a much greater variety of high-quality structural variants compared with Illumina sequencing. We also present results and methods using a pan-genome graph representation of 12 individual medaka lines from the MIKK panel. This graph-based reference MIKK panel genome revealed novel differences between the MIKK panel lines compared to standard linear reference genomes. We found additional MIKK panel-specific genomic content that would be missing from linear reference alignment approaches. We were also able to identify and quantify the presence of repeat elements in each of the lines. Finally, we investigated line-specific CpG methylation and performed differential DNA methylation analysis across the 12 lines. We thus present a detailed analysis of the MIKK panel genomes using long and short read sequence technologies, creating a MIKK panel specific pan genome reference dataset allowing for the investigation of novel variation types that would be elusive using standard approaches.

## Introduction

The Japanese medaka fish (*Oryzias latipes*) has a long history as a vertebrate model organism [1,2]. We took advantage of its unusually high tolerance to inbreeding to establish the Medaka Inbred Kiyosu-Karlsruhe (MIKK) panel: the largest collection of near-isogenic vertebrate lines derived from a single wild population [3]. In the companion article published with this one, we provide a detailed genetic characterisation of the 80 individual MIKK panel lines [4], based on the alignment of Illumina short reads to the closest, fully-assembled reference genome – the southern Japanese medaka inbred strain, *HdrR*. Although this allowed us to discover much of the genetic variation in the MIKK panel relative to *HdrR*, the approach inevitably kept certain variants hidden, including larger and more complex structural variation – “dark variation” – that is likely to have functional consequences for each of the lines. Here we describe how we used Oxford Nanopore Technologies (ONT) long read sequencing to uncover some of this dark variation in 12 of the MIKK panel lines, giving us a more complete assessment of their genomic variation, and paving the way for future studies to elaborate on how structural variants (SVs) affect phenotypes of interest.

The traditional approach for detecting genetic variation is to align reads to a linear reference genome. There are at least three high-quality medaka reference genomes based on inbred strains from different geographical regions in eastern Asia [5,6]. These include *HdrR* (southern Japan), *HNI* (northern Japan), and *HSOK* (Korea), all of which have been characterised in depth at both phenotypic and genomic levels [7,8]. Using such linear reference genomes makes it relatively straightforward to determine the functional consequences of genetic variants relative to those references. Although this reference-anchored approach is convenient, it introduces a “reference bias” that can give rise to an underrepresentation or even incorrect interpretation of genetic variation [9]. Specifically, it makes it difficult to discover complex structural variation, such as large insertions and nested variations.

Variation pangenome graphs offer a compelling alternative approach, allowing for the representation of different classes of SVs using universal semantics [10–13]. The sequencing costs and mapping ambiguity of short reads has so far hindered the widespread adoption of graph genomes. However, recent advances in long-read sequencing technologies [14,15], and the availability of efficient graph assembly algorithms [11,16], now make it possible to generate pangenome graphs from multiple draft assemblies at a reasonable cost. These individual assemblies additionally confer the ability to map and quantify different types of repeats [17,18], which was previously limited when using short read technology alone. Although much progress is being made, this pangenome approach does come with its own limitations, as the graph representation is more difficult to understand and interpret [19]. Nevertheless, in this study we demonstrate how these modern assembly generation and aggregation approaches have allowed for a more complete assessment of genomic variation in 12 of the MIKK panel lines.

Even when applying the traditional reference-anchored approach, using Illumina short read information together with ONT long read information can create a highly-accurate representation of large-scale genomic variation. Illumina short read technology has known limitations for the discovery of large structural variants (SVs), leaving large chunks of the genome poorly characterised [8,20]. It is clear that SVs impact important traits in humans [21] and it is essential to accurately characterise them to gain a more complete picture of the variation between genomes [22]. Using the combination of long and short read information takes advantage of their complementary strengths: long reads can span highly-repetitive regions, helping to resolve complex SVs; whereas short reads are often of higher quality overall, allowing increased base-calling and mapping accuracy when used to polish the long read SV calls. Numerous methods for using both technologies in concert have been developed over the years [23], and although there still remain certain challenges associated with SV detection [24], methods that can leverage the combined information from different modern sequencing technologies are likely to provide the highest SV detection accuracy [25]. Here we show how we used ONT long read information to discover large SVs in 12 MIKK panel lines using the traditional reference-anchored approach, and how polishing the SVs with Illumina short reads substantially improved their mapping accuracy.

Finally, in addition to enabling the construction of pangenome graphs and the discovery of larger SVs, ONT sequencing also allows one to directly detect DNA modifications, such as DNA methylation [26,27]. We used ONT here to characterise DNA methylation in 12 MIKK panel lines. Altogether, we demonstrate the advantages of using combined short and long read technologies, together with both traditional and modern alignment and assembly approaches, in order to more fully characterise large and complex genomic variation. Ultimately, this more complete assessment of the differences between genomes will lead to a more detailed and sophisticated understanding of how genetic variation causes phenotypic differences.

## Results

### Line-specific assemblies and medaka pangenome graph assembly

We selected 12 MIKK panel lines (including 3 pairs of sibling lines) and sequenced brain samples with ONT long read technology to a median of 20x coverage per line, with 37 million reads overall. We multiplexed 4 samples per PromethION flowcell and obtained more than 10 Gb per sample with a mean genome coverage between 13X and 30X as compared with 31X to 39X for illumina sequencing (**Supplementary Table 1**). The median N50 of the reads was 7,411 bp (**Supplementary Table 1**). The analysis consisted of 4 steps: 1) Linear draft assembly for each MIKK line using both short and long reads; 2) Pangenome graph construction combining known medaka reference genomes and MIKK panel draft assemblies; 3) Alignment of ONT reads to the graph; and 4) Extraction of complex structural variations.

### Individual MIKK line assemblies

We generated individual assemblies for each line using a hybrid Illumina/ONT strategy. To this end, we first built a scaffold with ONT data using wtdbg2 [28] and then polished the resulting assemblies using the Illumina reads with Pilon [29]. The quality of the draft assemblies was evaluated with Quast [30] against the *HdrR* reference assembly. The assemblies have between 2500 to 4400 contigs amounting to total lengths of 721 Mb to 742 Mb, with N50 values between 404 kb and 971 kb (**Table 1** and **Figure 1A**). Assembly lengths are highly consistent with the length of the medaka *HdrR* reference (734 Mb), as are the percentages of CG (**Table 1** and **Figure 1D**). However, when aligning the contigs to the *HdrR* reference the median alignment length (NA50) scores drop to values between 105 to 280 kb, although many alignments are over 1 Mb long. This is very likely due to the presence of structural variations interrupting alignments and a significant divergence of the MIKK genomes as compared with *HdrR*. Indeed, on average only 80% of the bases from the MIKK panel genomes are aligned unambiguously to the *HdrR* reference and a similar trend is observed for the number of genes covered (**Table 1** and **Figure 1C,E**). As shown in more details in the following section (**Figure 2B**), the majority of the additional 20% present in the MIKK panel occurs in more than one MIKK line. Altogether, this suggests that the MIKK panel line genomes can be reasonably accurately assembled and contain a significant amount of genetic diversity compared with the HdrR line, the closest complete reference assembly.

**Table 1.** Summary statistics of individual MIKK lines assemblies

| Line id | Number of contigs | GC (%) | Total length | Largest contig | N50 | Total aligned length | Largest alignment | NA50 |
| --- | --- | --- | --- | --- | --- | --- | --- | --- |
| 4-1 | 2,886 | 40.66 | 730,816,425 | 5,635,124 | 802,725 | 599,170,757 | 2,340,443 | 259,311 |
| 4-2 | 2,762 | 40.71 | 737,637,241 | 6,376,848 | 971,613 | 612,988,975 | 2,781,976 | 279,257 |
| 7-1 | 2,512 | 40.69 | 732,447,291 | 5,851,261 | 942,347 | 608,014,583 | 2,243,072 | 265,102 |
| 7-2 | 2,892 | 40.69 | 732,448,405 | 4,099,264 | 845,096 | 607,409,964 | 1,761,859 | 253,015 |
| 11-1 | 3,368 | 40.56 | 728,542,858 | 4,525,370 | 624,727 | 545,652,612 | 1,541,261 | 180,262 |
| 69-1 | 3,077 | 40.59 | 727,390,278 | 6,080,511 | 708,738 | 573,096,833 | 1,823,612 | 220,342 |
| **79-2** | 3,053 | 40.62 | 730,357,166 | 6,658,276 | 742,721 | 584,535,579 | 2,384,437 | 235,007 |
| **80-1** | 4,374 | 40.49 | 720,948,860 | 3,238,833 | 404,886 | 491,501,885 | 1,304,457 | 105,372 |
| **117-2** | 2,810 | 40.73 | 732,113,747 | 5,059,334 | 903,899 | 620,809,739 | 2,195,961 | 270,130 |
| **131-1** | 3,651 | 40.72 | 741,963,499 | 5,589,066 | 737,055 | 563,763,739 | 2,231,101 | 180,572 |
| **134-1** | 3,142 | 40.71 | 731,842,804 | 4,138,056 | 747,064 | 606,908,341 | 2,042,573 | 245,116 |
| **134-2** | 3,977 | 40.75 | 729,131,624 | 3,835,876 | 475,444 | 621,355,180 | 1,698,873 | 209,441 |

**Figure 1.**
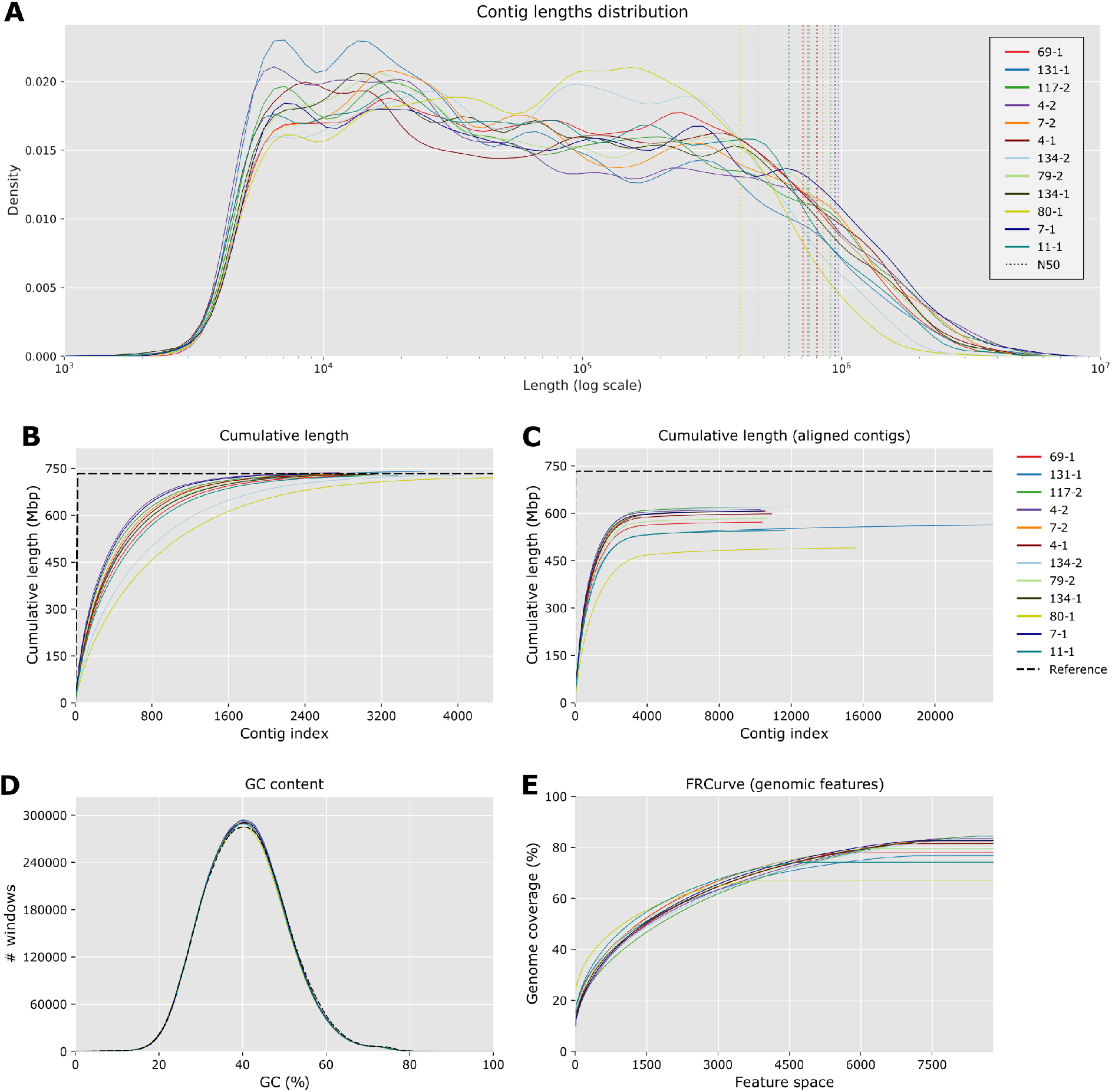
Quality metrics for individual assemblies. **A**. Normalised distribution of contigs length for each assembly. Dashed lines represent the N50 values. **B**. Cumulative length of contigs and **C**. Cumulative length for contig blocks aligned on *HdrR*, in comparison with the *HdrR* reference chromosomes (dashed black line). **D**. Distribution of CG content of assemblies in comparison with the *HdrR* reference (dash black line). **E**. Feature-Response Curve for *HdrR* genes annotation, showing the quality of the assemblies as a function of the maximum number of possible genes allowed in the contigs.

**Figure 2.**
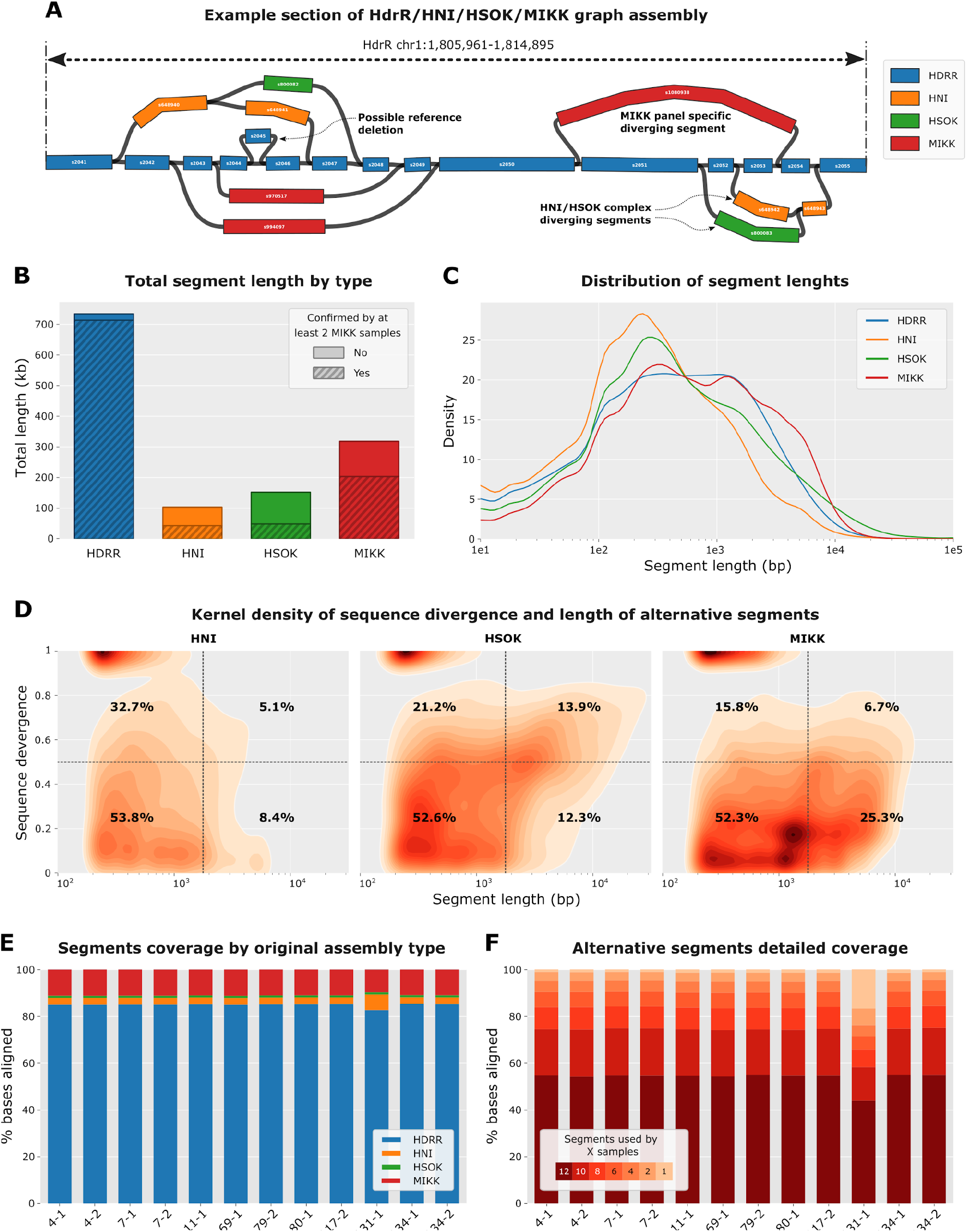
Pangenome graph reference characterisation. **A**. Example section of the graph for chromosome 1 showing different paths through the graph via segments originating from the 4 types of assemblies used to build the graph. **B**. Total length of segments contained into the graph by type of assembly. Dashed areas represent the proportion of bases for segments covered by at least 2 samples with at least 5% of the average coverage over the *HdrR* reference segments. **C**. Distribution of the length of segment by type of assembly normalised by the total length of segments. **D**. Kernel density plots of the length of alternative segments according to their divergence when aligned onto the *HdrR* reference. The quadrants defined by vertical dashed lines (length = 2kb) and horizontal dashed lines (divergence = 0.5) separate the segments into 4 categories according to their length and divergence score. The numbers displayed correspond to the percentages of segments within each of the 4 quadrants. **E**. Percentages of bases from nanopore reads aligning on each type of assembly for the 12 MIKK samples. **F**. Detailed percentages of bases aligned on alternative segments for each MIKK sample depending on segments cross-usage by the other samples, from 12 (all other samples) to 1 (only the current sample). A segment was considered used by a sample when its coverage was at least 5% of the average *HdrR* reference coverage.

### Pangenome graph assembly and read alignment

To better represent the complexity of the MIKK panel and the relationships with existing medaka reference genomes, we built a pangenome variation graph with minigraph [11] containing all the individual MIKK assemblies together with the *HdrR*, *HNI* and *HSOK* reference assemblies (http://utgenome.org/medaka_v2). Previous phylogenetic analyses showed that the MIKK panel is genetically closest to *HdrR*, then *HNI* and finally *HSOK* [3]. Thus, we used an iterative strategy to build our pangenome graph, starting with *HdrR* as the primary anchor, followed by *HNI*, then *HSOK* and finally all the MIKK panel assemblies one by one. By doing so, we can identify the segments of the graph that are specific to the MIKK panel, while having information on which is the closest reference for every graph segment. The presence of non-*HdrR* alternative segments can indicate insertions or significant divergence from the reference, whereas missing *HdrR* segments are indicative of possible deletions (Example shown graphically in **Figure 2A**). We obtained a graph containing over 1.1 million segments totaling 1.3 Gb, which is around 1.8 times larger than the *HdrR* reference genome. Together, the 12 MIKK lines bring an additional 211,836 segments to the graph (318 Mb, 24.3% of total) of which 161,533 are covered by at least 2 lines (203 Mb, 20.2%. The segments only found in the MIKK panel have an N50 of nearly 4000 bp and a median sequence identity of 73.7% when aligned to the *HdrR* reference (**Table 2** and **Figure 2B–D**). In summary, the MIKK panel contains a large number of relatively low divergent paths through the graph mostly consisting of segments ranging from 100 to 5000 bp. In comparison both *HNI* and *HSOK* bring fewer segments but with a greater sequence divergence as compared with the *HdrR* reference (62.9% and 60.4% identity respectively). Interestingly, *HSOK* has a sizable population of long and divergent segments (> 2000 bp and > 0.5 divergence) which represent 13.9% of segments as opposed to 5.1% and 6.7% for *HNI* and MIKK, respectively (**Figure 2D**). This is in line with the established phylogeography of Medaka fish, in that the Korean derived HSOK line is geographically isolated from Japanese medaka lines and earlier branching in evolutionary time.

**Table 2.** Pangenome graph reference statistics. Segment type indicates which assembly the segments originally come from. For the “Segments used by at least 2 MIKK samples’’ columns we defined a segment as being used if its coverage is at least 10% of the average coverage over the *HdrR* reference segments.

| Segment type | All segments in graph |  | Segments used by at least 2 MIKK samples |  | Median segment length | longest segment | N50 | Median % identity |
| --- | --- | --- | --- | --- | --- | --- | --- | --- |
|  | Length (bp) | # segments | Length (bp) | # segments |  |  |  |  |
| *HdrR* | 734,100,826 | 648,692 | 713,609,808 | 615,564 | 401 | 675,459 | 3000 | NA |
| *HNI* | 103,507,879 | 148,689 | 43,204,187 | 47,854 | 239 | 175,667 | 2003 | 62.9% |
| *HSOK* | 152,043,332 | 100,275 | 49,055,247 | 23,881 | 371 | 236,527 | 5803 | 60.4% |
| MIKK | 318,174,656 | 211,836 | 203,539,620 | 161,533 | 559 | 89,792 | 3998 | 73.7% |
| All | 1,307,826,693 | 1,109,492 | 1,009,408,862 | 848,832 | 389 | 675,459 | 3342 | NA |

Finally, we analysed the graph usage after aligning raw Nanopore reads for each individual MIKK sample and computing the coverage for each segment. Overall, the MIKK lines behave similarly in terms of the reference types to which they align, with *HdrR* holding the bulk of the coverage (median = 85.1%) followed by MIKK specific segments (11%) *HNI* segments (3%) and *HSOK* (0.9%) (**Figure 2E** and **Supplementary Table 2**). Among the non-*HdrR* alternative segments, we also investigated the cross-usage of each segment across the 12 MIKK samples. The samples overwhelmingly use segments that are also covered by at least half of all the samples (median = 90.36%) and even by all the samples in the majority of the cases (54.61%) **(Figure 2D** and **Supplementary Table 3**). However, there is one notable exception for line 131-1 for which *HNI* type segments get a much larger fraction of the coverage (6.8%). The samples also tend to align on alternative segments supported by fewer samples, with 28.6% of the bases aligned on segments used by fewer than 6 samples, including 16.7% specific to 131-1 line (see discussion and **Supplementary File 1**).

### Novel genetic sequences and large scale insertions and deletions in the MIKK panel

Pangenome variation graphs offer new options to discover structural variations that are not available with conventional SV approaches based on linear reference genomes. In particular, they are better suited to represent genomic intervals which accumulated a large number of small variations as divergent alternative paths. We analysed the presence of such paths in our medaka pangenome graph and their potential functional impact. To do so we identified branches of the graph containing segments which have: 1) a low identity compared with the *HdrR* reference; 2) a robust DNA-Seq and RNA-Seq support from multiple MIKK panel samples; 3) a total cumulative length exceeding 10 kb; and 4) with at least one annotated exon overlapped (see precise criteria in **Methods**). With this strict set of criteria we found 19 such alternative paths in our graph (**Supplementary Table 5**). The 2 examples presented in **Figure 3A**/**B** and **Supplementary Table 4** show the layout of the graph with the reference and the alternative divergent paths. To investigate the precise RNA usage pattern, we generated local linear assemblies for the 2 branches of each selected loci, and aligned short RNA-Seq data obtained for 50 MIKK liver samples. In both cases, the exonic coverage pattern over the reference and alternative paths is strikingly different, showing the impact on the transcriptional landscape around these loci of the structural variation.

**Figure 3.**
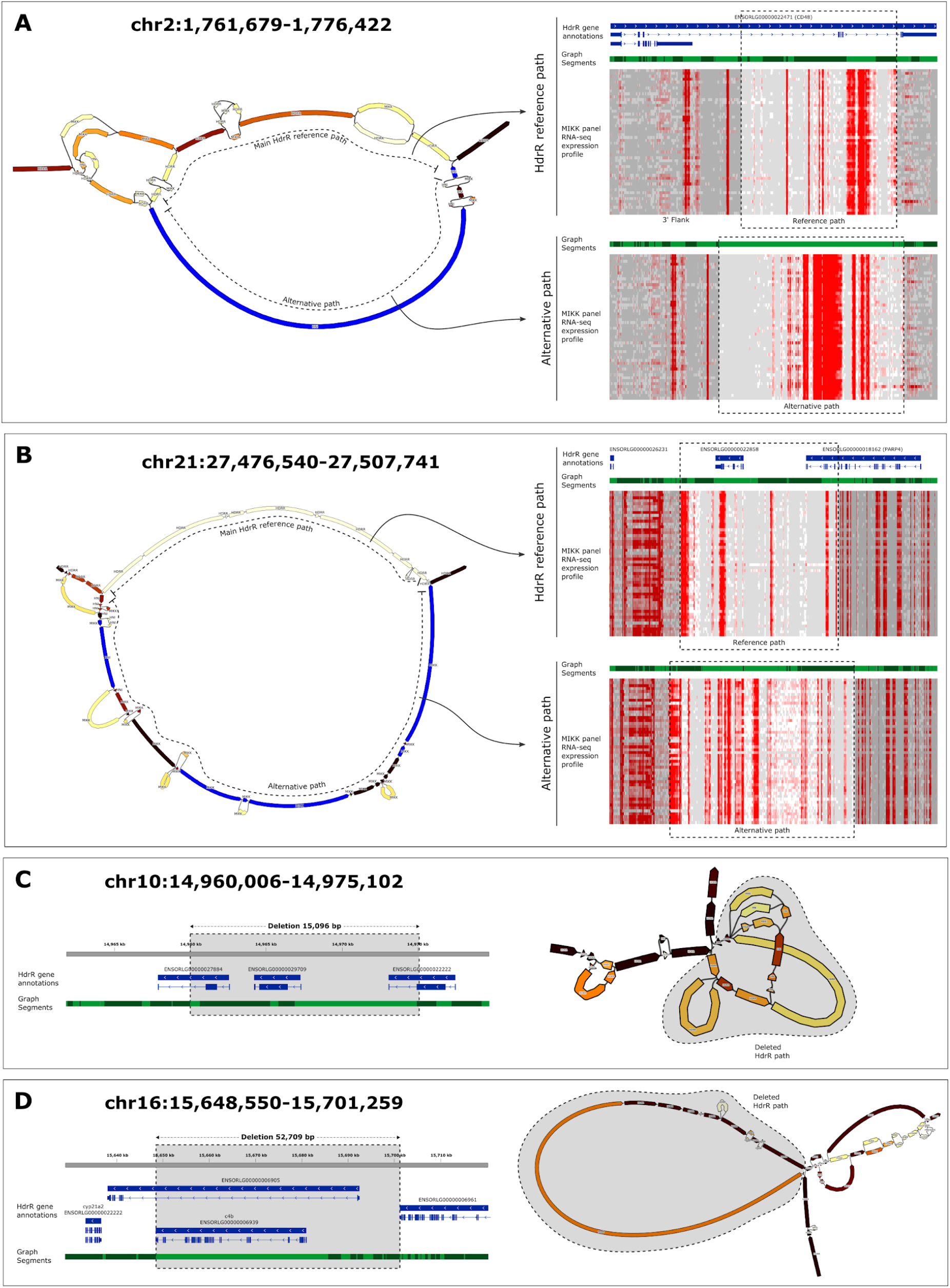
Example structural variations identified in the pangenome graph. **A/B**. Visualisation of alternative divergent paths in the graph. For both selected examples, the left side panel shows a Bandage plot indicating the reference *HdrR* and the alternative path. Each graph segment is color-coded according to the number of samples with at least 50% of the reference coverage for the ONT DNA-Seq dataset, from white (None) to deep red (All). Blue segments are supported by multiple samples for both Illumina RNA-Seq and ONT DNA-Seq. The right panel shows the linear structure of local assemblies for both the reference and the alternative paths. For the reference the top blue track represents the existing Medaka *HdrR* annotations. The light and dark green tracks correspond to the segments layout from the graph. Finally, the heatmaps show the RNA expression intensities for all 50 medaka samples sequenced along the represented sections of the graph (grey = not found, white = less than 5 reads, dark red = more than 100 reads). **C/D**. Visualisation of large scale deletions in the graph. For both selected examples, the left panel shows the Medaka *HdrR* annotations (blue) and the graph segment layout (light and dark green), overlaid with the deletion position (grey rectangle). The Bandage plots on the right are color-coded as previously described. The shaded area indicates the reference sequence deletions robustly supported by a direct connection between distant reference segments (Link coverage > 50% of reference coverage for at least 9 samples).

Large scale rearrangements, deletions in particular, can easily be detected in a graph by analysing the usage of links between segments. We selected links 1) connecting 2 *HdrR* segments distant by more than 10kb; 2) with a strong coverage in multiple MIKK panel samples; and 3) skipping at least 1 annotated exon (see precise criteria in **Methods**). We obtained a list of 16 of these large scale deletions (**Supplementary Table 4**), 2 examples of which are presented in **Figure 3C/D**. When lowering the length to 1 kb and not restricting to deletions overlapping exons, we found 2059 such events, showing the widespread occurrence of such deletions with a disruptive potential.

Altogether, our graph genome analysis generated a comprehensive dataset from these 12 lines that has allowed us to identify complex variants. We were able to highlight potential functional consequences including disruption of genes exonic pattern and removal of entire genes. Further computational tools in read mapping and annotation will be needed to robustly identify non-reference genic content in the graph genome. This new way to look at population genomics allowed us to visualize highly complex SVs in medaka fish at an unprecedented resolution and to provide our graph assembly as a resource for the community.

### Structural variation and breakpoint mapping in the MIKK panel

As an alternative to the variation pangenome approach, we also explored structural variation (SVs) in a reference-anchored manner, similar to many human studies. Differences in SVs between panel lines is another important class of genetic variation that could cause or contribute to significant phenotypic differences. Having sequenced the entire panel using short reads we were able to accurately characterise copy number variation (CNV) genome-wide for all panel lines (MIKK panel paer). Here we also used Nanopore sequencing data obtained for 9 of the 12 selected lines allowing us to further characterise larger SVs in the MIKK panel and to create a more extensive picture of genomic rearrangements compared to available medaka reference genomes. We first called structural variants using only the ONT long reads, producing a set of structural variants classified into five types: deletions (DEL), insertions (INS), translocations (TRA), duplications (DUP) and inversions (INV). We then “polished” the called DEL and INS variants with Illumina short reads to improve their accuracy. The polishing process filtered out 7.4% of DEL and 12.8% of INS variants, and adjusted the breakpoints (i.e. start and end positions) for 75-77% of DEL and INS variants in each sample by a mean of 23 bp for the start position, and 33 bp for the end position (**Supplementary Figure 1**). This process produced a total of 143,326 filtered SVs.

The 9 “polished” samples contained a mean per-sample count of approximately 37K DEL variants (12% singletons), 29.5K INS variants (14%), 3.5K TRA variants (9%), 2.5K DUP (7%) and 600 INV (7%) (**Figure 4D**). DEL variants were up to 494 kb in length, with 90% of unique DEL variants shorter than 3.8 kb. INS variants were only up to 13.8 kb in length, with 90% of unique INS variants shorter than 2 kb. DUP and INV variants tended to be longer, with a mean length of 19 and 70.5 kb respectively (**Figure 4A**). **Figure 4E** shows the per-sample distribution of DEL variants across the genome. Most large DEL variants over 250 kb in length were common among the MIKK panel lines. A number of large DEL variants appear to have accumulated within the 0-10 Mb region of chromosome 2, which is enriched for repeats in the *HdrR* reference genome (**Supplementary Figure 2**).

**Figure 4.**
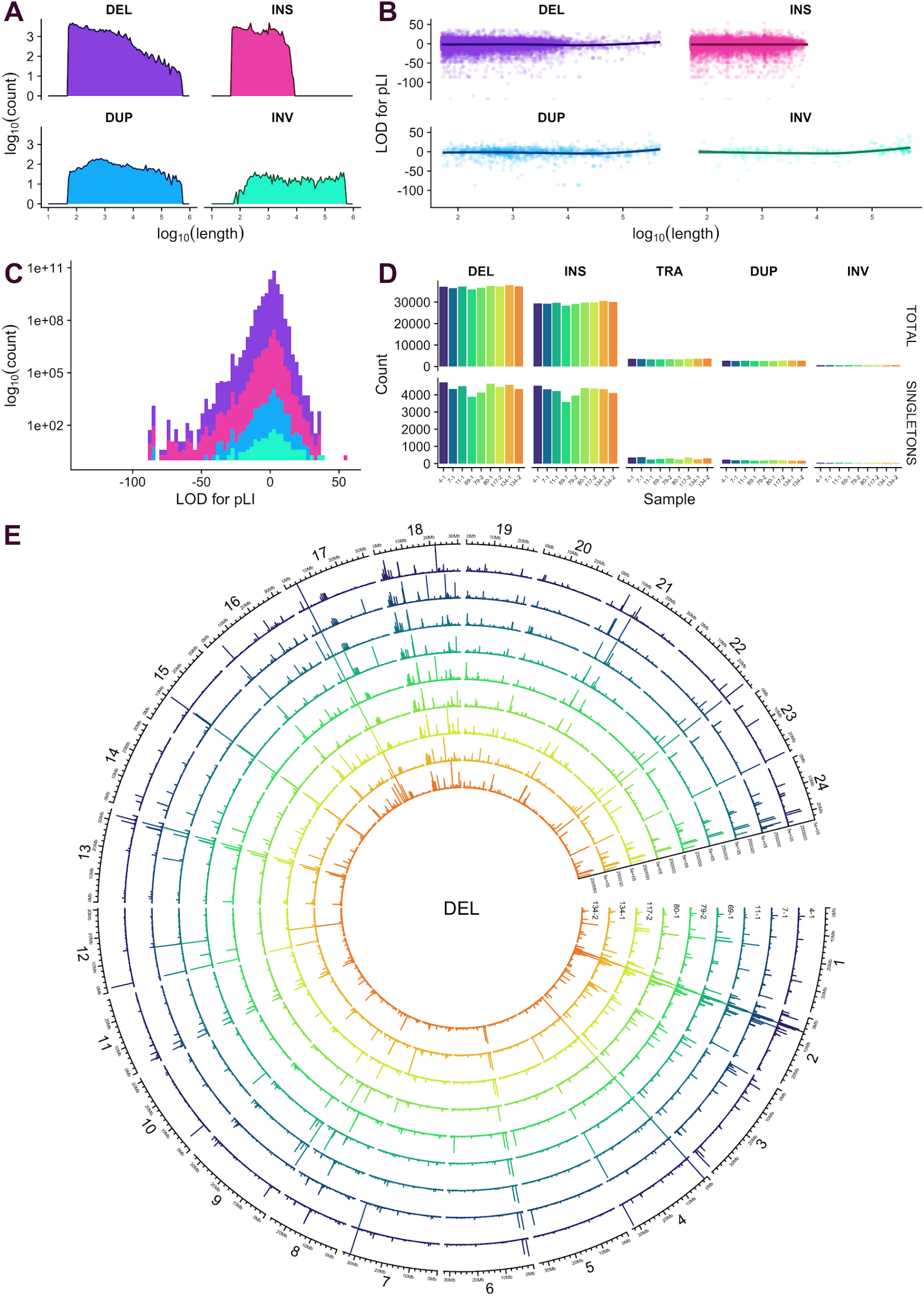
Polished SVs in 9 MIKK panel lines sequenced with ONT. *DEL:* deletion; *INS:* insertion; *TRA* : translocation; *DUP*: duplication; *INV*: inversion. **A**. Aggregate log_10_ counts and lengths of distinct SVs by type, excluding TRA. **B**. pLI LOD scores in distinct SVs by SV type. **C**. Histogram of LOD scores by SV type. **D**. Total and singleton counts of SV types per sample. **E**. Circos plot showing per-sample distribution and lengths of DEL variants across the genome. Circos plots for each of the other SV types are included in **Supplementary File 2**.

SVs were generally enriched in regions covered by repeats. While only 16% of bases in the *HdrR* reference were classified as repeats (irrespective of strand), those bases overlapped with 72% of DEL, 63% of DUP, 81% of INV and 35% of TRA variant regions. However, repeat bases only overlapped with 21% of INS variants. We also assessed each SV’s probability of being loss-of-function (pLI) [31] by calculating the logarithm of odds (LOD) for the pLI scores of all genes overlapping the variant (**Figure 4B,C**). 30,357 out of 134,088 DEL, INS, DUP and INV variants overlapped at least one gene, and 9% of those had a score greater than 10, indicating a high probability that the SV would cause a loss of function. Two INS variants on chr2 had an outlying LOD score of 57 as a result of overlapping medaka gene ENSORLG00000003411, which has a pLI score of 1 – the highest intolerance to variants causing a loss of function. This gene is homologous with human genes *SCN1A*, *SCN2A* and *SCN3A*, which encode sodium channels and have been associated with neuronal and sleep disorders. We did not find evidence that longer SVs tended to have a higher probability of causing a loss of function (**Figure 4B**).

### Differential methylation analysis

Native DNA sequencing with nanopore technology can be used to robustly detect CpG methylation at single base and single read resolution. There are a number of software methods which can be used to identify methylated positions [26,27], however finding differentially methylated areas of the genome between samples requires processing these methylation calls across samples. To do so we developed an analytic framework [32] to identify CpG islands of interest in the 12 panel lines sequenced with Nanopore DNA-Seq.

We were able to use the sequencing data previously generated to explore CpG methylation differences in the brain samples of the 12 selected MIKK panel lines. We found 4,459 significantly differentially methylated regions (DMR) among the 271,294 CpG islands included in the analysis (FDR=1%), using a Kruskal Wallis test. Significant DMR are distributed across the entire genome, with a possible enrichment towards chromosomal extremities (**Figure 5C**). In addition, we observed a sharp enrichment of significant DMR near genes transcription start sites compared with non-significant regions (**Figure 5B**). When clustering the significant DMR across the entire genome the 3 sib-lines pairs included in the analysis cluster together. This suggests that the methylome is conserved across multiple generations; the simplest explanation being that much of the methylation variation is genetically determined. Among the top hits, we found interesting candidates in close proximity to coding genes including *onecut3a*, *gart*, *dnase1* and *galm*. Detailed interactive reports for the 100 top significant DMR are available at https://birneylab.github.io/MIKK_genome_companion_paper/DNA_methylation/results/pycometh_html/pycoMeth_summary_report.html and a list of all significant hits is provided as supplementary material (Supplementary File 3).

**Figure 5.**
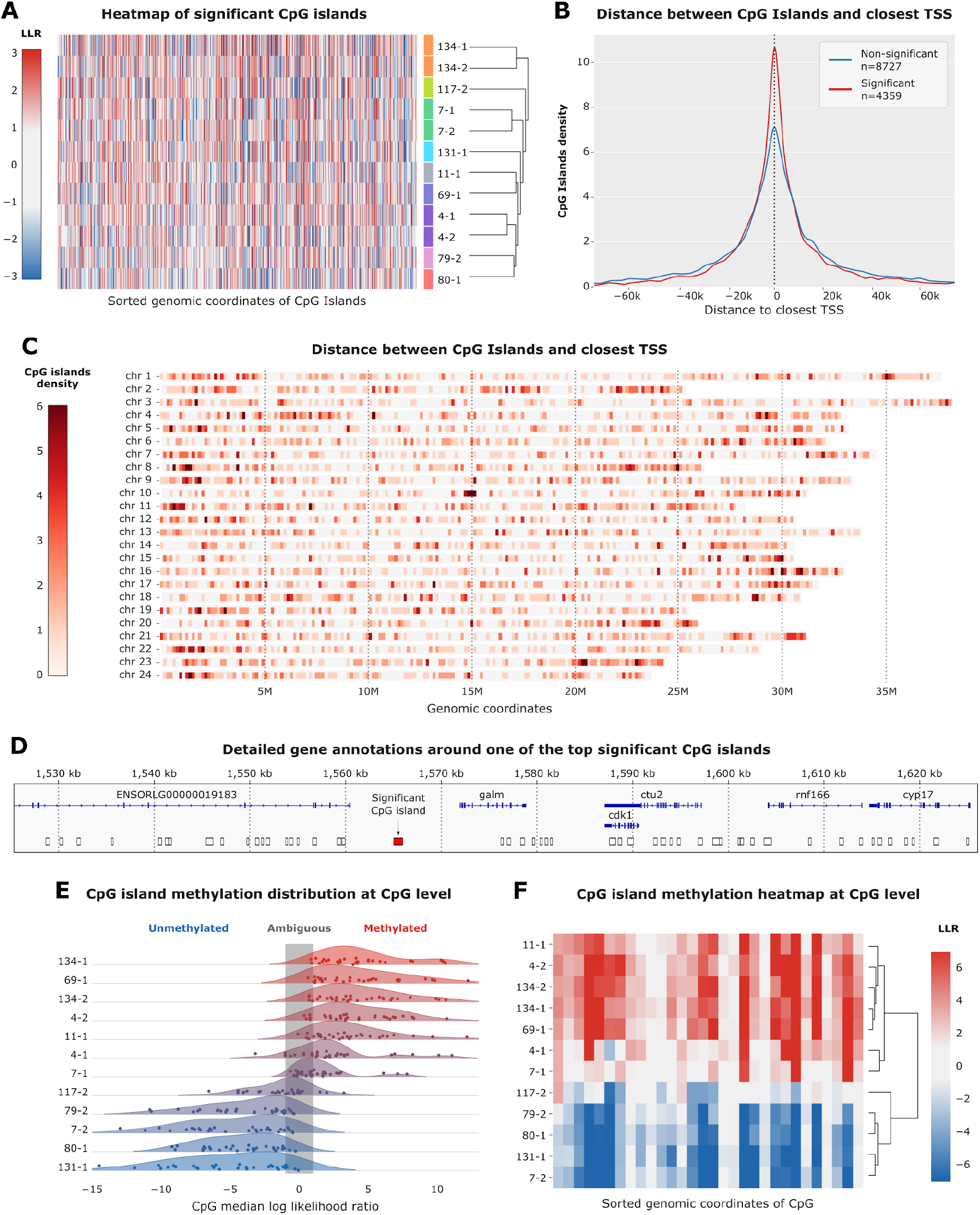
DNA methylation analysis. **A**. Heatmap of all significant CpG islands differentially methylated across the MIKK panel samples. CpG islands are sorted by genomic positions and the X axis. Samples are ordered by hierarchical clustering on the Y axis and color coded so that sibling lines are indicated with the same colors. The color scheme is according to the value of the median log-likelihood ratio from −3 (blue) to 3 (red) with ambiguous values between −1 and 1 in white. **B**. Distribution of distances to the closest gene TSS for significantly differentially methylated CpG in red (n=4249) and all non-significant islands in blue (n=8727). **C**. Number of significant CpG islands by genomic windows for the 24 HdrR chromosomes, from 0 (white) to 6 (dark red). **D**. Example 100kb region containing a significant CpG island in red (chr15:1565040-1565987) as opposed to non-significant ones in white. **E**. CpG level log-likelihood ratio kernel density plot for the CpG island highlighted in panel D. Samples are sorted on the Y axis by decreasing median llr. Individual CpG values are indicated by dots. **F**. Heatmap of log-likelihood ratio with hierarchical clustering by sample for the CpG island highlighted in panel D. On the X axis are individual CpGs sorted by genomic position.

## Discussion

Improvements to the accessibility and affordability of long read genome sequencing technologies opens up new possibilities for a deeper genome characterisation of eukaryotic genomes, and a more complete understanding of intra-species genetic variations. Standard linear reference alignment approaches can only partially handle large and complex structural variation, and a considerable fraction of genetic variation between individuals is masked from sight. Such ‘dark’ genome variation comes in a variety of flavors and scales, including large scale novel sequence insertions, gene conversion and introgression from other compatible genomes.

Here we focused on providing a more complete view of the genetic variation observed across 12 members of the MIKK panel by assembly and integration of all draft genomes anchored to the three high quality medaka reference genomes (*HdrR*, *HNI* and *HSOK*). Each of the 12 MIKK panel members are inbred medaka samples derived from the same wild founder population in Kiyosu Japan and can be considered as a southern strain most closely related to the *HdrR* reference. First we set out to create high quality draft genomes for each of the 12 MIKK lines using a combination of high coverage ONT and Illumina sequence data from the same samples. Here we observe overall good assembly quality metrics with the total sequence lengths being consistently close to the length of the *HdrR* reference genome. Interestingly when aligning these draft assemblies against the *HdrR* reference NA50 values see a marked decrease indicating that there are likely to be a considerable fraction of MIKK panel genomes that are not present in the *HdrR* reference. When aligning these draft references to *HdrR* we often observed large sequences being split up and fragmented into many smaller sections mapping to different genome locations, suggesting a marked divergence in MIKK panel genomes as a result of many structural variations or novel insertions interrupting the *HdrR* sequence. As more and more high quality reference genome assemblies are generated for medaka [33] and other species [34] new approaches for genome representation and comparison are becoming increasingly valuable to allow a deeper understanding of genetic variation within and between species. To further characterise some of the genetic differences across the MIKK panel and the medaka reference genomes we used a recently developed graph based alignment approach, allowing us to represent all 12 MIKK assemblies on the same genome graph together with 3 reference medaka assemblies. Not only does this approach allow us to represent all MIKK panel assemblies on the same scaffold but also provides an intuitive way of assessing genome differences compared to strains considered to be more distant.

The MIKK panel lines, being southern medaka are expected to be most closely related to the *HdrR* reference and this is clearly reflected when looking at the total sequence length within the graph that can be assigned to the *HdrR* reference genome, however there are an appreciable number of MIKK panel specific sequences, the majority of which are supported by 2 or more MIKK panel assemblies. These high confidence novel MIKK panel specific sequences would be masked when using standard linear genome alignment approaches and thereby missing (or incorrectly represented) during downstream analysis, resulting in the best case incomplete genome variation calls. Furthermore, when looking at novel sequences from the two more distant reference genomes, interestingly we observe that the northern *HNI* strain contributes less novel sequences to MIKK panel assemblies than the southern Korean *HSOK* strain. However for high confidence sequences (those supported by 2 or more MIKK panel assemblies) the total length of sequences are approximately equal for *HNI* and *HSOK*. This pattern of sequence sharing compared to the three genomes references is remarkably consistent between all MIKK panel assemblies with the exception of line 131-1, which shows higher numbers of sequences assigned to *HNI* which may have been due to introgression of *HNI* in the facility during the inbreeding process. Overall the pattern of sequence sharing between MIKK panel lines and the three reference genomes is highly similar between lines, consistent with the founder population location (Kiyosu) and we see little evidence of significant introgression from more distant strains.

It is notable that there are clear genetic consequences for some of the structural variations observed, with a conservative set of 74,271 novel contigs that alter gene content and 11,448 that overlap exonic regions. Having a comprehensive set of functional changes on haplotypes, including these complex variations, is critical in the process of understanding the functional impact of variation. This will be of great importance when using the MIKK panel for genetic association studies of phenotypes, where a full catalog of structural variation will facilitate to attribute a mapped genetic locus to the correct functional gene. As well as allowing a more complete representation of genome structure, the detailed characterisation of genome variation using advanced assembly aggregation approaches provides important information that can be utilised to further refine our understanding of gene organisation and function. By using a graph based approach combined with RNA sequence alignment we were able to show distinct expression profile patterns between a standard linear view (or the *HdrR* path) compared to MIKK panel specific alternatives for expressed genes. This analysis also shows the value in a graph genome approach to understand functional impacts of structural variation; it is hard to even represent some of this variation against a single linear genome, and certainly the “nested” variation, where one has variation inside of a large variation to the reference, is virtually impossible to handle. Variation population graphs are clearly more appropriate at least for medaka fish variation, and most likely for all genomes.

One very interesting aspect of DNA sequencing using Oxford Nanopore (ONT) is that along with providing long reads suitable for reference assembly it is also possible to detect DNA modifications, primarily DNA methylation at single base resolution. We were able to detect thousands of differentially methylated bases using ONT sequence data across 12 MIKK panel lines providing a further deep characterisation of the genome variation in medaka fish. Interestingly, methylation patterns appear consistent across multiple generations, with the MIKK panel sibling lines clustering together based on their methylation profiles alone. This suggests that, like in other species [35] methylation is a heritable trait in medaka fish capable of persisting across multiple generations and is likely to impact phenotypic traits.

With this pilot set of 12 MIKK panel lines we have demonstrated the feasibility of generating independent assemblies for the MIKK panel and the rich functional differences present in the structural variation between MIKK individuals. Without these long read based assemblies and subsequent variation population graph we would have been ignorant of the substantial differences in genomic content between the lines. Although there is still a long way to go to make pangenome variation graph a robust alternative to linear genome, there is no doubt that they will play a central role in the ever expanding catalogues of individual genomes sequenced. In addition we have shown here that the DNA modification detection by Oxford Nanopore works robustly in medaka fish. This already provides a useful resource to explore functional differences, but most importantly gives confidence that complete MIKK panel sequencing by Oxford Nanopore will be useful, both for structural variation studies and also providing a key intermediate molecular readout via methylation status. Thus we present here a methodology for genome analysis that has the potential to become a standard approach for population studies.

## Acknowledgements

The authors acknowledge Thomas Seitz, and Irina Oks and Marzena Majewski for supporting the fish husbandry, and Alicia Günthel, Rachel Müller and Beate Wittbrodt for laboratory assistance. The work was funded by the Helmholtz funding programme BIFTM to F. Loosli, N. Wolf, N. Kusminski, C. Herder and N. Aadepu. E. Birney, T. Fitzgerald, A. Leger, C. Barton, J. Monahan and I. Brettell were funded by the EMBL European Bioinformatics Institute (EMBL-EBI). This work was supported by Heidelberg University Core Funding to T. Tavhelidse, T. Thumberger and J. Wittbrodt. This project has received funding from the European Research Council (ERC) under the European Union’s Horizon 2020 research and innovation programme (grant agreement No 810172), from the NIH UH-3338-03 (JW), the German Ministry for Research (BMBF: HIGH-life 05K19VH1, Code-Vita 05K16VH1, JW) and the German Center for Heart Diseases DZHK (JW, JG).

## Author contribution

Conception of the project: E.B., T.F., F.L., K.N., J.W.; Project management and supervision: E.B., T.F., F.L., J.W.; Sampling of wild fish: K.N.; Inbreeding: N.K., F.L., N.W.; Sample preparation: N.A., C.B., J.G., C.H., E.H., O.T.H., C.L., K.L., F.L., R.S., E.T., T.Ta., T.Th., P.W., B.W., J.W.; Data analysis: C.B., E.B., I.B., T.F., A.L., F.L., J.M., J.W.; Manuscript writing: E.B., I.B., T.F., A.L., F.L., J.W.

## Materials and Methods

### Fish husbandry and dissection

Medaka (*Oryzias latipes*) fish were maintained at the medaka facility of the Institute of Biological and Chemical Systems, Biological Information Processing (IBCS-BIP). Animal husbandry and experimental procedures were performed in accordance with local and European Union animal welfare standards (Tierschutzgesetz §11, Abs. 1, Nr. 1, AZ35-9185.64/BH). The facility is under the supervision of the local representative of the animal welfare agency.

The MIKK panel lines were established from a wild Medaka (*Oryzias latipes*) population as detailed in our back to back companion paper [4]. Liver and brain samples dissection procedures are also described in detail in the companion paper. In this paper we selected the following lines: **4–1**, **4–2**, **7–1**, **7–2**, 11-1, 69-1, 79-2, 80-1, 117-2, 131-1, **134–1** and **134–2**. Line ids starting with the same number (in bold) are sibling lines, derived from the same F1 founder family. The selection was done before full stabilisation of the final MIKK panel lines, leading to the following lines not being present in the official 80 stable MIKK lines: 7-1, 131-1 and 134-2.

### Sample preparation and sequencing

Briefly, RNA extraction from liver samples and DNA extraction from brain samples were performed on a Qiagen automated extraction platform using QIAsymphony RNA and DNA Kits, respectively. Samples were prepared for Illumina DNA-Sequencing using the standard PCR-free Illumina protocol and RNA-Sequencing using the NEBNext Ultra II Directional RNA Library Prep Kit following the manufacturer’s instructions.

For Nanopore DNA-Sequencing, Brain DNA samples were prepared with the ligation sequencing kit (SQK-LSK109), multiplexed with the native barcoding expansion kit (EXP-NBD104) and finally loaded in a FLO-PRO002 flow-cell on a PromethION instrument, all following the manufacturer’s instructions (Oxford Nanopore, Oxford, UK). To reduce sequencing costs while targeting a coverage of around 15X we multiplexed 4 samples per flowcell.

### Bioinformatic methods and data

Raw sequencing data can be retrieved from ENA linked to the following project ID:

- Nanopore DNA sequencing data: PRJEB43089
- Illumina DNA sequencing data: PRJEB17699
- Illumina RNA sequencing data: PRJEB43091

All the scripts and metadata used for this study are extensively described in the associated github repository available at https://github.com/birneylab/MIKK_genome_companion_paper.

### Nanopore data processing

#### Basecalling

After Nanopore sequencing, raw nanopore data in FAST5 format was transferred securely from Sanger Institute storage to the EBI high performance compute cluster, where all the analyses were performed. FAST5 files were basecalled and demultiplexed according to the 4 expected barcodes for each run with ONT-Guppy (v4.0.14). See detailed analysis and metadata at https://birneylab.github.io/MIKK_genome_companion_paper/Nanopore_basecalling/

#### Alignment and QC

We developed a Snakemake pipeline [37] called pycoSnake [38] to run the entire analysis, including mapping, quality control, differential methylation analysis and structural variation calling. For this study we ran pycoSnake v0.1a3 (commit hash 6d248c0fddfedd8f27d59b59f94f63f64d16e9bd), DNA_ONT workflow v0.2. All the tools and environment are version controlled in individual conda environments. Briefly, reference genome and annotations were obtained from ensembl Release 99 (Japanese medaka *HdrR* ASM223467v1, https://www.ensembl.org/Oryzias_latipes/Info/Index). Basecalled reads are merged and filtered using pyBiotools v0.2.0.9 [39], then aligned to the reference using Minimap2 v2.15 [40]. Alignments are filtered to keep only high quality primary reads using pyBiotools v0.2.0.9 [39], and quality control checks are performed using pycoQC v2.5.0.23 [41]. The detailed parameters used to run each tool as well as the sample QC can be found at https://birneylab.github.io/MIKK_genome_companion_paper/Nanopore_processing/.

#### DNA methylation analysis

The differential methylation analysis was performed as part of the pycoSnake pipeline, after the alignment steps described before. In brief, CpG methylated sites are called at single read level with nanopolish call_methylation v0.11.1 [26]. Methylation log likelihood ratio (LLR) are aggregated at genomic position level, then within CpG islands. Finally for each CpG island with sufficient coverage the differential methylation analysis is performed using pycoMeth v0.4.25 [32]. Briefly all median LLR values for each CpG positions within a given CpG island are compared between samples using a Kruskal Wallis test and all resulting p-values are adjusted for multiple tests using the Benjamini & Hochberg procedure for controlling the false discovery rate (FDR). We also performed extra analyses to produce the final paper figure in a Jupyter notebook. Additional information on the differential methylation analysis can be found at: https://birneylab.github.io/MIKK_genome_companion_paper/DNA_methylation/index

#### Structural variant calling

Structural variant calling was also performed as part of pycoSnake pipeline. Reads were re-aligned with NGMLR v0.2.7, followed by a first round of SV detection with Sniffles v1.0.12 [42]. Variants were subsequently filtered and merged with SURVIVOR v1.0.7 (https://github.com/fritzsedlazeck/SURVIVOR). Then a second round of Sniffles SV calling was done using the merge calls to constrain the detection to the common filtered variants previously collected. Finally, all calls are merged in a single unified VCF file. Additional information on the Structural variant calling analysis can be found at: https://birneylab.github.io/MIKK_genome_companion_paper/Nanopore_SV_analysis/. To polish the calls with Illumina reads, we used the Illumina reads and VCF described in [4] with SViper v2.0.0 (https://github.com/smehringer/SViper) to produce a “polished” set of structural variants for 9 of the 12 MIKK panel samples. We used bcftools v1.9 and Picard v2.25.0 [43,44] to further process the data, then R version 4.0.4 and a suite of R packages [45–55] to carry out the analysis set out in full at: https://brettellebi.github.io/mikk_genome/20210409_sv_notebook.html.

#### Prediction and annotation of repetitive and transposable elements

The *RepeatModeler* pipeline (v2.0.0) [56] for the automated *de novo* identification of repetitive and transposable elements was run on all chromosomes in the *HdrR* genome assembly [8]. RepeatModeler was run with its default parameters and the additional long terminal repeat (LTR) structural discovery sub-pipeline that includes the *LTRharvest [57]* and *LTR_retriever [58]* tools.

The RepeatModeler library of repeats was filtered to remove non-TE protein coding sequences by using a protein BLAST (Altschul *et al*., 1990) to align (*E-value* ≤ 1e-5) the *Oryzias latipes* proteome (Ensembl v99) and *pfam* peptide database (v32) against the RepeatMasker peptide library. Finally, a nucleotide BLAST was used to remove any RepeatModeler repeats that aligned (*E-value* ≤ 1e-10) against the corresponding transcripts.

RepeatMasker (v4.1.0) [59] was used to align the chromosomes in the *HdrR* assembly against the filtered RepeatModeler library of consensus repeats and the existing RepeatMasker repeat families.

Additionally, *Exonerate* (Slater and Birney, 2005) was used to align the two subtypes of the *Teratorn* mobile element found in the *Oryzias latipes* genome against the *HdrR* reference. (The *Teratorn* element being the result of a fusion between a *piggyBac* DNA transposon and a member of the *Alloherpesviridae* family [17]).

### Assembly and graph analysis

#### De novo assembly of MIKK panel genomes

For each line, Nanopore FASTQ raw sequences were assembled using the long-read assembler wtdbg2 in Nanopore (ONT) mode to create draft assemblies for the 12 MIKK panel genomes [28]. We then polished each of the draft assemblies with their corresponding ∼30X Illumina sequences using 2 rounds of the Pilon [29]. The draft assemblies quality were evaluated using QUAST v5.1.0rc1 [30] and FASTA were deposited at ENA under the same study accession as the nanopore reads (PRJEB43089). Additional information on the analysis and access to raw data can be obtained at https://birneylab.github.io/MIKK_genome_companion_paper/Individual_assemblies/.

#### Variation pangenome graph assembly

On top of the MIKK panel lines draft assemblies, we also used 3 high quality medaka reference assemblies *HdrR*, *HNI* and *HSOK*, including unanchored contigs, to scaffold the graph (http://utgenome.org/medaka_v2). Prior pangenome assembly each contig from every reference was prefixed with the reference name it belongs to, to allow unambiguous identification of the origin of graph segments (eg, HdrR_1 for chromosome 1 of the *HdrR* reference). We assembled the graph pangenome using minigraph2 v0.10 [11] (−x ggs mode) adding iteratively each reference in the following order HdrR, *HNI*, *HSOK*, then the MIKK lines 69-1, 131-1, 117-2, 4-2, 7-2, 4-1, 134-2, 79-2, 134-1, 80-1, 7-1 and finally 11-1. The resulting graph in [rGFA format](https://github.com/lh3/gfatools/blob/master/doc/rGFA.md#the-reference-gfa-rgfa-format) was parsed to extract descriptive statistics as well as graph anchored annotations for Bandage [60] and IGV [61] using python scripting. The analysis notebook and the raw data can be found at https://birneylab.github.io/MIKK_genome_companion_paper/Graph_assembly/.

#### Graph alignment and segment usage analysis

We aligned the DNA-Seq nanopore reads for each of our 12 MIKK samples to the pangenome graph using minigraph2 v0.10 [11] (−x lr mode) and obtained alignment files in [GAF format](https://github.com/lh3/gfatools/blob/master/doc/rGFA.md#the-graph-alignment-format-gaf). We also aligned the 50 Illumina RNA-Seq datasets obtained from MIKK lines liver samples described in [4] to the graph. However since pair-end mapping is not supported by minigraph, we first merged overlapping pairs together using Flash v1.2.11 [62], then aligned the merged reads to the graph with minigraph2 v0.10 [11] (−x sr mode). We then computed the length normalised coverage of segments and junctions between segments for each sample and generated statistics on graph segment usage per samples, using python scripting. The analysis notebook and the raw data can be found at https://birneylab.github.io/MIKK_genome_companion_paper/Graph_usage/.

#### Graph structural variation analysis

Based on the normalised coverage of graph segments and junctions we investigated the presence of 2 types of genetic variations in our MIKK panel: large scale divergent insertions with DNA and RNA-Seq supports and complex deletions. For the divergent insertions analysis we searched for alternative non-*HdrR* paths longer than 10kb, containing segments with a sequence diverging by more than 50%, supported by at least 2/12 samples for DNA-Seq (50% of mean *HdrR* coverage) and 8/50 samples for RNA-Seq (10% of mean *HdrR* coverage) and overlapping at least 1 annotated gene exon. With this very strict set of criteria we found a set of 19 such paths (Table SX). For the complex deletions, we leveraged the coverage information for the junction/link between segments instead. We identified 16 deletions, supported by junctions connecting 2 *HdrR* segments distant by more than 10kb, 2) with a coverage greater than 50% of the average *HdrR* supported by at least half of the panel lines3) skipping at least 1 full annotated *HdrR* exon. These candidate insertions and deletions were then manually investigated using Bandage [60] for visualisation in graph space and IGV [61] for HdrR anchored linear genome visualisation. The jupyter notebook containing the full analysis and raw data can be found at https://birneylab.github.io/MIKK_genome_companion_paper/Graph_SV/.

