## Supplementary figures for "Genomic variations and epigenomic landscape of the Medaka Inbred Kiyosu-Karlsruhe (MIKK) panel"

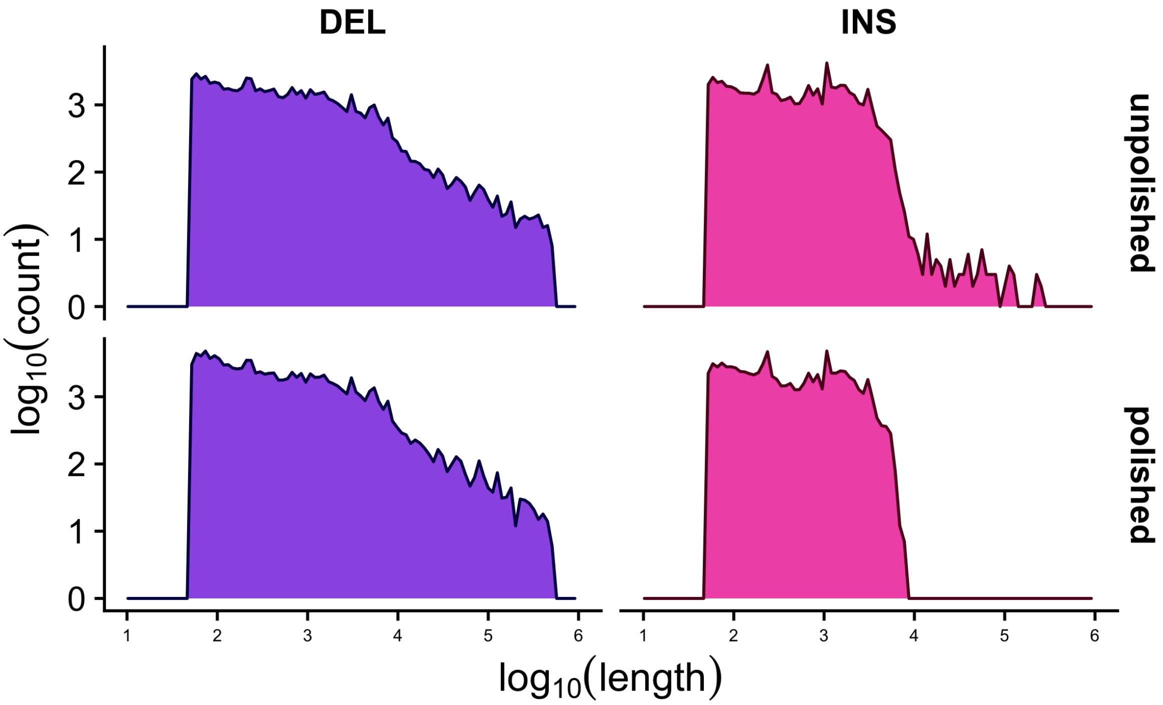


**Supplementary Figure 1**: *$log_{10}$ lengths and counts of deletion (DEL) and insertion (INS) structural variants before and after polishing with SViper.*

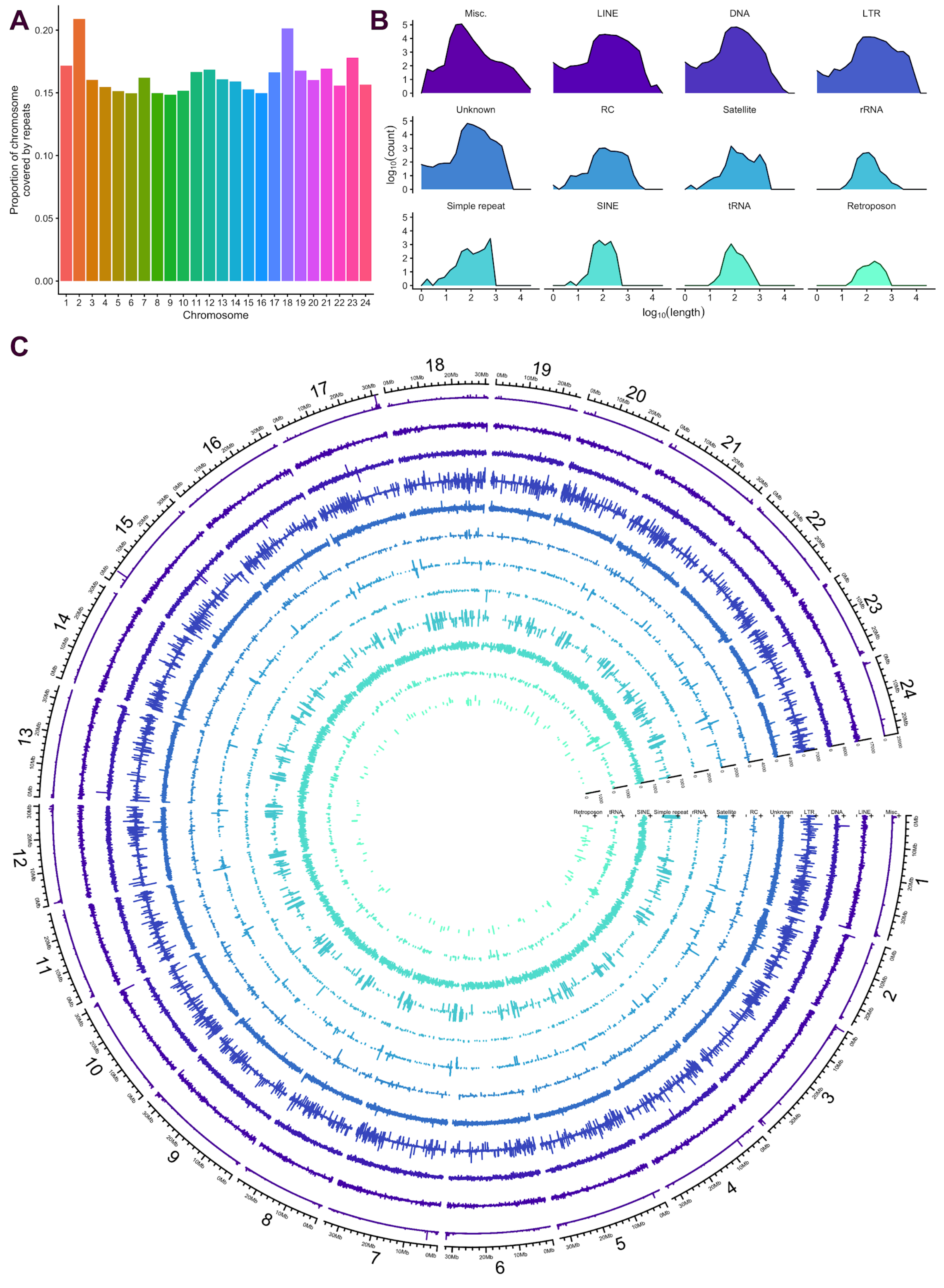


**Supplementary Figure 2**: *Repeat content in the* HdrR *genome based on RepeatMasker results (**Methods**). **A**. Proportion of repeat content per-chromosome. **B**. $log_{10}$ of repeat lengths and counts per repeat class. "Misc" includes all repeats assigned to their own specific class, for example "(GAG)n" or "(GATCCA)n". **C**. Circos plot showing repeat length (radial axes) by locus (angular axis) and repeat class (track). The code and methods used to generate the figure are set out here:*
https://birneylab.github.io/MIKK_genome_main_paper/20210409_repeats.html
