## Supplementary file 1 for "Genomic variations and epigenomic landscape of the Medaka Inbred Kiyosu-Karlsruhe (MIKK) panel"

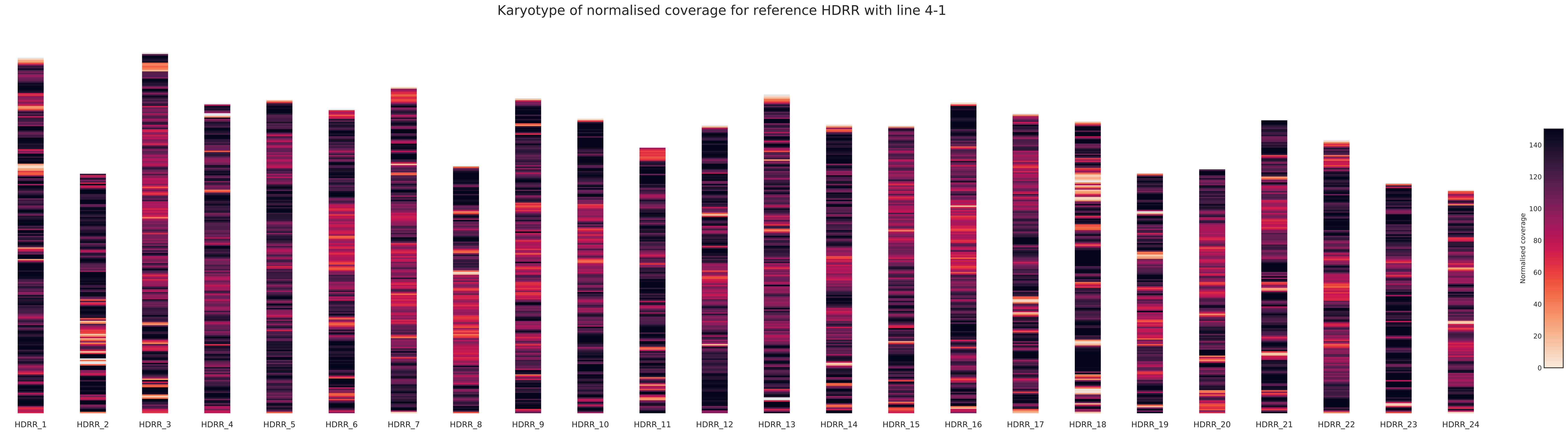

Karyotype of normalised coverage for reference HDRR with line 4-1
HDRR_1
HDRR_2
HDRR_3
HDRR_4
HDRR_5
HDRR_6
HDRR_7
HDRR_8
HDRR_9
HDRR_10
HDRR_11
HDRR_12
HDRR_13
HDRR_14
HDRR_15
HDRR_16
HDRR_17
HDRR_18
HDRR_19
HDRR_20
HDRR_21
HDRR_22
HDRR_23
HDRR_24
Normalised coverage
140
120
100
80
60
40
20
0

Karyotype of normalised coverage for reference HDRR with line 4-2

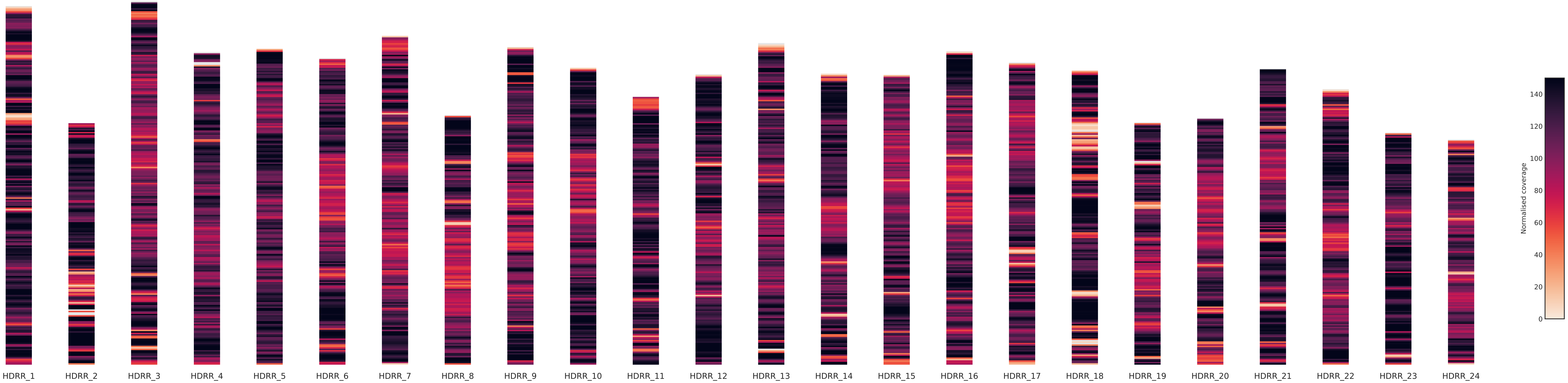

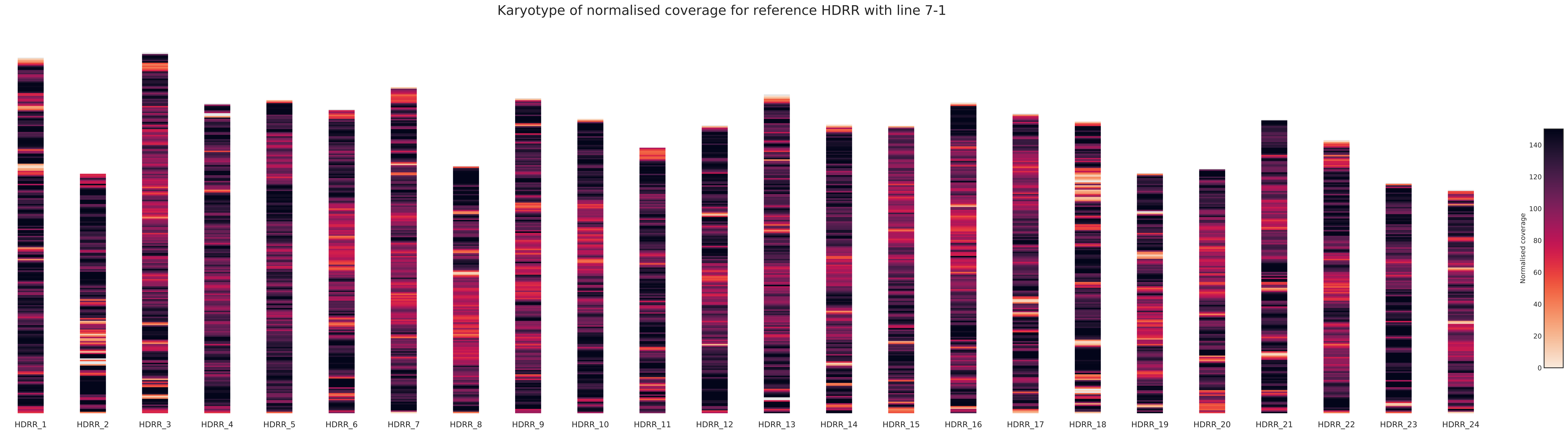
Karyotype of normalised coverage for reference HDRR with line 7-1
HDRR_1
HDRR_2
HDRR_3
HDRR_4
HDRR_5
HDRR_6
HDRR_7
HDRR_8
HDRR_9
HDRR_10
HDRR_11
HDRR_12
HDRR_13
HDRR_14
HDRR_15
HDRR_16
HDRR_17
HDRR_18
HDRR_19
HDRR_20
HDRR_21
HDRR_22
HDRR_23
HDRR_24
Normalised coverage
0
20
40
60
80
100
120
140

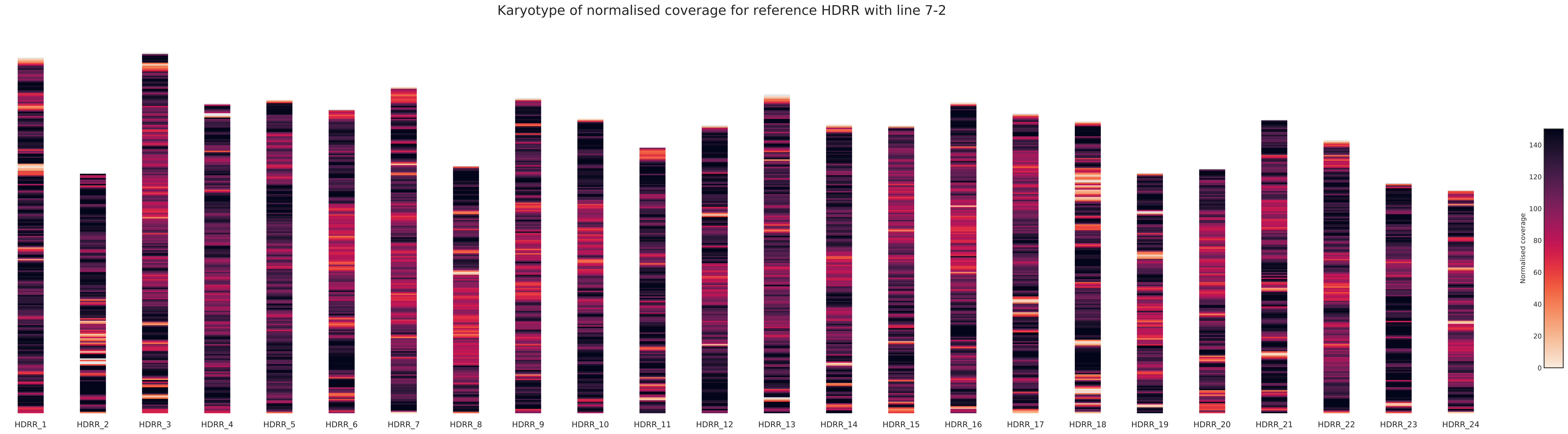
Karyotype of normalised coverage for reference HDRR with line 7-2
HDRR_1
HDRR_2
HDRR_3
HDRR_4
HDRR_5
HDRR_6
HDRR_7
HDRR_8
HDRR_9
HDRR_10
HDRR_11
HDRR_12
HDRR_13
HDRR_14
HDRR_15
HDRR_16
HDRR_17
HDRR_18
HDRR_19
HDRR_20
HDRR_21
HDRR_22
HDRR_23
HDRR_24
Normalised coverage
0
20
40
60
80
100
120
140

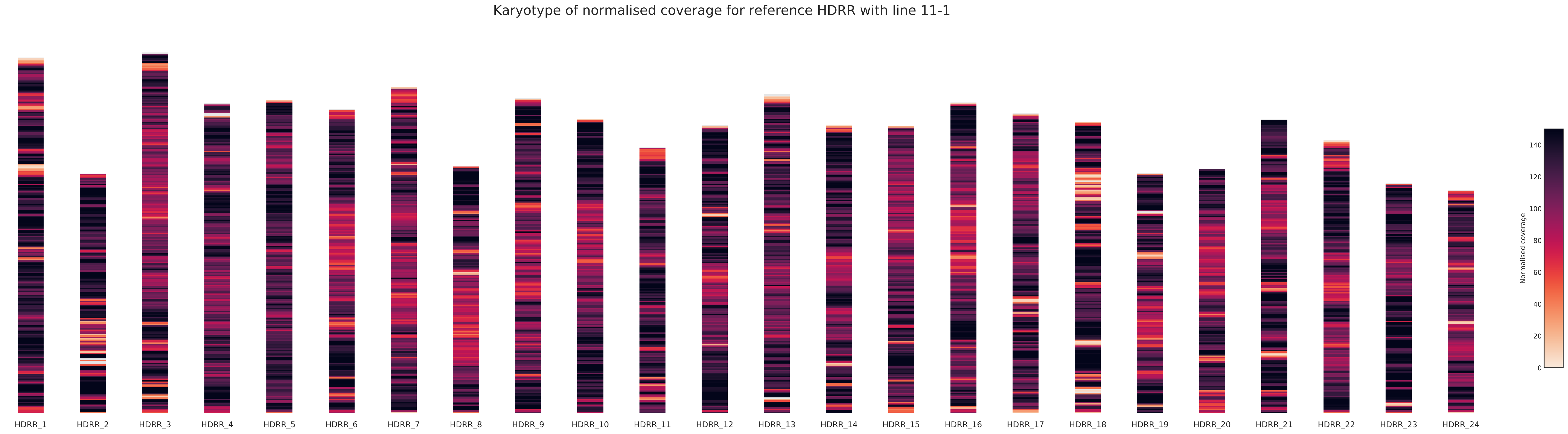
Karyotype of normalised coverage for reference HDRR with line 11-1
HDRR_1
HDRR_2
HDRR_3
HDRR_4
HDRR_5
HDRR_6
HDRR_7
HDRR_8
HDRR_9
HDRR_10
HDRR_11
HDRR_12
HDRR_13
HDRR_14
HDRR_15
HDRR_16
HDRR_17
HDRR_18
HDRR_19
HDRR_20
HDRR_21
HDRR_22
HDRR_23
HDRR_24
Normalised coverage
0
20
40
60
80
100
120
140

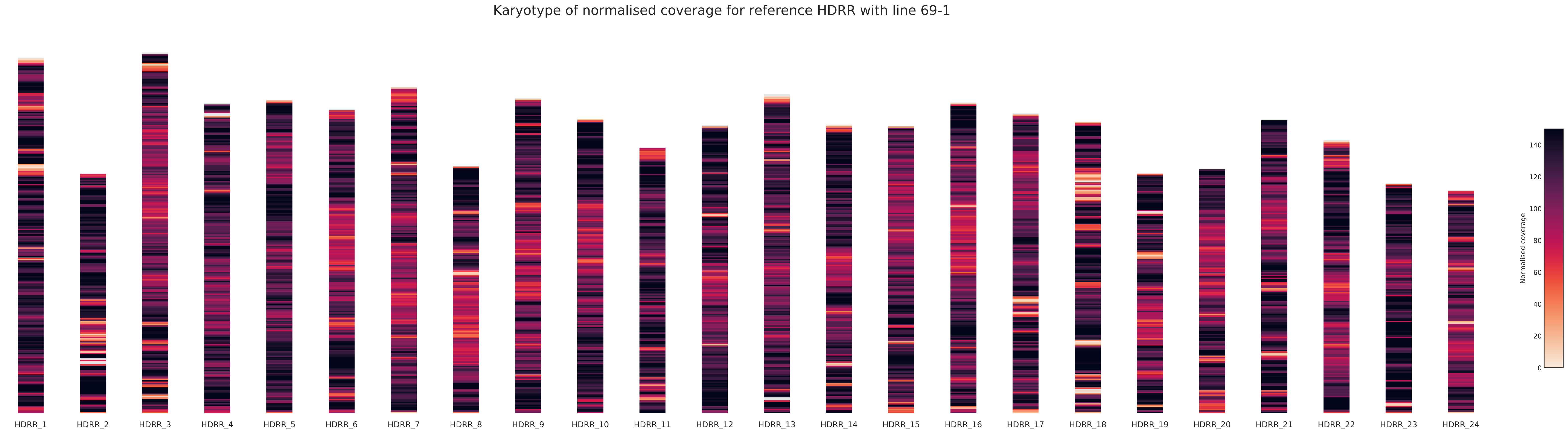

Karyotype of normalised coverage for reference HDRR with line 69-1
HDRR_1
HDRR_2
HDRR_3
HDRR_4
HDRR_5
HDRR_6
HDRR_7
HDRR_8
HDRR_9
HDRR_10
HDRR_11
HDRR_12
HDRR_13
HDRR_14
HDRR_15
HDRR_16
HDRR_17
HDRR_18
HDRR_19
HDRR_20
HDRR_21
HDRR_22
HDRR_23
HDRR_24
Normalised coverage
0
20
40
60
80
100
120
140

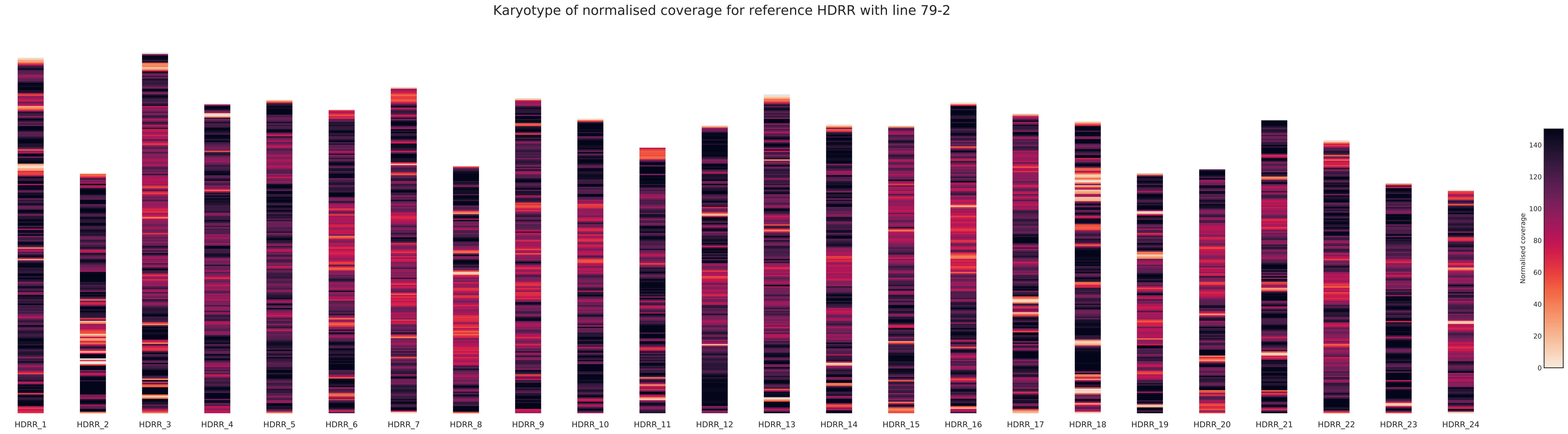
Karyotype of normalised coverage for reference HDRR with line 79-2
HDRR_1
HDRR_2
HDRR_3
HDRR_4
HDRR_5
HDRR_6
HDRR_7
HDRR_8
HDRR_9
HDRR_10
HDRR_11
HDRR_12
HDRR_13
HDRR_14
HDRR_15
HDRR_16
HDRR_17
HDRR_18
HDRR_19
HDRR_20
HDRR_21
HDRR_22
HDRR_23
HDRR_24
Normalised coverage
0
20
40
60
80
100
120
140

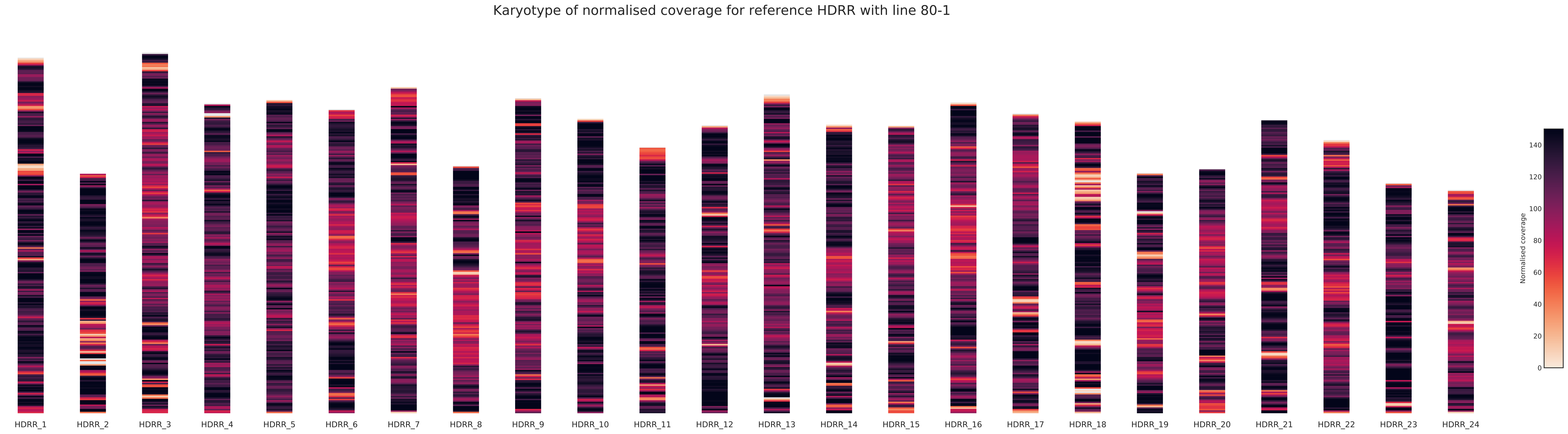
Karyotype of normalised coverage for reference HDRR with line 80-1
HDRR_1
HDRR_2
HDRR_3
HDRR_4
HDRR_5
HDRR_6
HDRR_7
HDRR_8
HDRR_9
HDRR_10
HDRR_11
HDRR_12
HDRR_13
HDRR_14
HDRR_15
HDRR_16
HDRR_17
HDRR_18
HDRR_19
HDRR_20
HDRR_21
HDRR_22
HDRR_23
HDRR_24
Normalised coverage
0
20
40
60
80
100
120
140

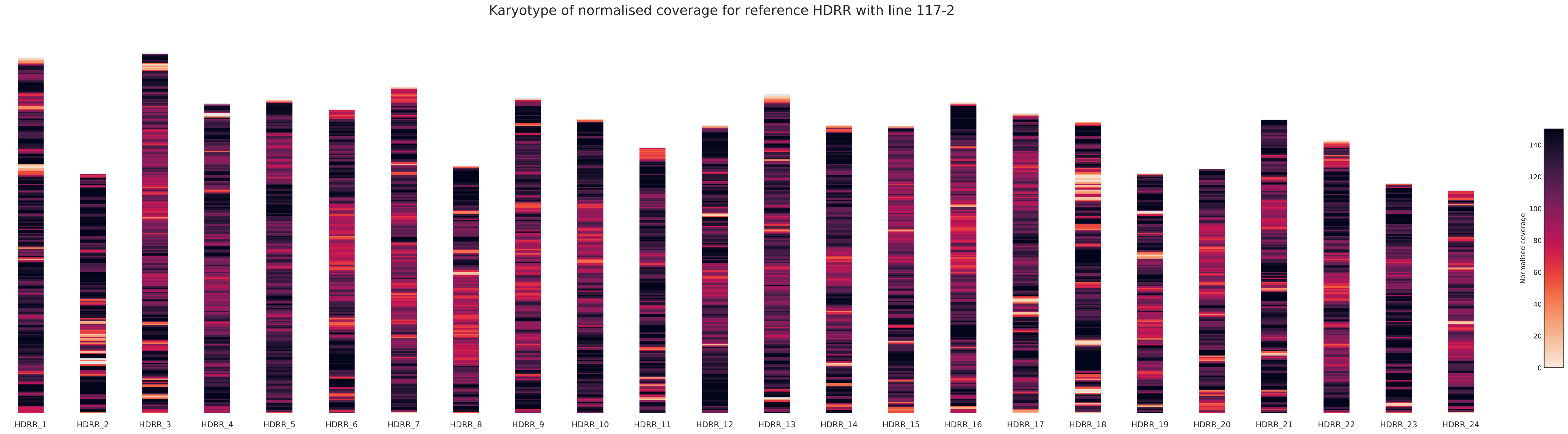
Karyotype of normalised coverage for reference HDRR with line 117-2
HDRR_1
HDRR_2
HDRR_3
HDRR_4
HDRR_5
HDRR_6
HDRR_7
HDRR_8
HDRR_9
HDRR_10
HDRR_11
HDRR_12
HDRR_13
HDRR_14
HDRR_15
HDRR_16
HDRR_17
HDRR_18
HDRR_19
HDRR_20
HDRR_21
HDRR_22
HDRR_23
HDRR_24
Normalised coverage
0
20
40
60
80
100
120
140

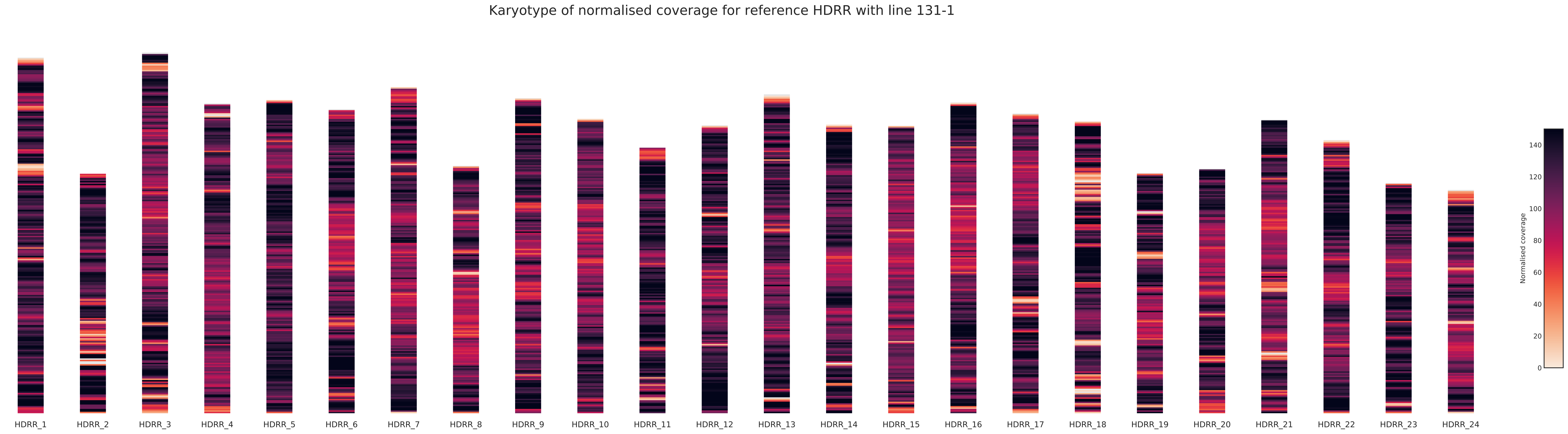
Karyotype of normalised coverage for reference HDRR with line 131-1
HDRR_1
HDRR_2
HDRR_3
HDRR_4
HDRR_5
HDRR_6
HDRR_7
HDRR_8
HDRR_9
HDRR_10
HDRR_11
HDRR_12
HDRR_13
HDRR_14
HDRR_15
HDRR_16
HDRR_17
HDRR_18
HDRR_19
HDRR_20
HDRR_21
HDRR_22
HDRR_23
HDRR_24
Normalised coverage
0
20
40
60
80
100
120
140

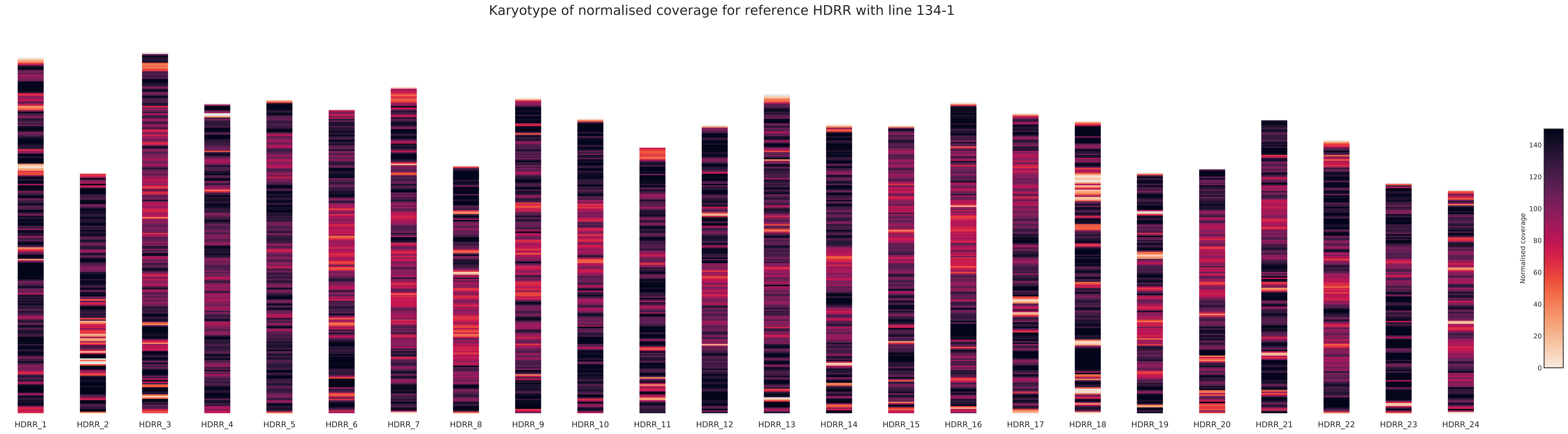
Karyotype of normalised coverage for reference HDRR with line 134-1
HDRR_1
HDRR_2
HDRR_3
HDRR_4
HDRR_5
HDRR_6
HDRR_7
HDRR_8
HDRR_9
HDRR_10
HDRR_11
HDRR_12
HDRR_13
HDRR_14
HDRR_15
HDRR_16
HDRR_17
HDRR_18
HDRR_19
HDRR_20
HDRR_21
HDRR_22
HDRR_23
HDRR_24
Normalised coverage
0
20
40
60
80
100
120
140

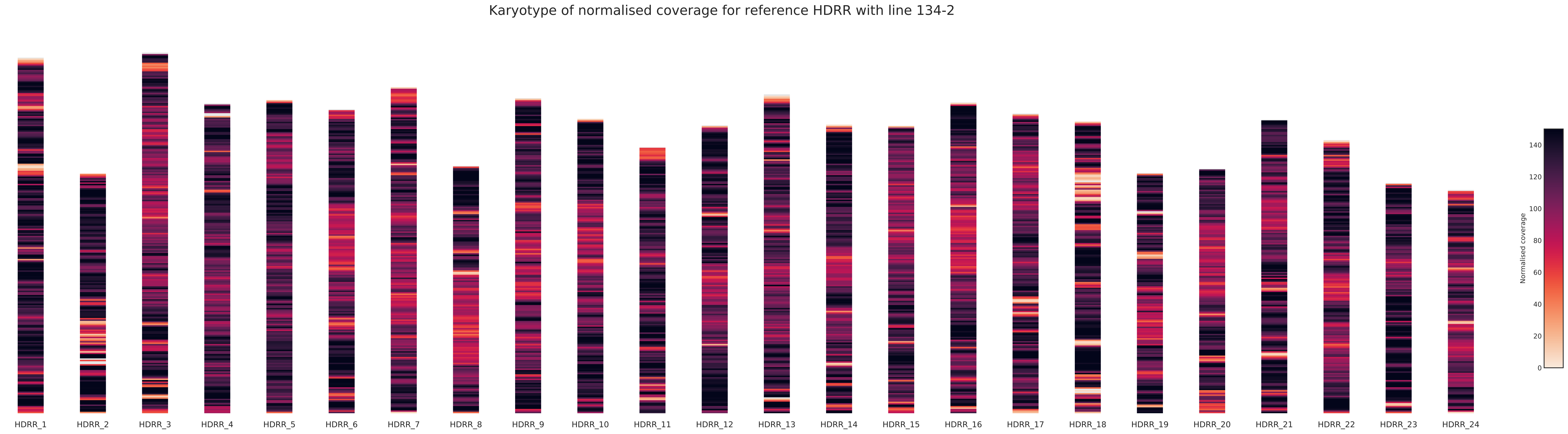

Karyotype of normalised coverage for reference HDRR with line 134-2
HDRR_1
HDRR_2
HDRR_3
HDRR_4
HDRR_5
HDRR_6
HDRR_7
HDRR_8
HDRR_9
HDRR_10
HDRR_11
HDRR_12
HDRR_13
HDRR_14
HDRR_15
HDRR_16
HDRR_17
HDRR_18
HDRR_19
HDRR_20
HDRR_21
HDRR_22
HDRR_23
HDRR_24
Normalised coverage
0
20
40
60
80
100
120
140

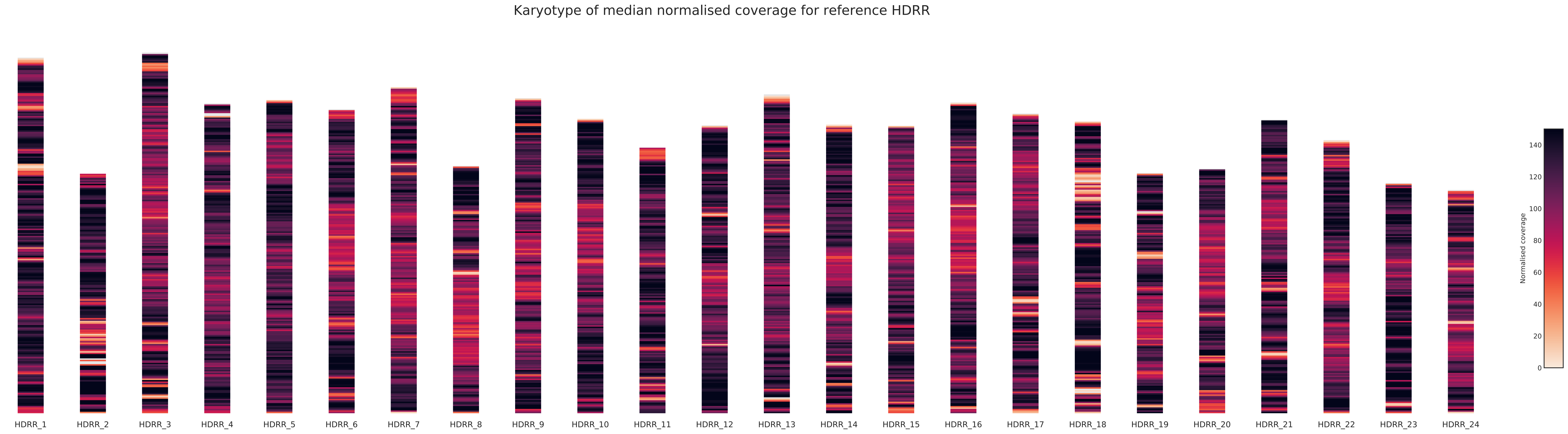
Karyotype of median normalised coverage for reference HDRR
HDRR_1
HDRR_2
HDRR_3
HDRR_4
HDRR_5
HDRR_6
HDRR_7
HDRR_8
HDRR_9
HDRR_10
HDRR_11
HDRR_12
HDRR_13
HDRR_14
HDRR_15
HDRR_16
HDRR_17
HDRR_18
HDRR_19
HDRR_20
HDRR_21
HDRR_22
HDRR_23
HDRR_24
Normalised coverage
0
20
40
60
80
100
120
140

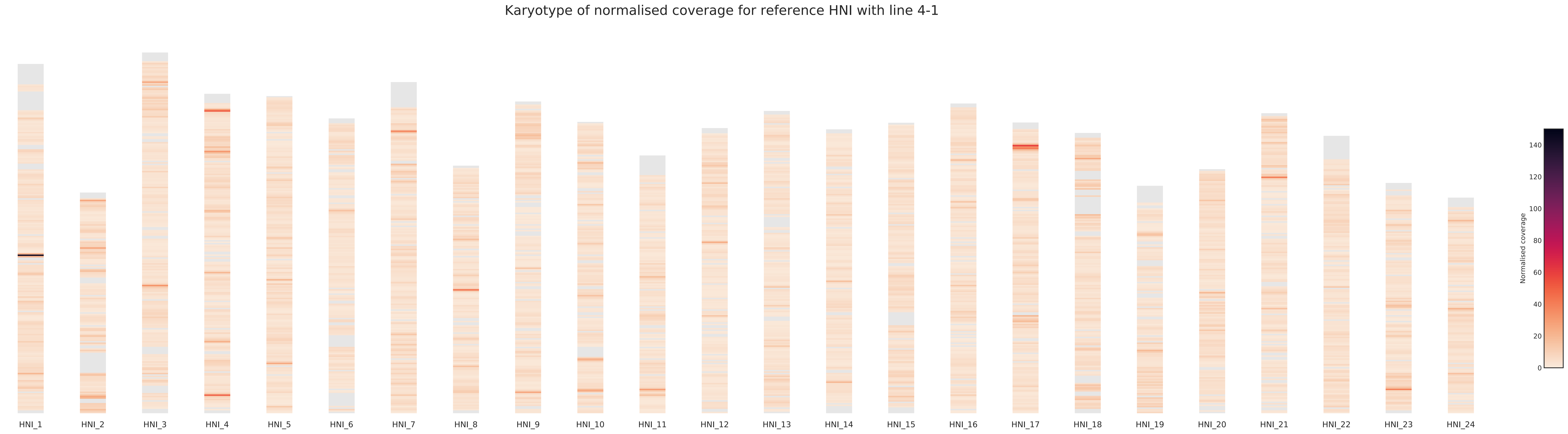

Karyotype of normalised coverage for reference HNI with line 4-1
HNI_1
HNI_2
HNI_3
HNI_4
HNI_5
HNI_6
HNI_7
HNI_8
HNI_9
HNI_10
HNI_11
HNI_12
HNI_13
HNI_14
HNI_15
HNI_16
HNI_17
HNI_18
HNI_19
HNI_20
HNI_21
HNI_22
HNI_23
HNI_24
Normalised coverage
0
20
40
60
80
100
120
140

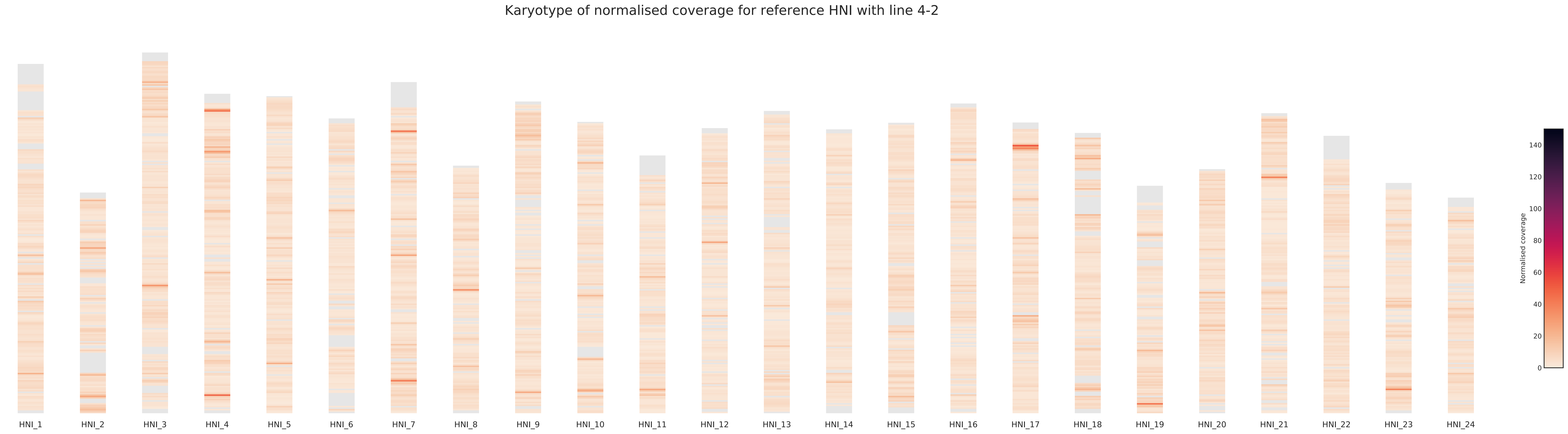
Karyotype of normalised coverage for reference HNI with line 4-2
HNI_1
HNI_2
HNI_3
HNI_4
HNI_5
HNI_6
HNI_7
HNI_8
HNI_9
HNI_10
HNI_11
HNI_12
HNI_13
HNI_14
HNI_15
HNI_16
HNI_17
HNI_18
HNI_19
HNI_20
HNI_21
HNI_22
HNI_23
HNI_24
Normalised coverage
0
20
40
60
80
100
120
140

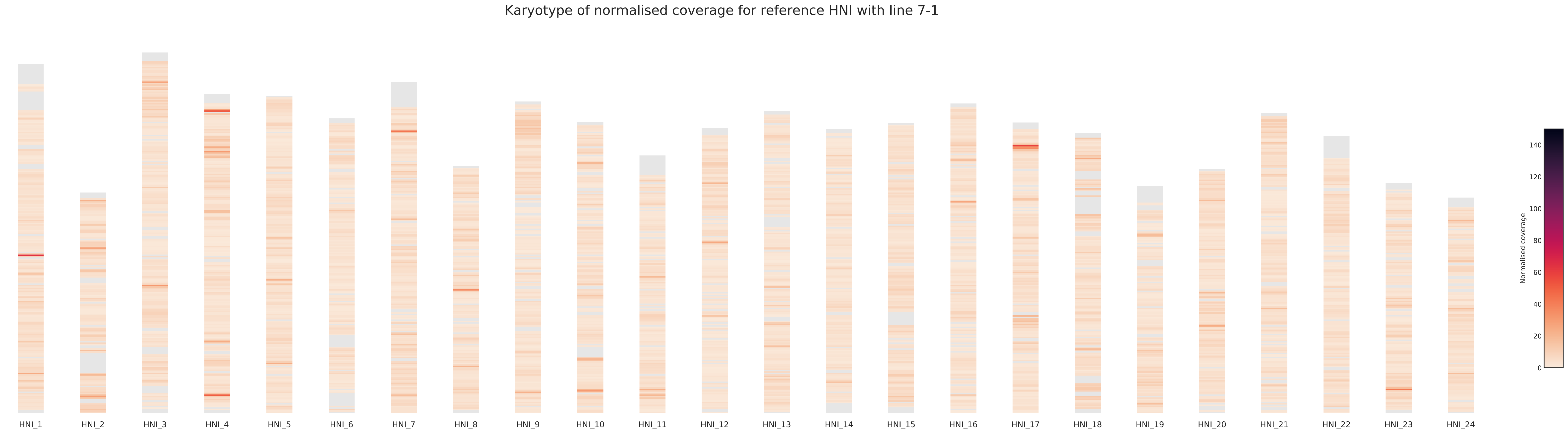
Karyotype of normalised coverage for reference HNI with line 7-1
HNI_1
HNI_2
HNI_3
HNI_4
HNI_5
HNI_6
HNI_7
HNI_8
HNI_9
HNI_10
HNI_11
HNI_12
HNI_13
HNI_14
HNI_15
HNI_16
HNI_17
HNI_18
HNI_19
HNI_20
HNI_21
HNI_22
HNI_23
HNI_24
Normalised coverage
0
20
40
60
80
100
120
140

Karyotype of normalised coverage for reference HNI with line 7-2

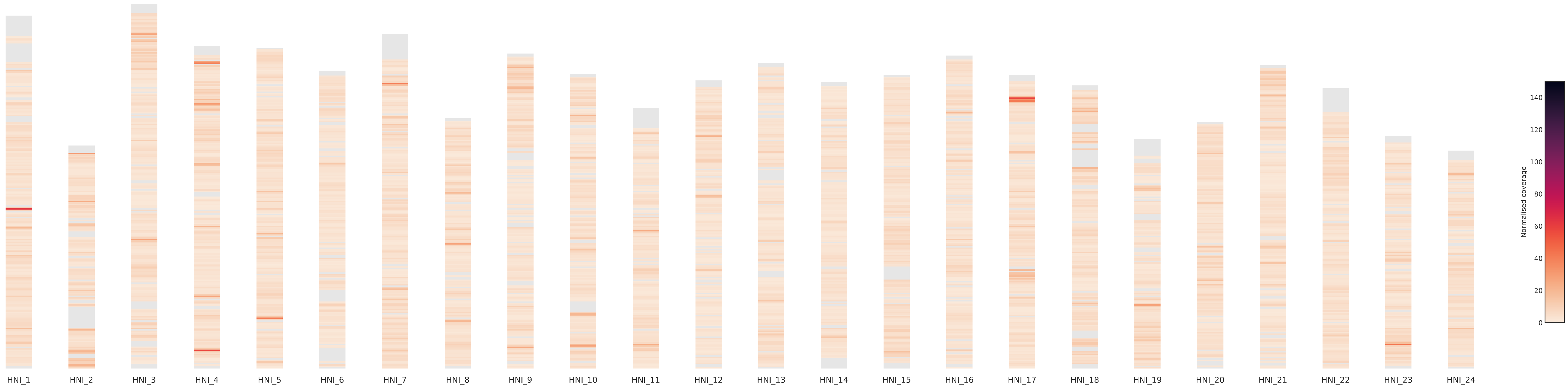

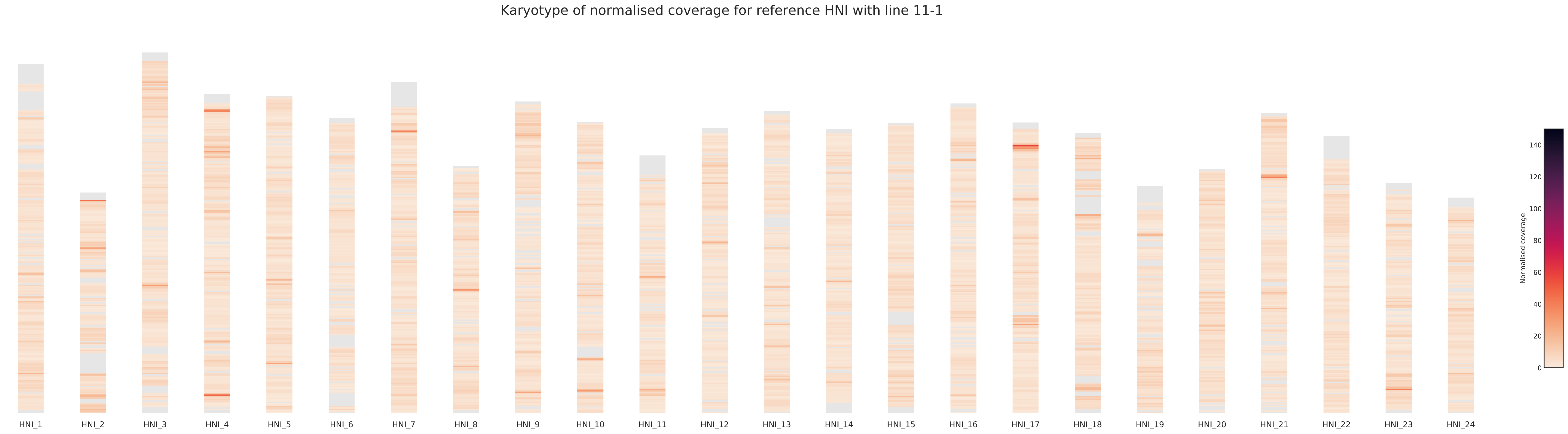
Karyotype of normalised coverage for reference HNI with line 11-1
HNI_1
HNI_2
HNI_3
HNI_4
HNI_5
HNI_6
HNI_7
HNI_8
HNI_9
HNI_10
HNI_11
HNI_12
HNI_13
HNI_14
HNI_15
HNI_16
HNI_17
HNI_18
HNI_19
HNI_20
HNI_21
HNI_22
HNI_23
HNI_24
Normalised coverage
0
20
40
60
80
100
120
140

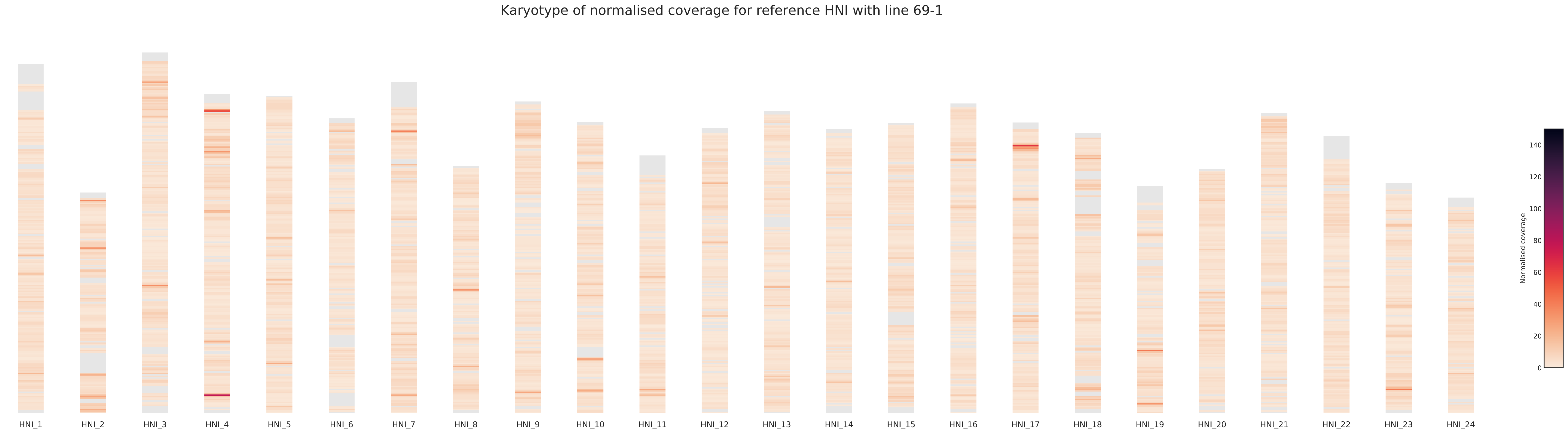

Karyotype of normalised coverage for reference HNI with line 69-1
HNI_1
HNI_2
HNI_3
HNI_4
HNI_5
HNI_6
HNI_7
HNI_8
HNI_9
HNI_10
HNI_11
HNI_12
HNI_13
HNI_14
HNI_15
HNI_16
HNI_17
HNI_18
HNI_19
HNI_20
HNI_21
HNI_22
HNI_23
HNI_24
Normalised coverage
0
20
40
60
80
100
120
140

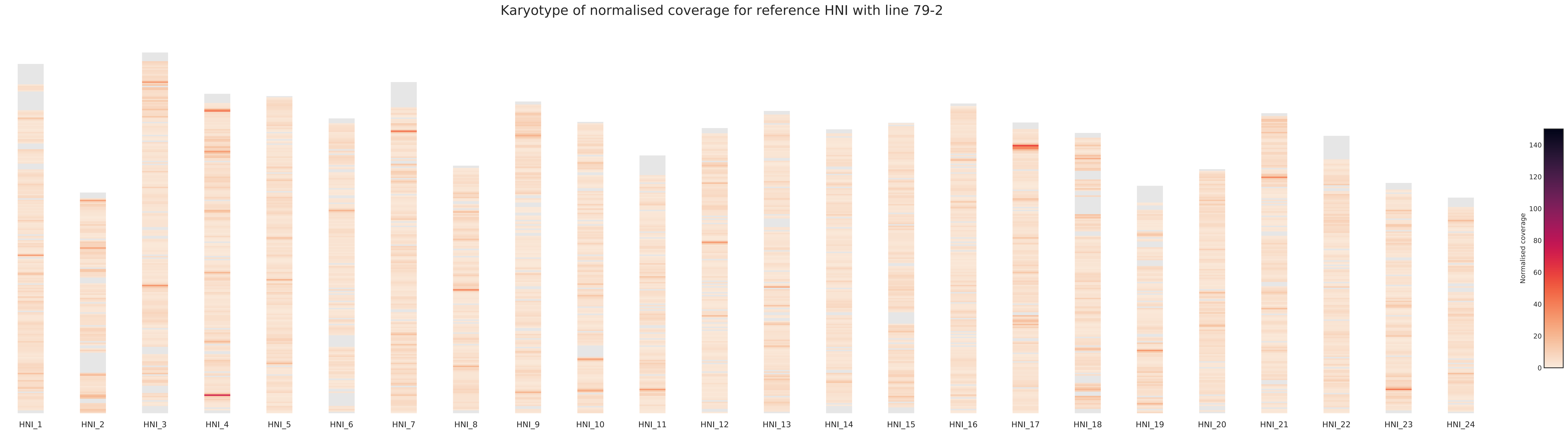

Karyotype of normalised coverage for reference HNI with line 79-2
HNI_1
HNI_2
HNI_3
HNI_4
HNI_5
HNI_6
HNI_7
HNI_8
HNI_9
HNI_10
HNI_11
HNI_12
HNI_13
HNI_14
HNI_15
HNI_16
HNI_17
HNI_18
HNI_19
HNI_20
HNI_21
HNI_22
HNI_23
HNI_24
Normalised coverage
0
20
40
60
80
100
120
140

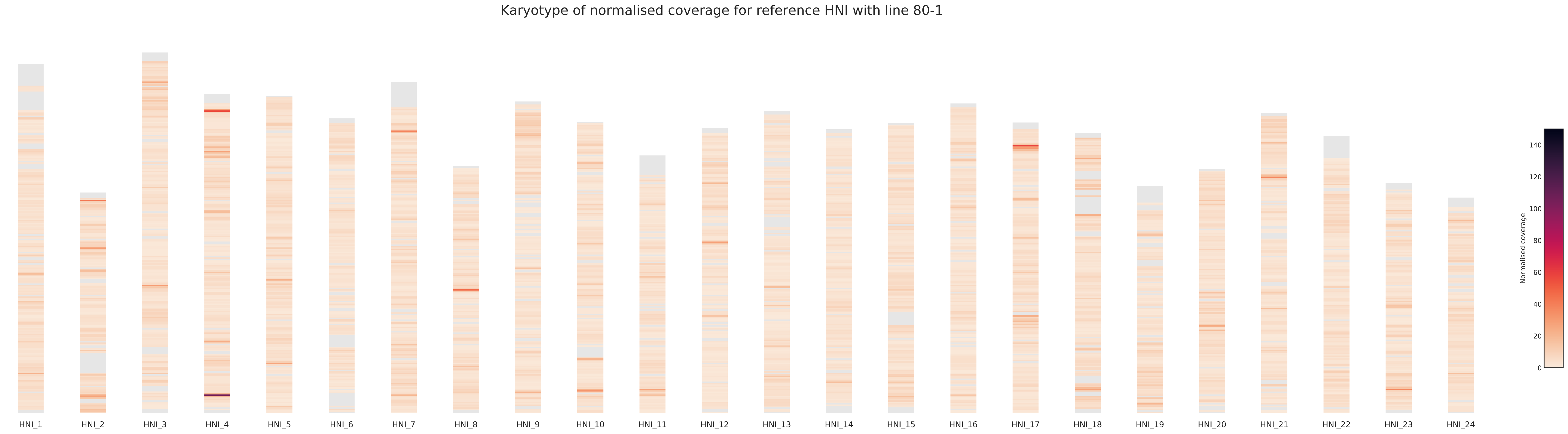
Karyotype of normalised coverage for reference HNI with line 80-1
HNI_1
HNI_2
HNI_3
HNI_4
HNI_5
HNI_6
HNI_7
HNI_8
HNI_9
HNI_10
HNI_11
HNI_12
HNI_13
HNI_14
HNI_15
HNI_16
HNI_17
HNI_18
HNI_19
HNI_20
HNI_21
HNI_22
HNI_23
HNI_24
Normalised coverage
0
20
40
60
80
100
120
140

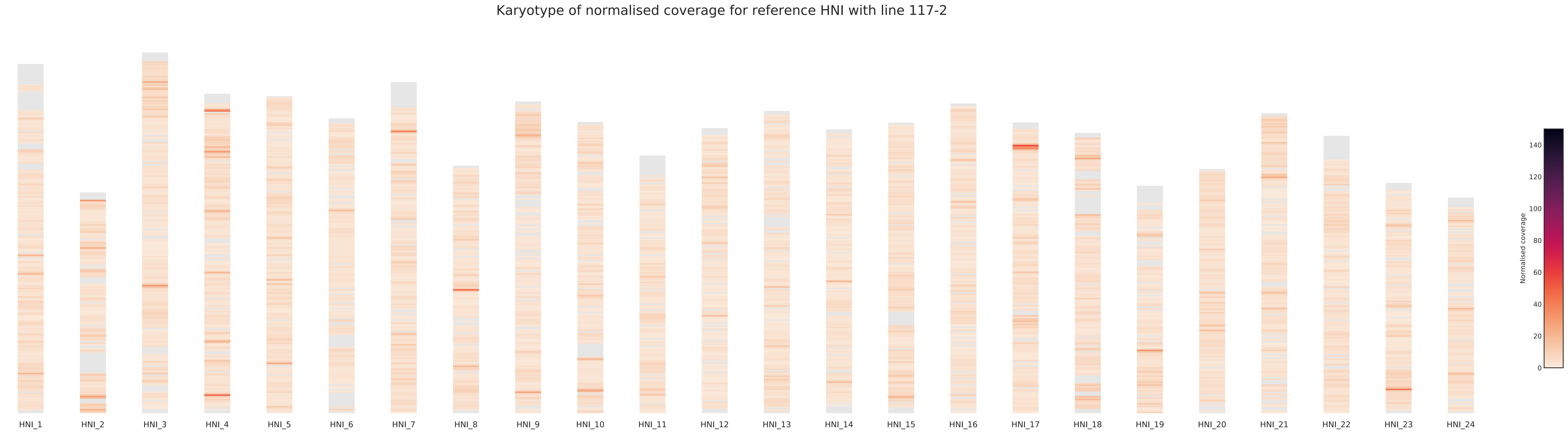
Karyotype of normalised coverage for reference HNI with line 117-2
HNI_1
HNI_2
HNI_3
HNI_4
HNI_5
HNI_6
HNI_7
HNI_8
HNI_9
HNI_10
HNI_11
HNI_12
HNI_13
HNI_14
HNI_15
HNI_16
HNI_17
HNI_18
HNI_19
HNI_20
HNI_21
HNI_22
HNI_23
HNI_24
Normalised coverage
0
20
40
60
80
100
120
140

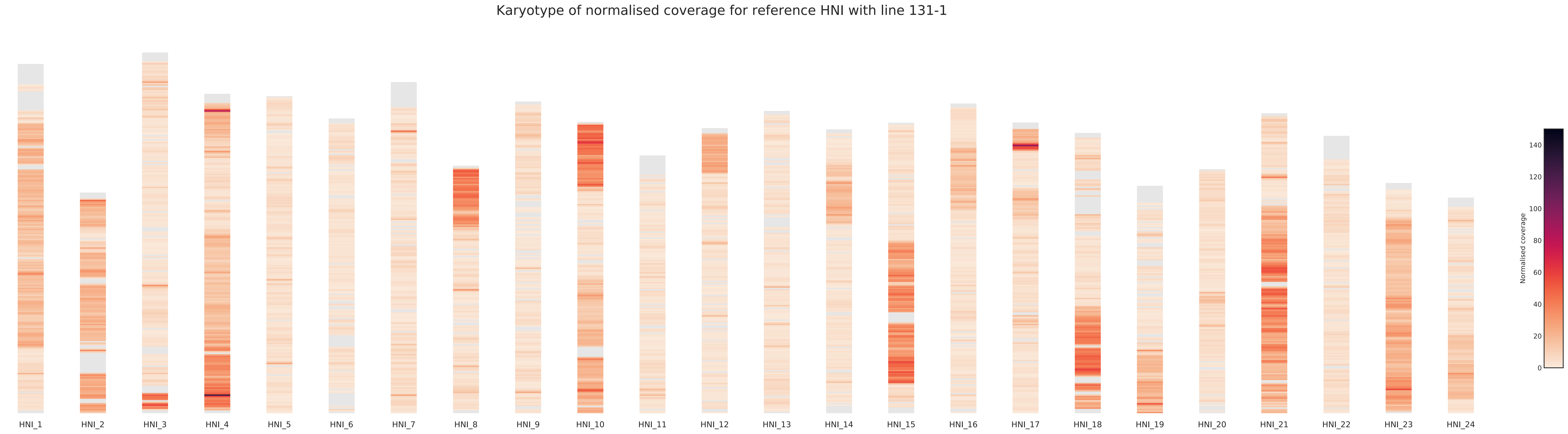
Karyotype of normalised coverage for reference HNI with line 131-1
HNI_1
HNI_2
HNI_3
HNI_4
HNI_5
HNI_6
HNI_7
HNI_8
HNI_9
HNI_10
HNI_11
HNI_12
HNI_13
HNI_14
HNI_15
HNI_16
HNI_17
HNI_18
HNI_19
HNI_20
HNI_21
HNI_22
HNI_23
HNI_24
Normalised coverage
0
20
40
60
80
100
120
140

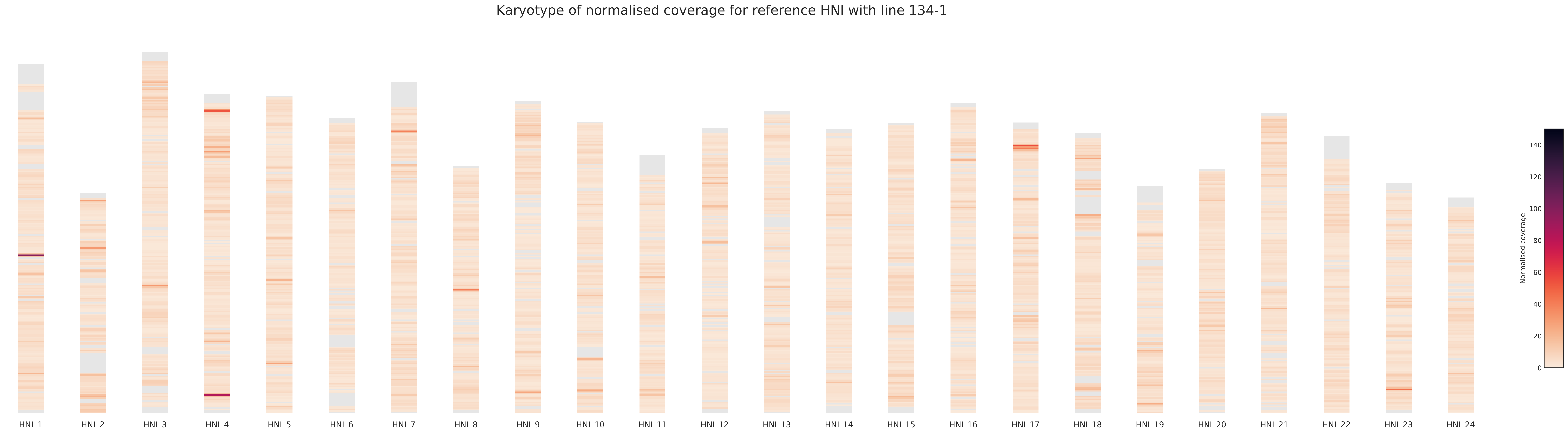
Karyotype of normalised coverage for reference HNI with line 134-1
HNI_1
HNI_2
HNI_3
HNI_4
HNI_5
HNI_6
HNI_7
HNI_8
HNI_9
HNI_10
HNI_11
HNI_12
HNI_13
HNI_14
HNI_15
HNI_16
HNI_17
HNI_18
HNI_19
HNI_20
HNI_21
HNI_22
HNI_23
HNI_24
Normalised coverage
0
20
40
60
80
100
120
140

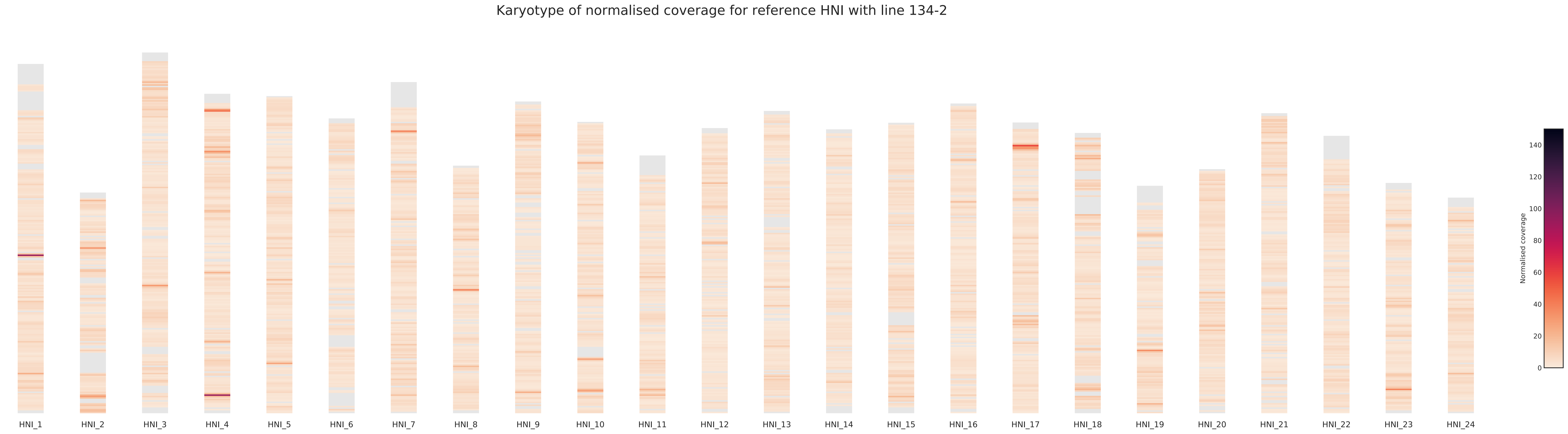
Karyotype of normalised coverage for reference HNI with line 134-2
HNI_1
HNI_2
HNI_3
HNI_4
HNI_5
HNI_6
HNI_7
HNI_8
HNI_9
HNI_10
HNI_11
HNI_12
HNI_13
HNI_14
HNI_15
HNI_16
HNI_17
HNI_18
HNI_19
HNI_20
HNI_21
HNI_22
HNI_23
HNI_24
Normalised coverage
0
20
40
60
80
100
120
140

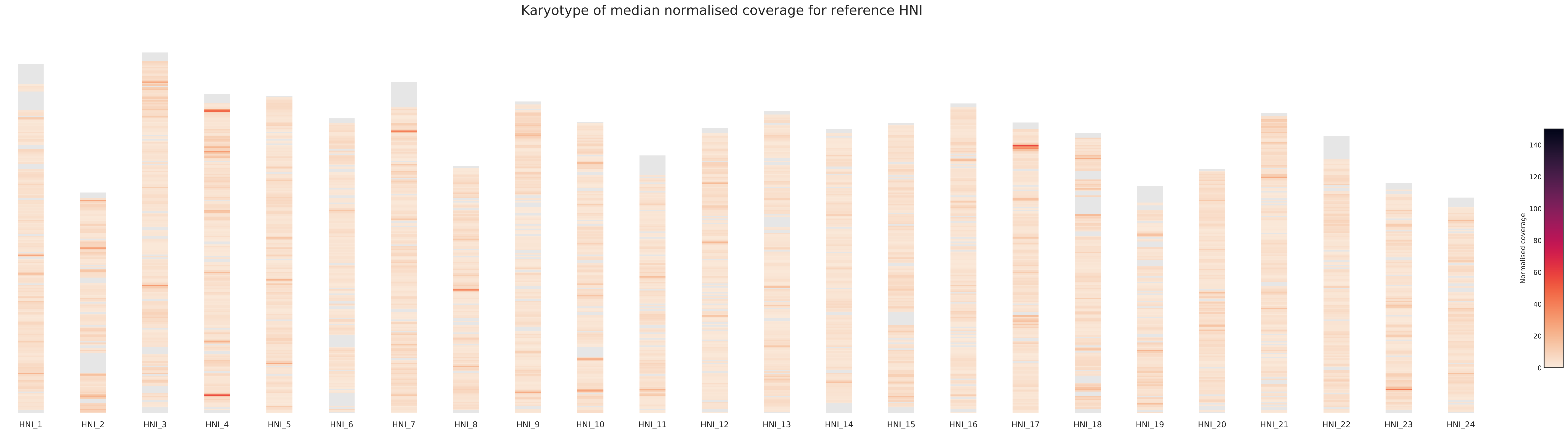

Karyotype of median normalised coverage for reference HNI
HNI_1
HNI_2
HNI_3
HNI_4
HNI_5
HNI_6
HNI_7
HNI_8
HNI_9
HNI_10
HNI_11
HNI_12
HNI_13
HNI_14
HNI_15
HNI_16
HNI_17
HNI_18
HNI_19
HNI_20
HNI_21
HNI_22
HNI_23
HNI_24
Normalised coverage
0
20
40
60
80
100
120
140

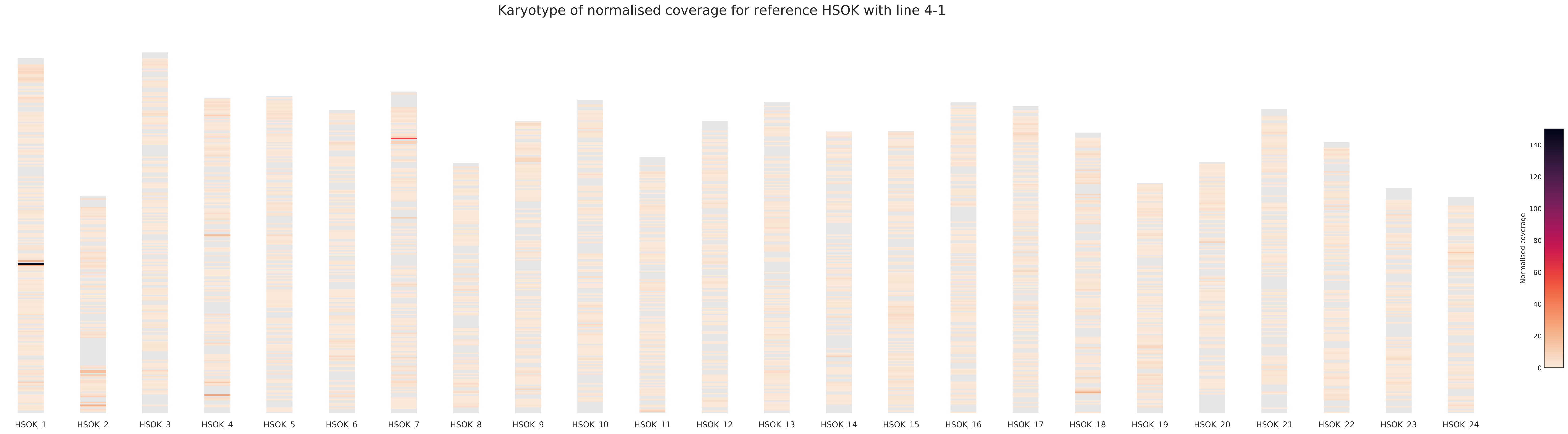
Karyotype of normalised coverage for reference HSOK with line 4-1
HSOK_1
HSOK_2
HSOK_3
HSOK_4
HSOK_5
HSOK_6
HSOK_7
HSOK_8
HSOK_9
HSOK_10
HSOK_11
HSOK_12
HSOK_13
HSOK_14
HSOK_15
HSOK_16
HSOK_17
HSOK_18
HSOK_19
HSOK_20
HSOK_21
HSOK_22
HSOK_23
HSOK_24
Normalised coverage
0
20
40
60
80
100
120
140

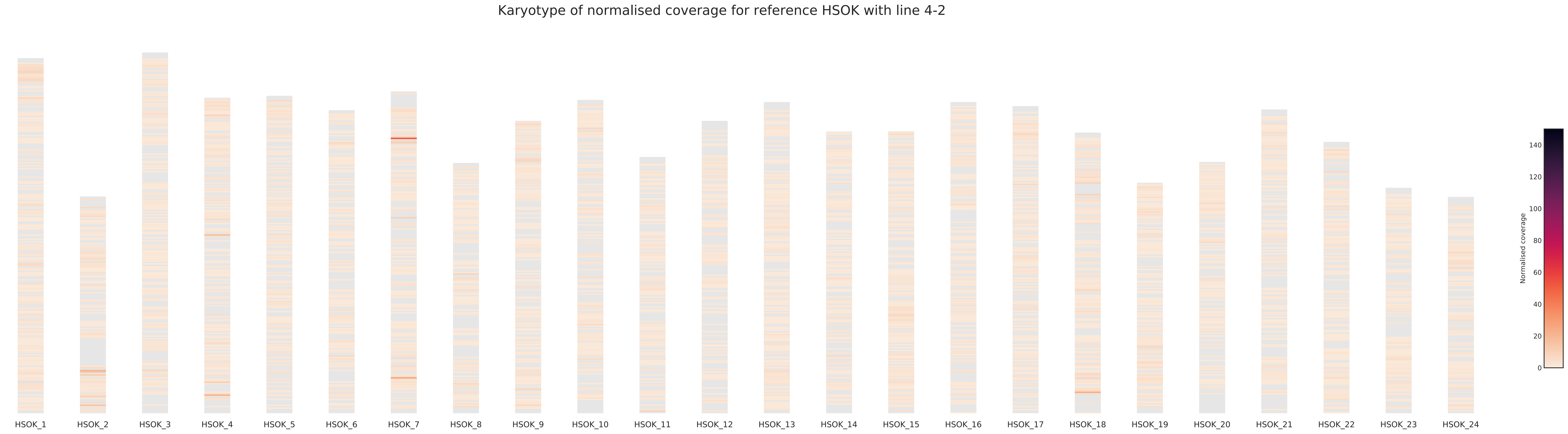
Karyotype of normalised coverage for reference HSOK with line 4-2
HSOK_1
HSOK_2
HSOK_3
HSOK_4
HSOK_5
HSOK_6
HSOK_7
HSOK_8
HSOK_9
HSOK_10
HSOK_11
HSOK_12
HSOK_13
HSOK_14
HSOK_15
HSOK_16
HSOK_17
HSOK_18
HSOK_19
HSOK_20
HSOK_21
HSOK_22
HSOK_23
HSOK_24
Normalised coverage
0
20
40
60
80
100
120
140

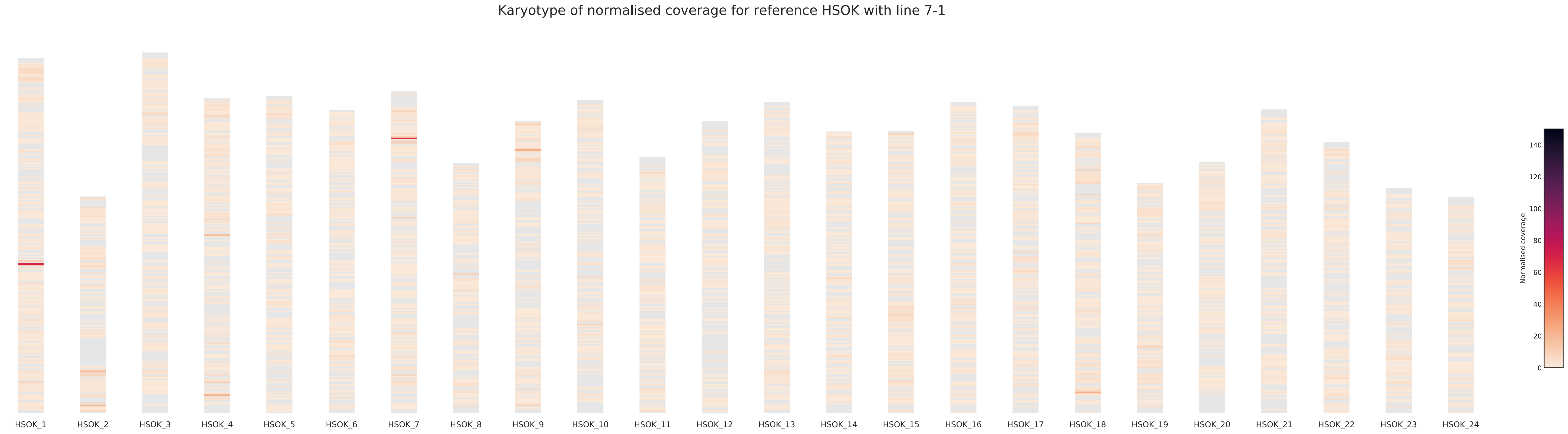
Karyotype of normalised coverage for reference HSOK with line 7-1
HSOK_1
HSOK_2
HSOK_3
HSOK_4
HSOK_5
HSOK_6
HSOK_7
HSOK_8
HSOK_9
HSOK_10
HSOK_11
HSOK_12
HSOK_13
HSOK_14
HSOK_15
HSOK_16
HSOK_17
HSOK_18
HSOK_19
HSOK_20
HSOK_21
HSOK_22
HSOK_23
HSOK_24
Normalised coverage
0
20
40
60
80
100
120
140

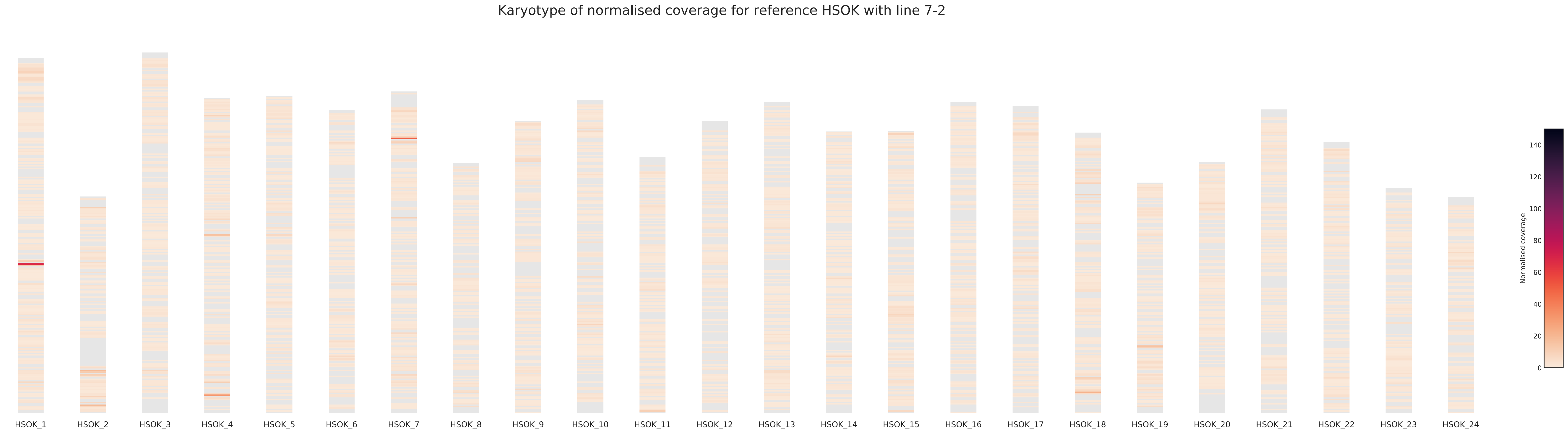
Karyotype of normalised coverage for reference HSOK with line 7-2
HSOK_1
HSOK_2
HSOK_3
HSOK_4
HSOK_5
HSOK_6
HSOK_7
HSOK_8
HSOK_9
HSOK_10
HSOK_11
HSOK_12
HSOK_13
HSOK_14
HSOK_15
HSOK_16
HSOK_17
HSOK_18
HSOK_19
HSOK_20
HSOK_21
HSOK_22
HSOK_23
HSOK_24
Normalised coverage
0
20
40
60
80
100
120
140

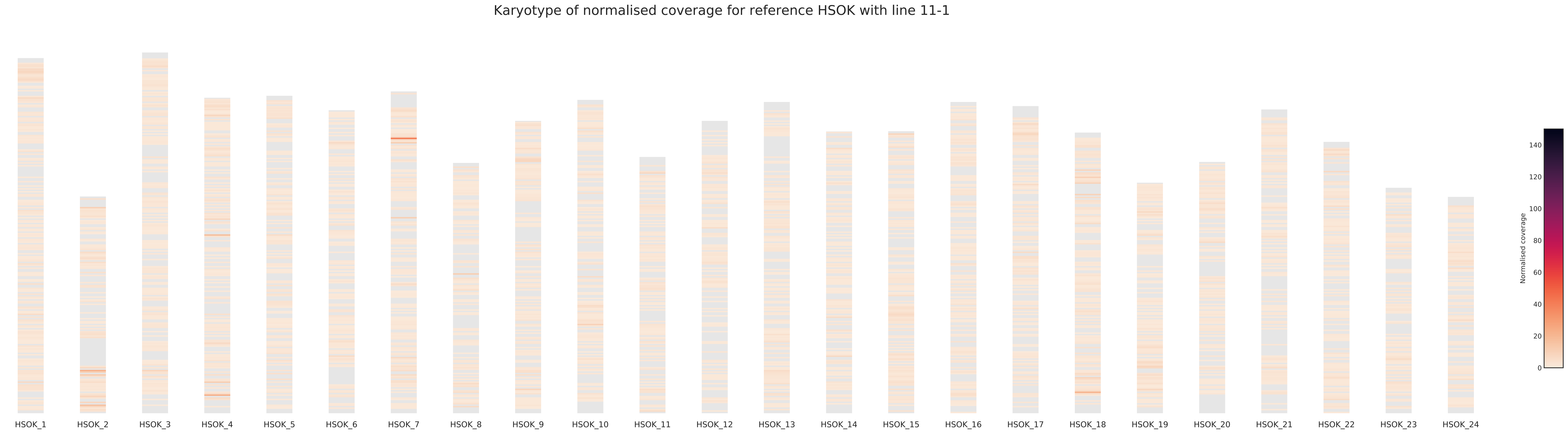
Karyotype of normalised coverage for reference HSOK with line 11-1
HSOK_1
HSOK_2
HSOK_3
HSOK_4
HSOK_5
HSOK_6
HSOK_7
HSOK_8
HSOK_9
HSOK_10
HSOK_11
HSOK_12
HSOK_13
HSOK_14
HSOK_15
HSOK_16
HSOK_17
HSOK_18
HSOK_19
HSOK_20
HSOK_21
HSOK_22
HSOK_23
HSOK_24
Normalised coverage
0
20
40
60
80
100
120
140

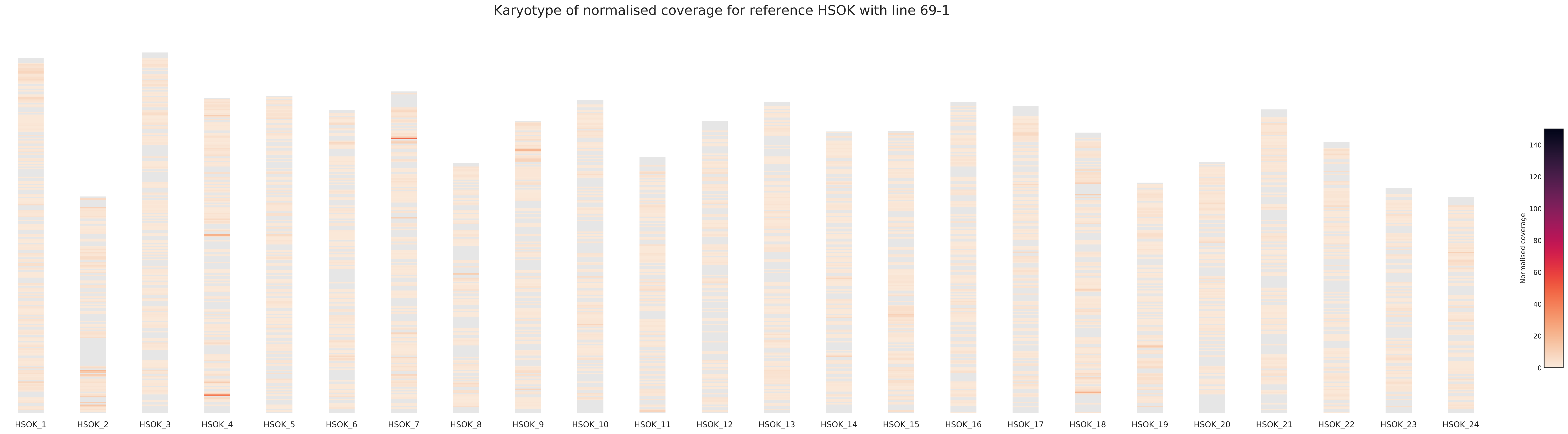
Karyotype of normalised coverage for reference HSOK with line 69-1
HSOK_1
HSOK_2
HSOK_3
HSOK_4
HSOK_5
HSOK_6
HSOK_7
HSOK_8
HSOK_9
HSOK_10
HSOK_11
HSOK_12
HSOK_13
HSOK_14
HSOK_15
HSOK_16
HSOK_17
HSOK_18
HSOK_19
HSOK_20
HSOK_21
HSOK_22
HSOK_23
HSOK_24
Normalised coverage
0
20
40
60
80
100
120
140

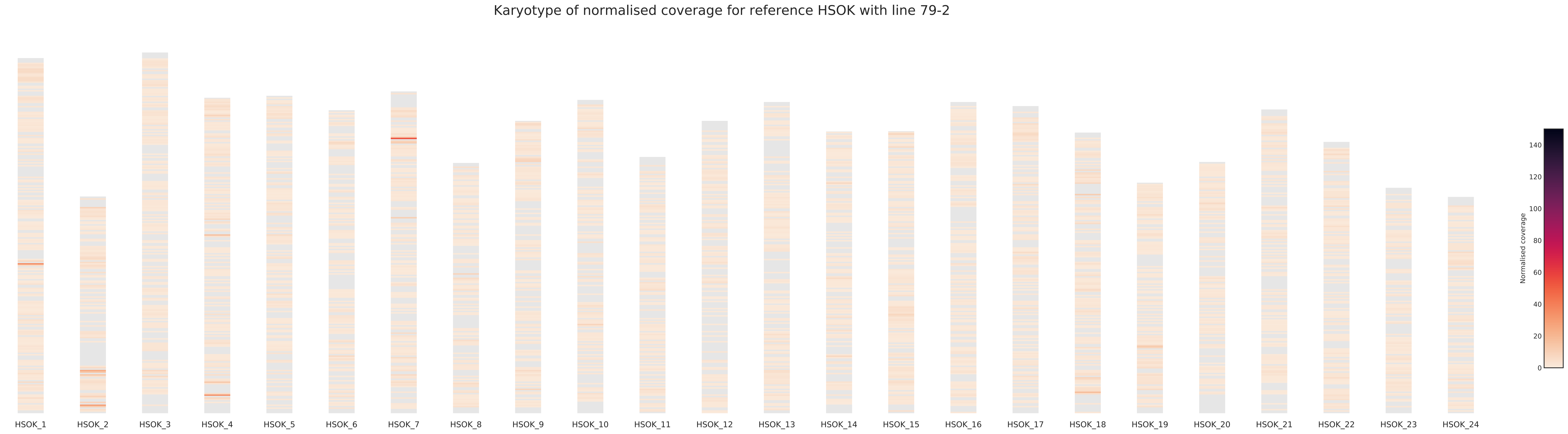
Karyotype of normalised coverage for reference HSOK with line 79-2
HSOK_1
HSOK_2
HSOK_3
HSOK_4
HSOK_5
HSOK_6
HSOK_7
HSOK_8
HSOK_9
HSOK_10
HSOK_11
HSOK_12
HSOK_13
HSOK_14
HSOK_15
HSOK_16
HSOK_17
HSOK_18
HSOK_19
HSOK_20
HSOK_21
HSOK_22
HSOK_23
HSOK_24
Normalised coverage
0
20
40
60
80
100
120
140

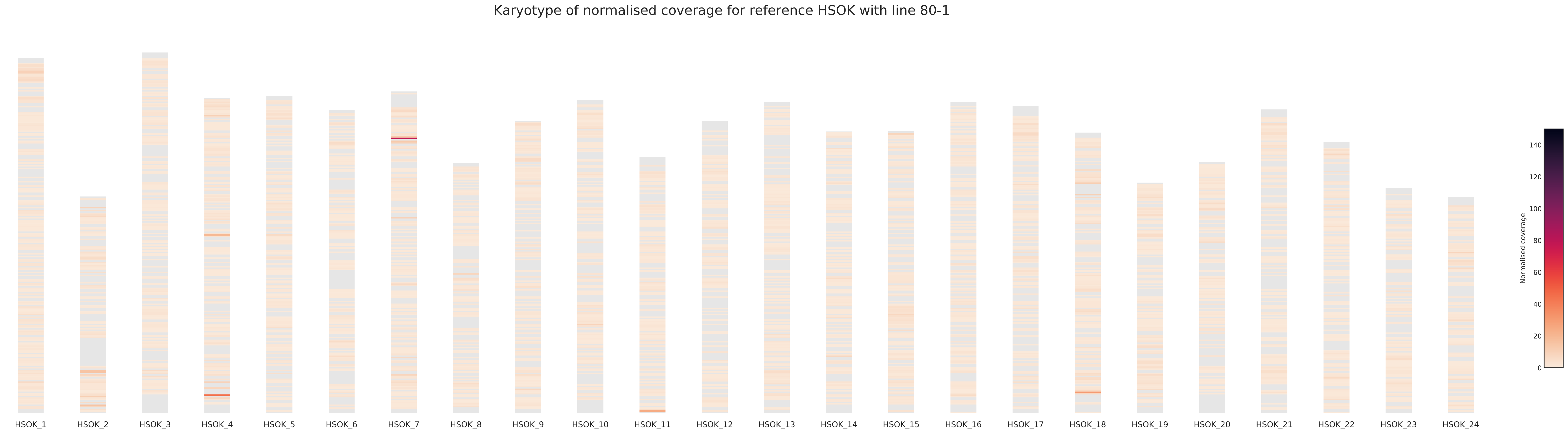
Karyotype of normalised coverage for reference HSOK with line 80-1
HSOK_1
HSOK_2
HSOK_3
HSOK_4
HSOK_5
HSOK_6
HSOK_7
HSOK_8
HSOK_9
HSOK_10
HSOK_11
HSOK_12
HSOK_13
HSOK_14
HSOK_15
HSOK_16
HSOK_17
HSOK_18
HSOK_19
HSOK_20
HSOK_21
HSOK_22
HSOK_23
HSOK_24
Normalised coverage
0
20
40
60
80
100
120
140

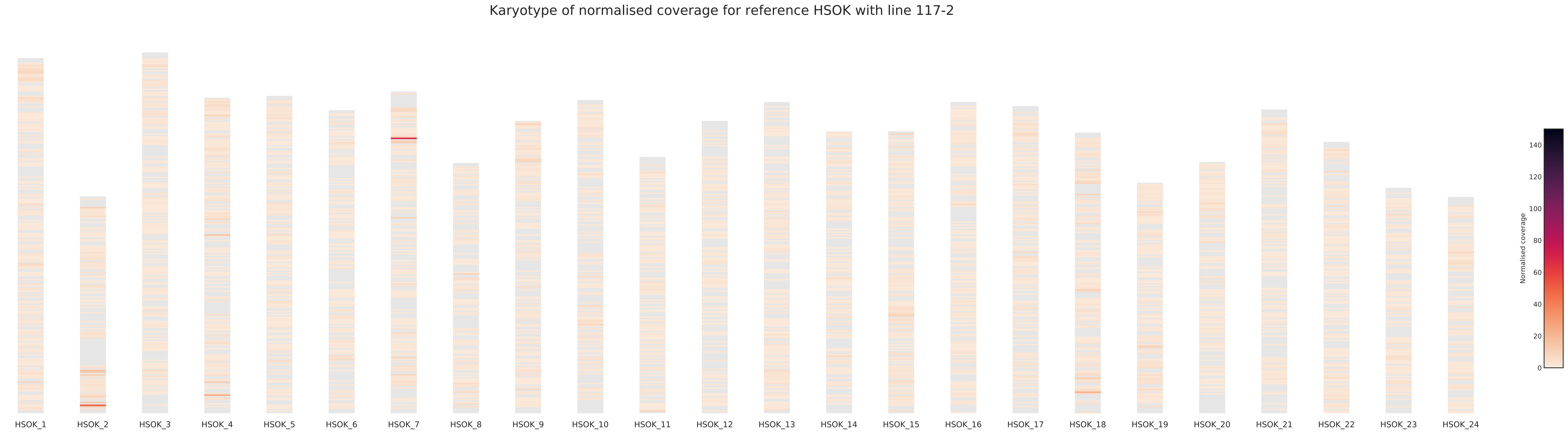
Karyotype of normalised coverage for reference HSOK with line 117-2
HSOK_1
HSOK_2
HSOK_3
HSOK_4
HSOK_5
HSOK_6
HSOK_7
HSOK_8
HSOK_9
HSOK_10
HSOK_11
HSOK_12
HSOK_13
HSOK_14
HSOK_15
HSOK_16
HSOK_17
HSOK_18
HSOK_19
HSOK_20
HSOK_21
HSOK_22
HSOK_23
HSOK_24
Normalised coverage
0
20
40
60
80
100
120
140

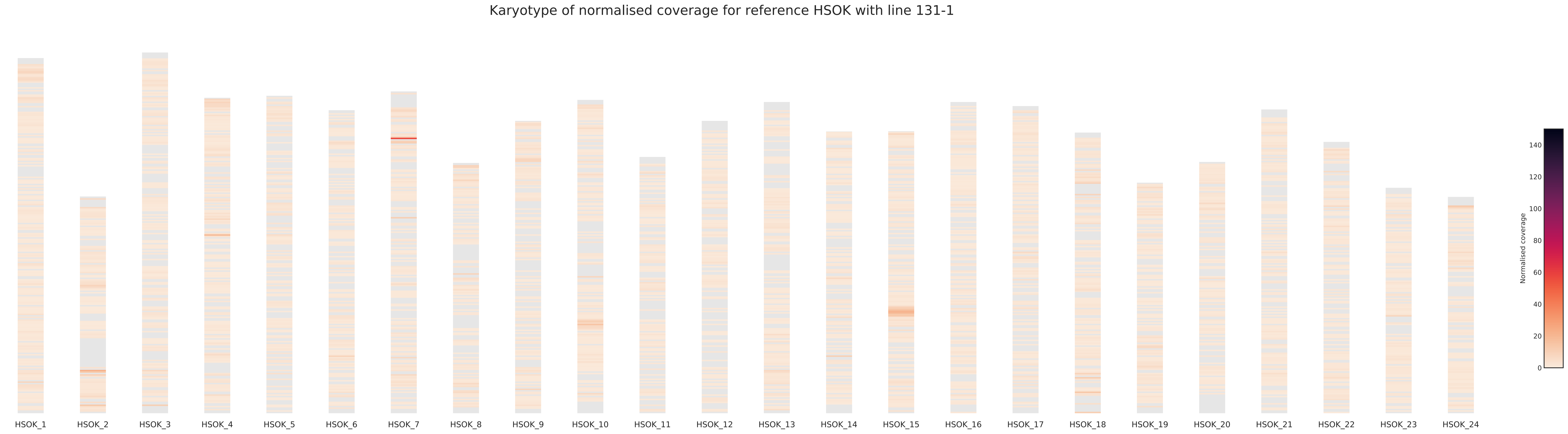

Karyotype of normalised coverage for reference HSOK with line 131-1
HSOK_1
HSOK_2
HSOK_3
HSOK_4
HSOK_5
HSOK_6
HSOK_7
HSOK_8
HSOK_9
HSOK_10
HSOK_11
HSOK_12
HSOK_13
HSOK_14
HSOK_15
HSOK_16
HSOK_17
HSOK_18
HSOK_19
HSOK_20
HSOK_21
HSOK_22
HSOK_23
HSOK_24
Normalised coverage
0
20
40
60
80
100
120
140

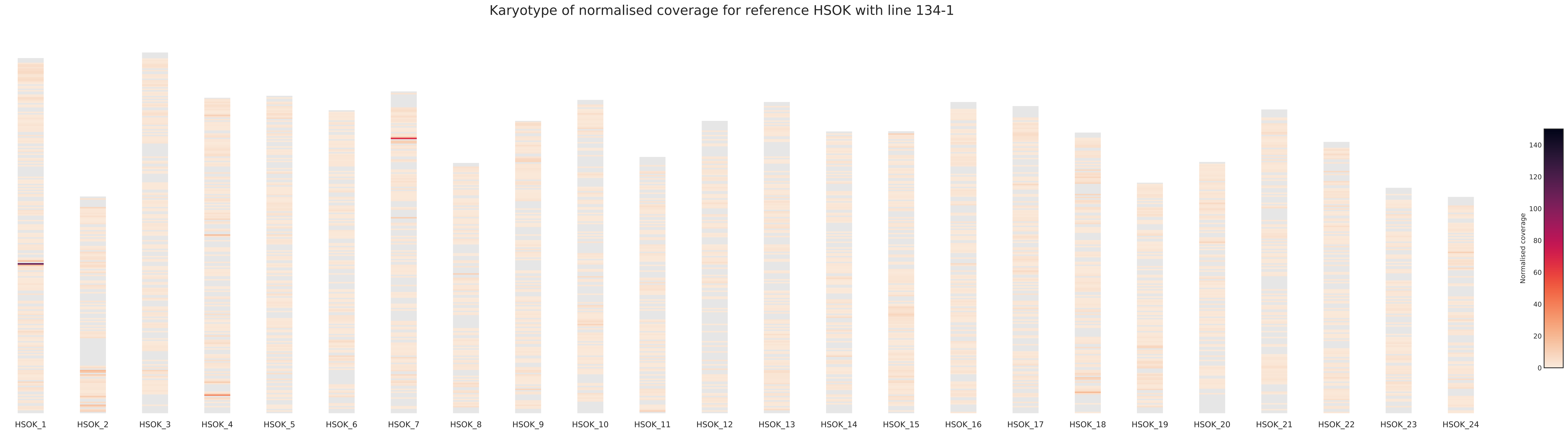
Karyotype of normalised coverage for reference HSOK with line 134-1
HSOK_1
HSOK_2
HSOK_3
HSOK_4
HSOK_5
HSOK_6
HSOK_7
HSOK_8
HSOK_9
HSOK_10
HSOK_11
HSOK_12
HSOK_13
HSOK_14
HSOK_15
HSOK_16
HSOK_17
HSOK_18
HSOK_19
HSOK_20
HSOK_21
HSOK_22
HSOK_23
HSOK_24
Normalised coverage
0
20
40
60
80
100
120
140

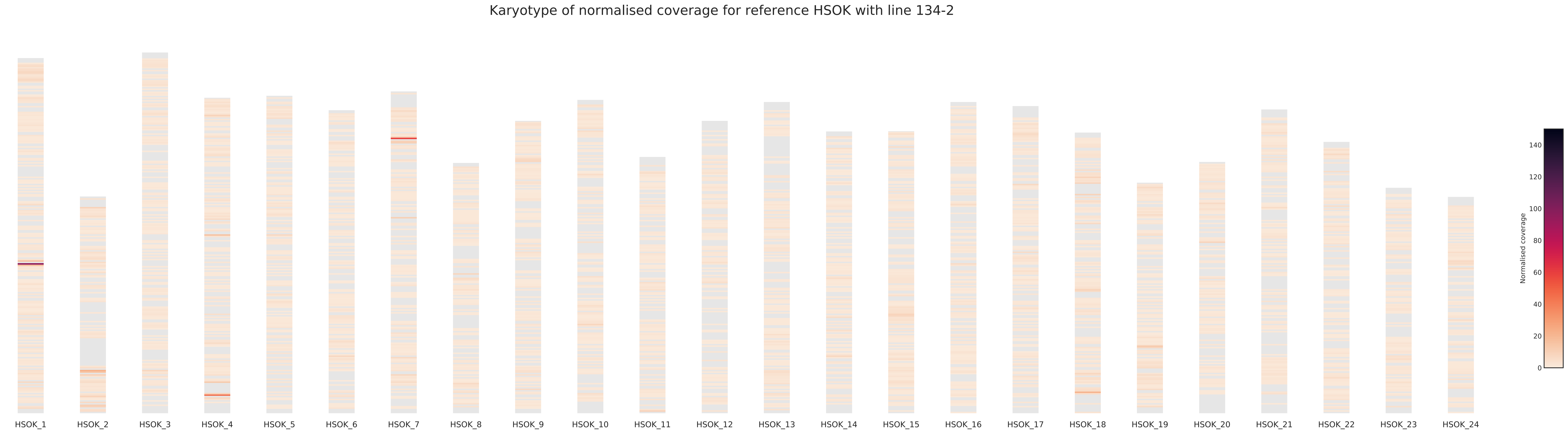

Karyotype of normalised coverage for reference HSOK with line 134-2
HSOK_1
HSOK_2
HSOK_3
HSOK_4
HSOK_5
HSOK_6
HSOK_7
HSOK_8
HSOK_9
HSOK_10
HSOK_11
HSOK_12
HSOK_13
HSOK_14
HSOK_15
HSOK_16
HSOK_17
HSOK_18
HSOK_19
HSOK_20
HSOK_21
HSOK_22
HSOK_23
HSOK_24
Normalised coverage
0
20
40
60
80
100
120
140

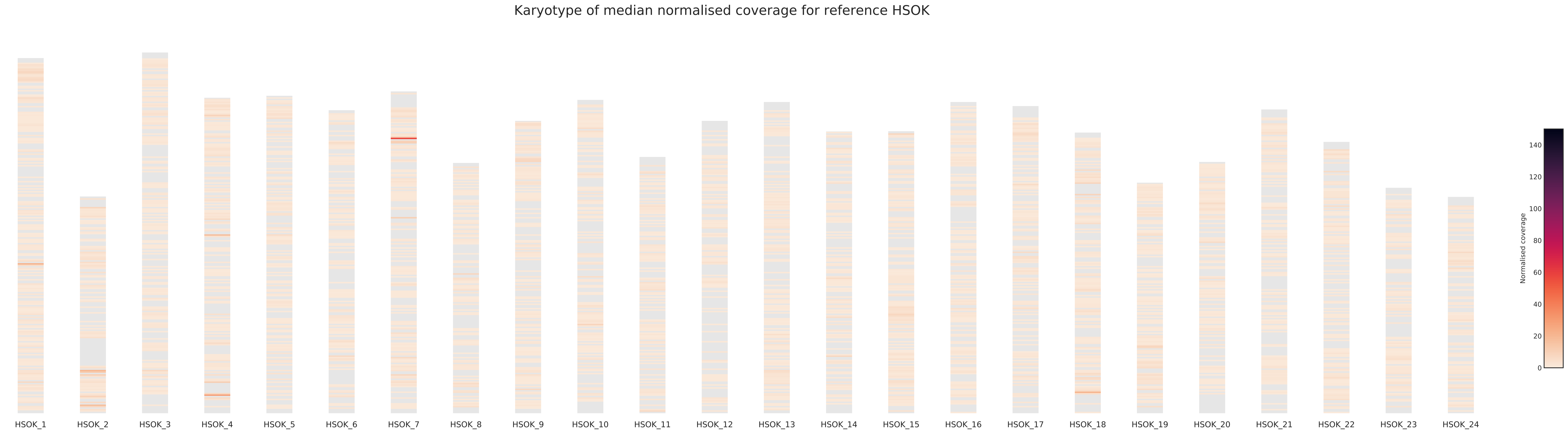

Karyotype of median normalised coverage for reference HSOK
HSOK_1
HSOK_2
HSOK_3
HSOK_4
HSOK_5
HSOK_6
HSOK_7
HSOK_8
HSOK_9
HSOK_10
HSOK_11
HSOK_12
HSOK_13
HSOK_14
HSOK_15
HSOK_16
HSOK_17
HSOK_18
HSOK_19
HSOK_20
HSOK_21
HSOK_22
HSOK_23
HSOK_24
Normalised coverage
0
20
40
60
80
100
120
140
