## Supplementary figures and images for "Genomic variations and epigenomic landscape of the Medaka Inbred Kiyosu-Karlsruhe (MIKK) panel"

### Supplementary file 2

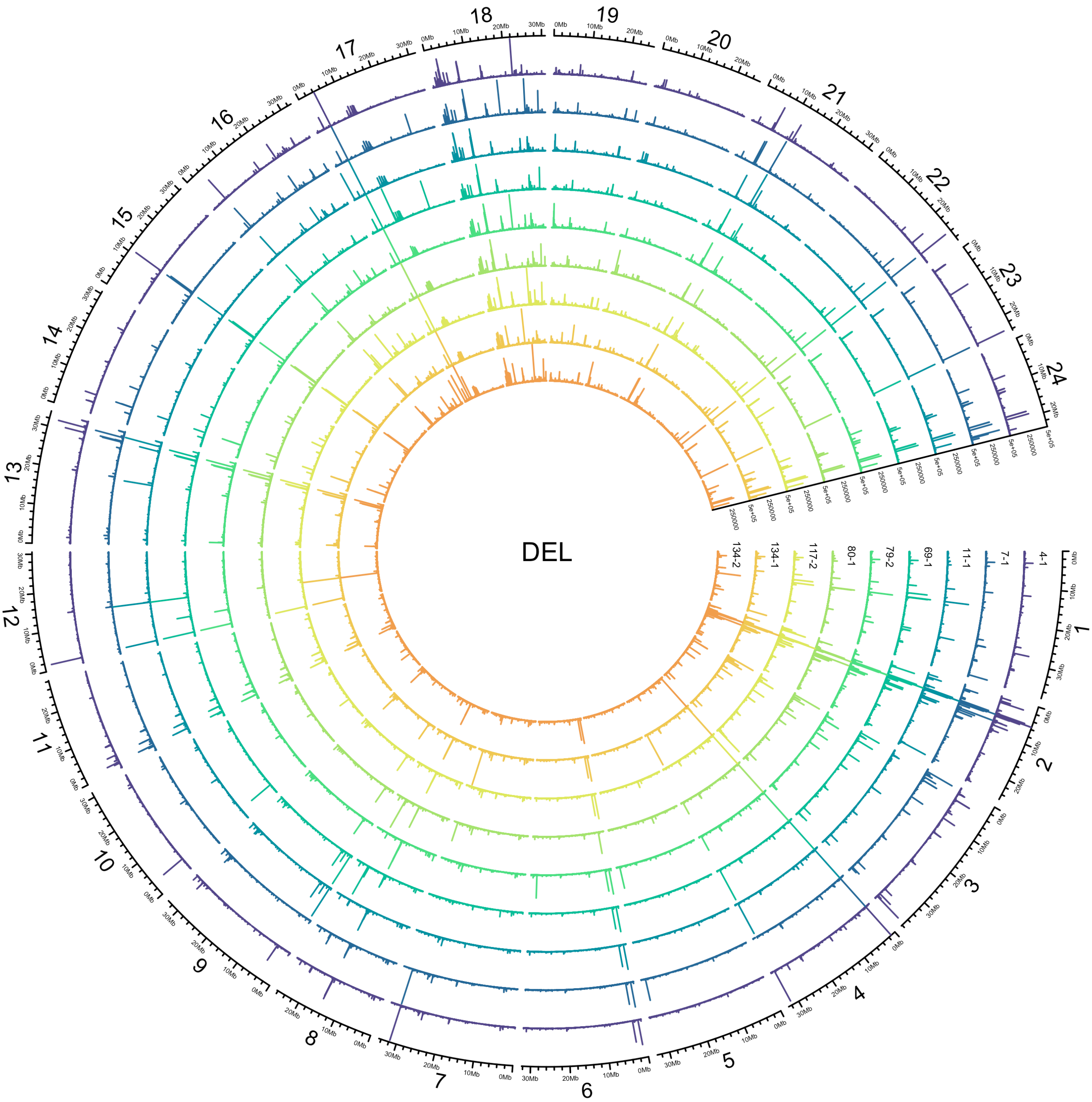
DEL
134-2
134-1
117-2
80-1
79-2
69-1
11-1
7-1
4-1
1
2
3
4
5
6
7
8
9
10
11
12
13
14
15
16
17
18
19
20
21
22
23
24
0Mb
10Mb
20Mb
30Mb
250000
5e+05

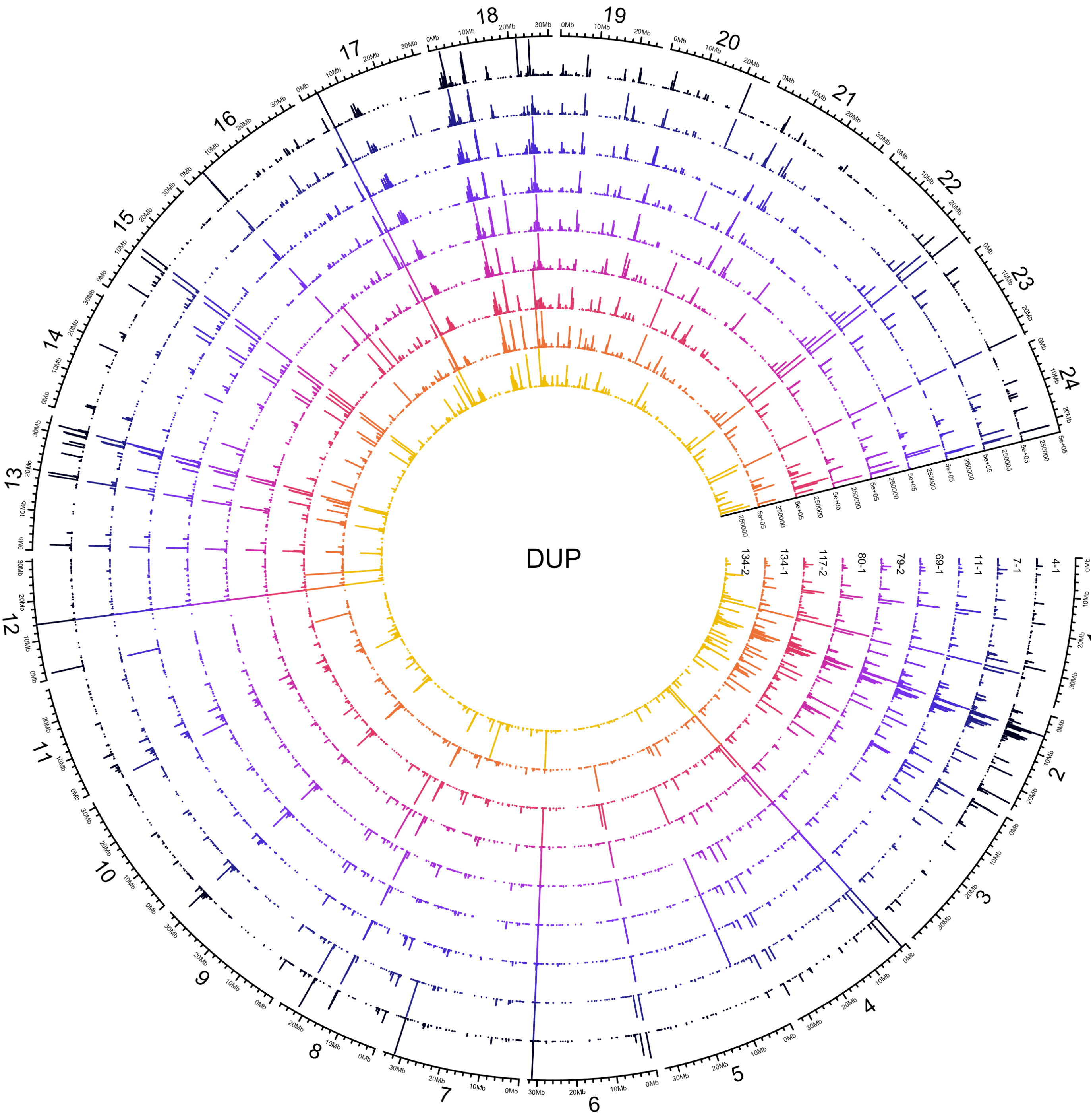
DUP
134-2
134-1
117-2
80-1
79-2
69-1
11-1
7-1
4-1
1
2
3
4
5
6
7
8
9
10
11
12
13
14
15
16
17
18
19
20
21
22
23
24

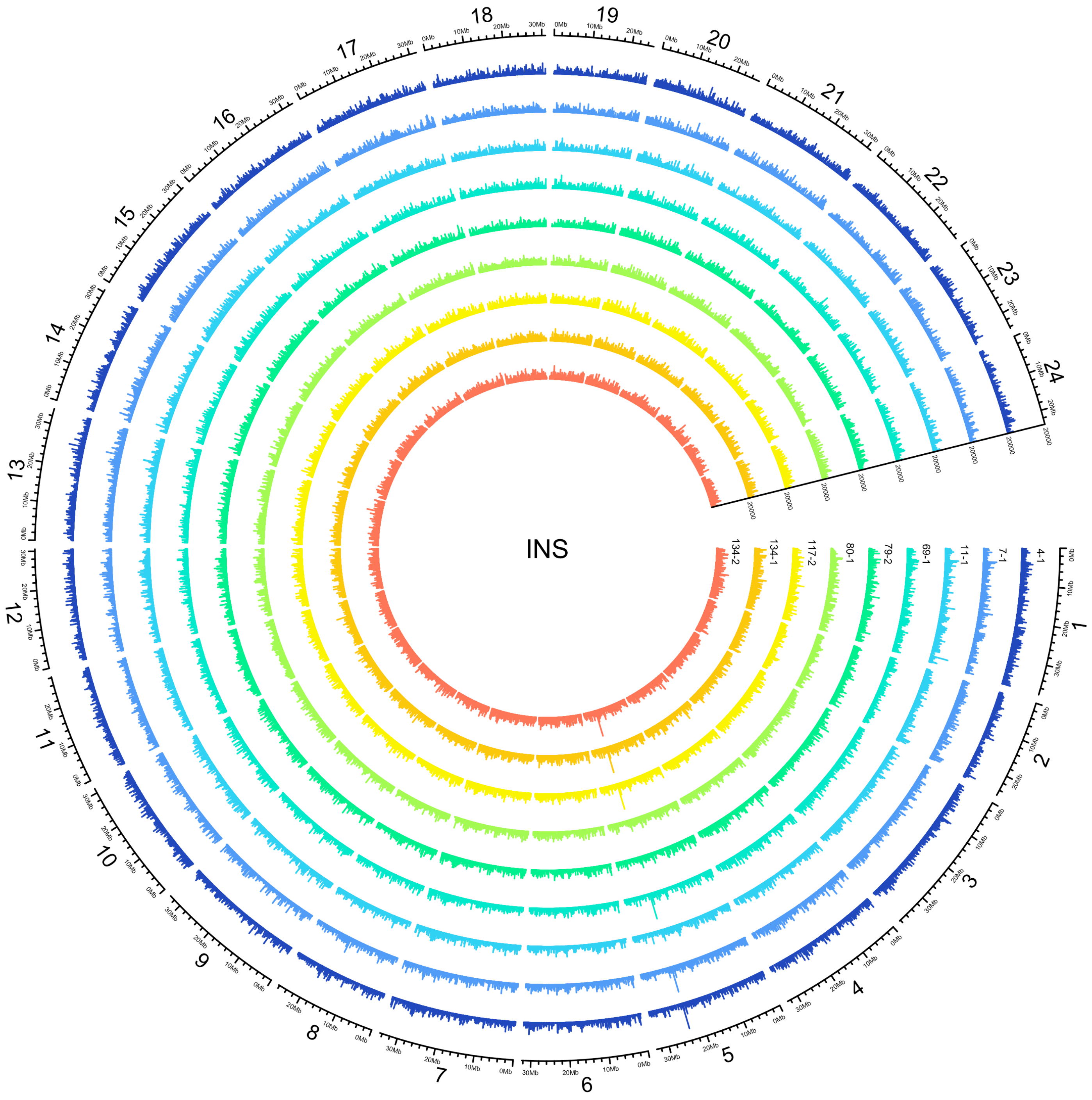
INS
134-2
134-1
117-2
80-1
79-2
69-1
11-1
7-1
4-1
20000
1
2
3
4
5
6
7
8
9
10
11
12
13
14
15
16
17
18
19
20
21
22
23
24

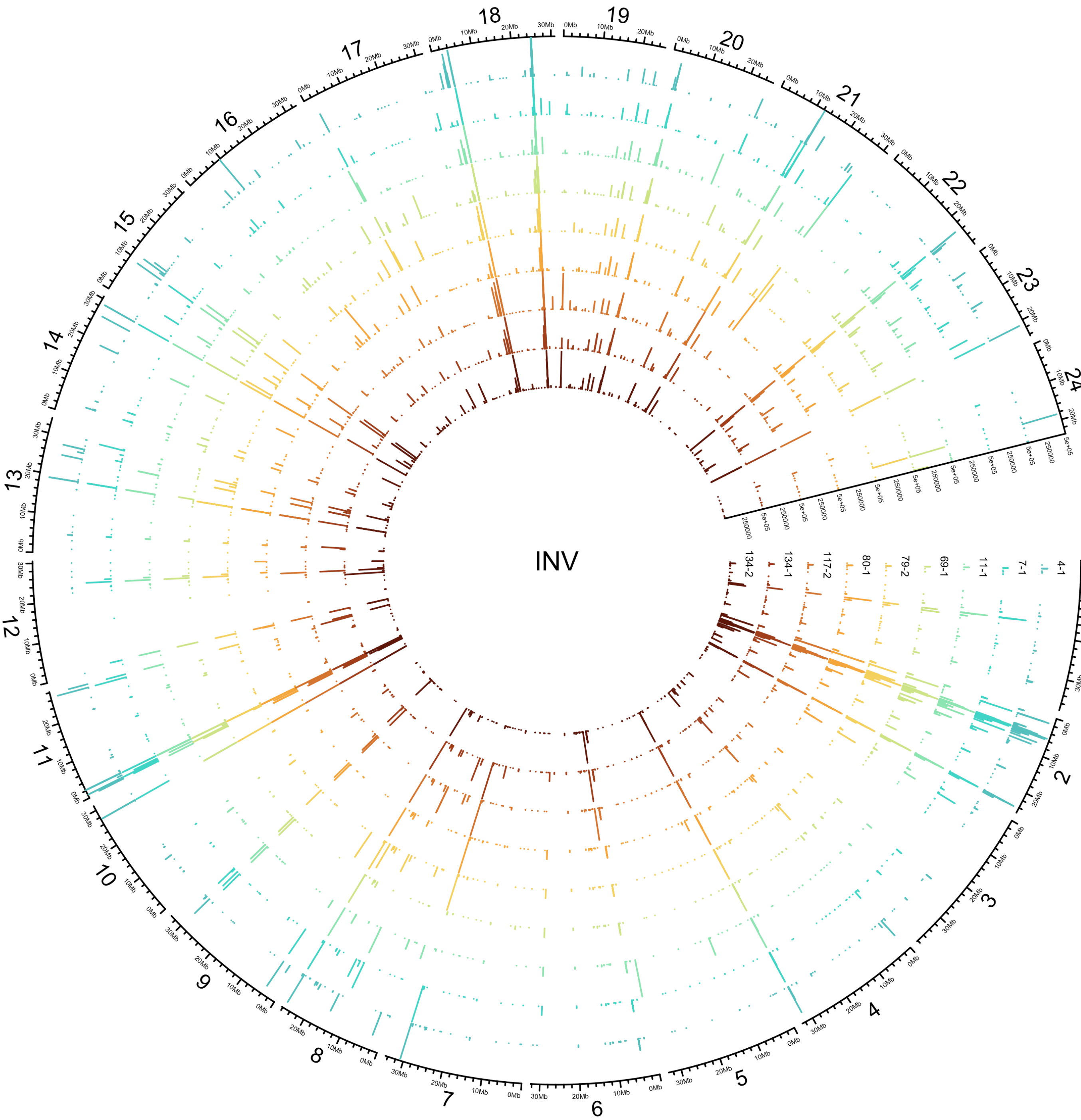

INV
134-2
134-1
117-2
80-1
79-2
69-1
11-1
7-1
4-1

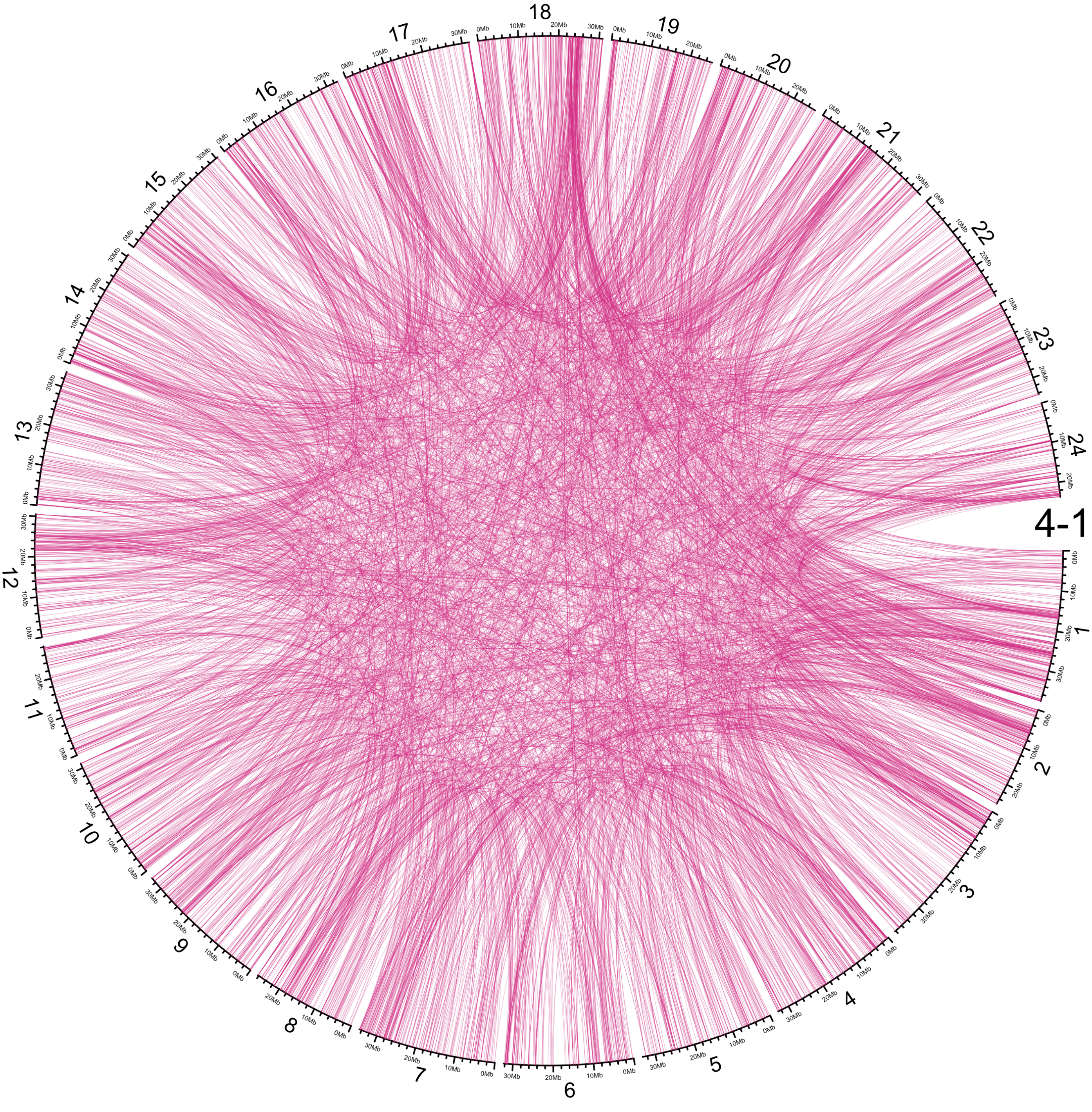
4-1
1
2
3
4
5
6
7
8
9
10
11
12
13
14
15
16
17
18
19
20
21
22
23
24

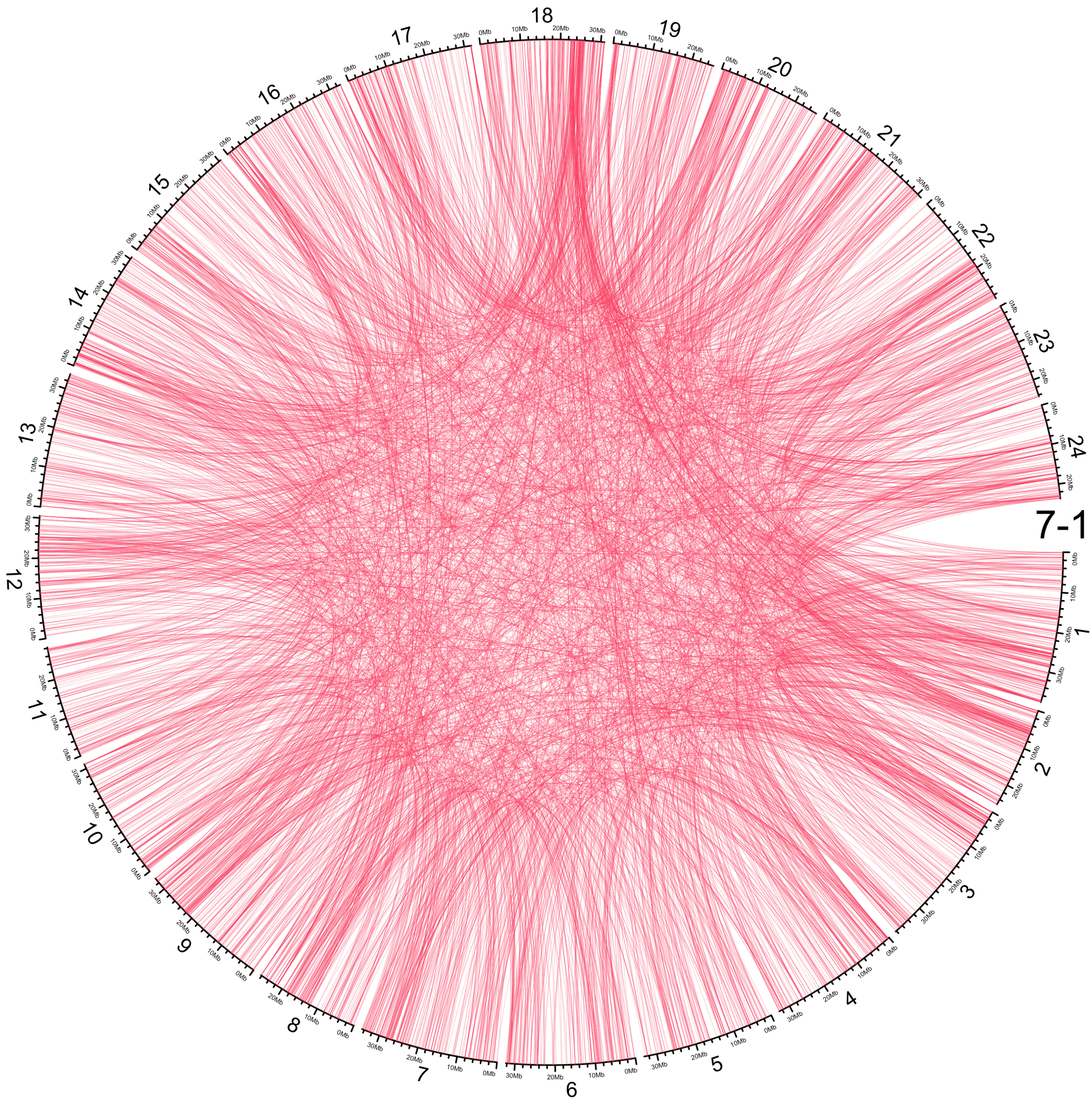
7-1
1
2
3
4
5
6
7
8
9
10
11
12
13
14
15
16
17
18
19
20
21
22
23
24

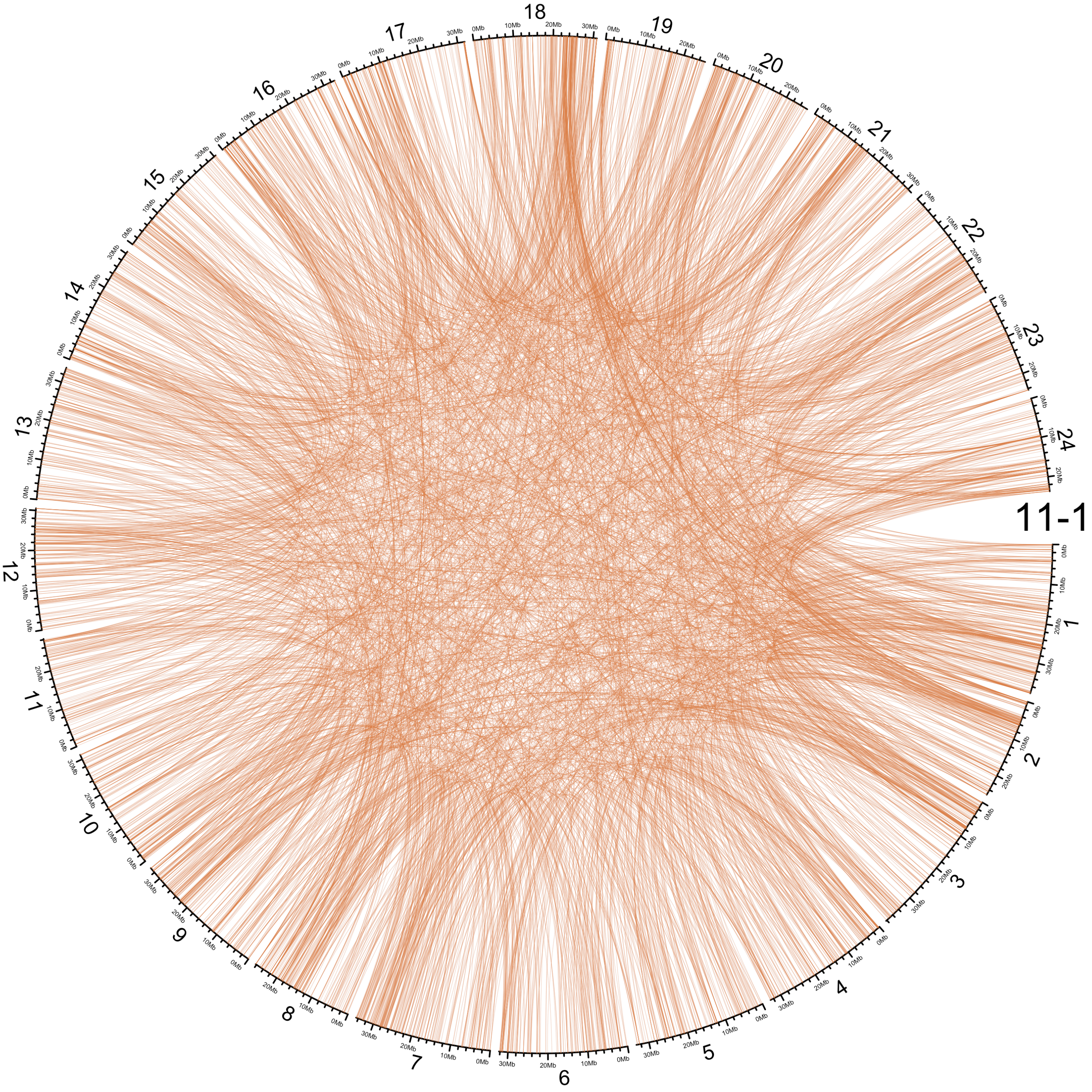
11-1
1
2
3
4
5
6
7
8
9
10
11
12
13
14
15
16
17
18
19
20
21
22
23
24

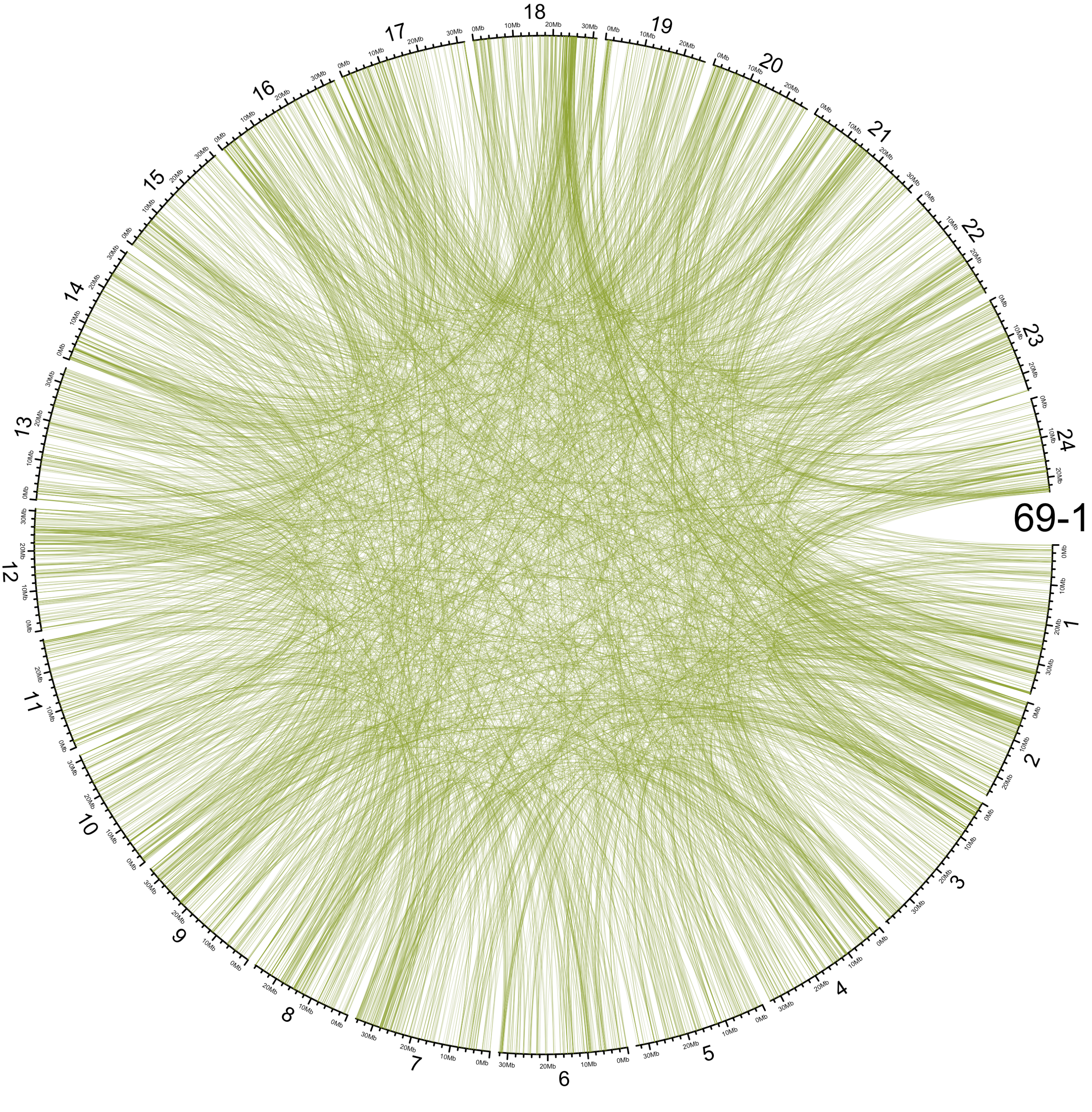
69-1
1
2
3
4
5
6
7
8
9
10
11
12
13
14
15
16
17
18
19
20
21
22
23
24

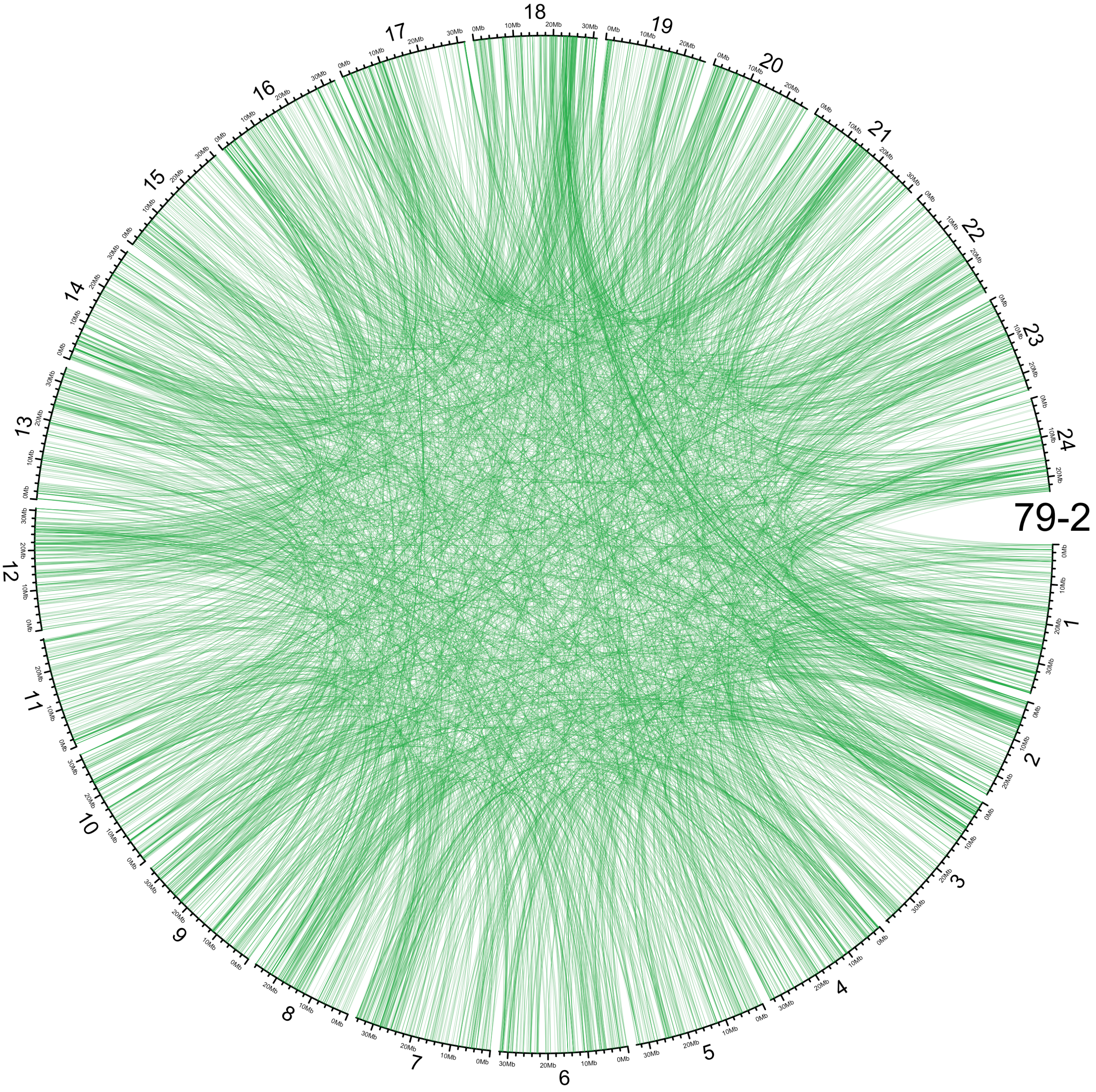
79-2
1
2
3
4
5
6
7
8
9
10
11
12
13
14
15
16
17
18
19
20
21
22
23
24

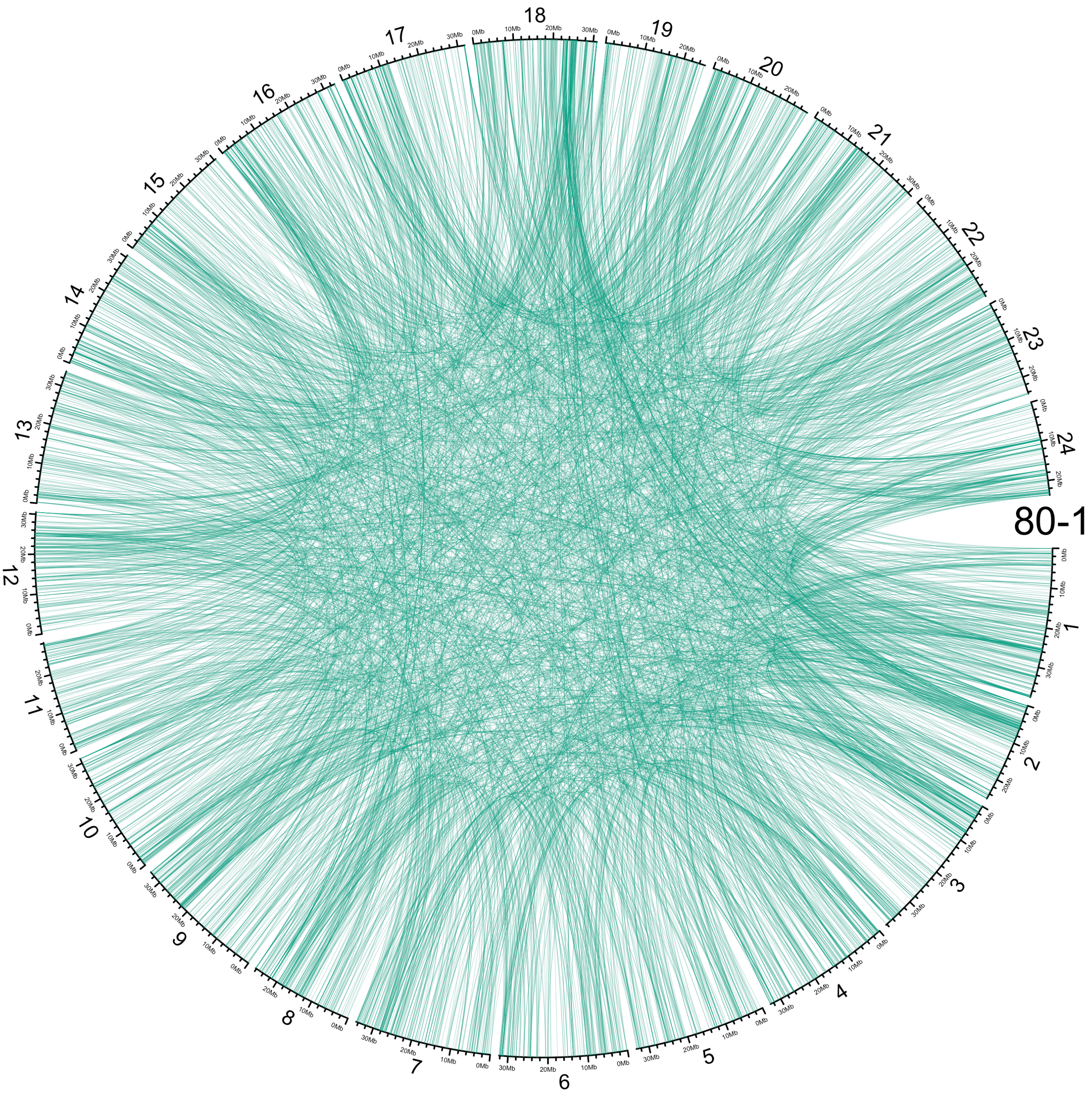
1
2
3
4
5
6
7
8
9
10
11
12
13
14
15
16
17
18
19
20
21
22
23
24
80-1

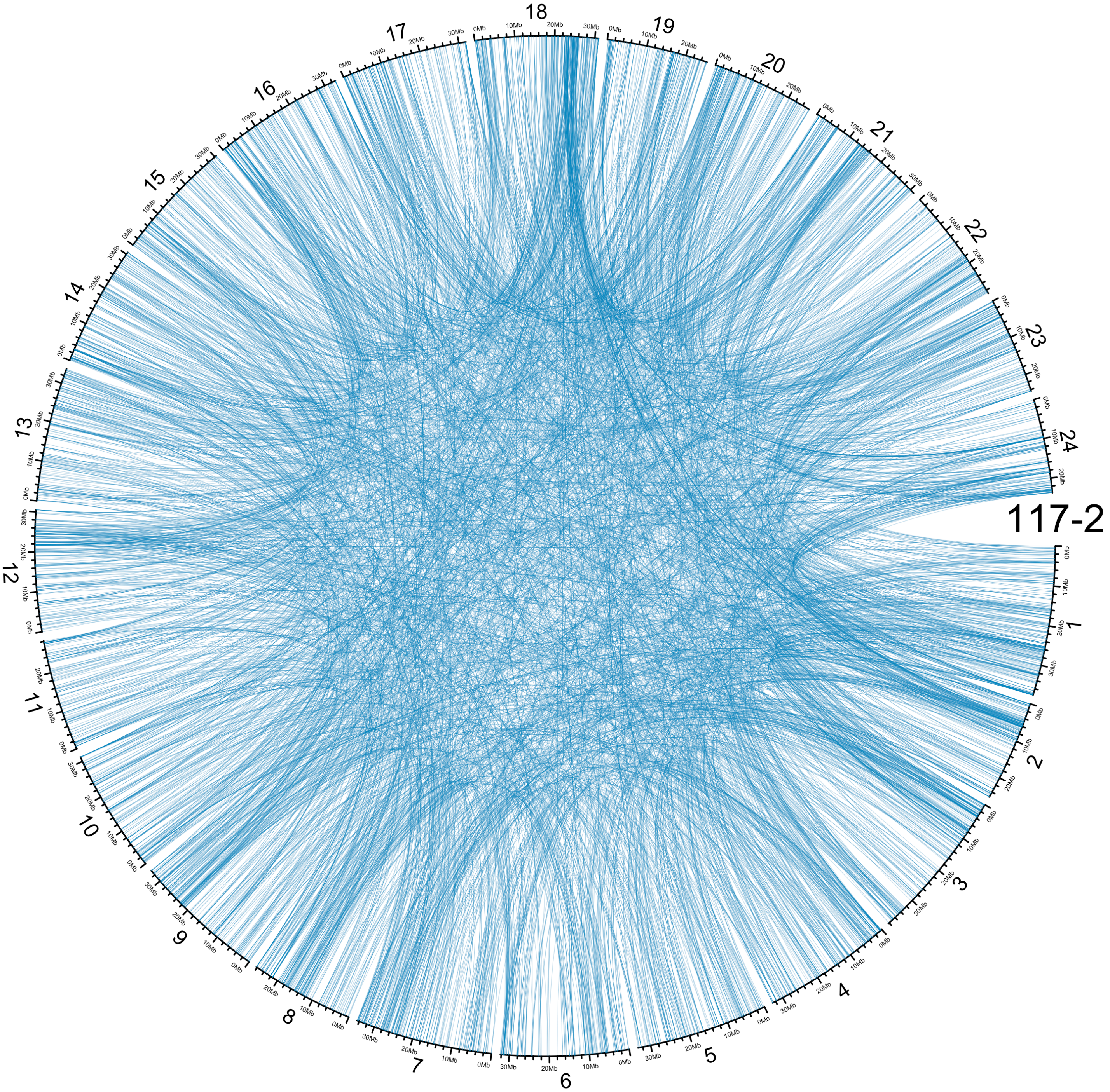
117-2
1
2
3
4
5
6
7
8
9
10
11
12
13
14
15
16
17
18
19
20
21
22
23
24

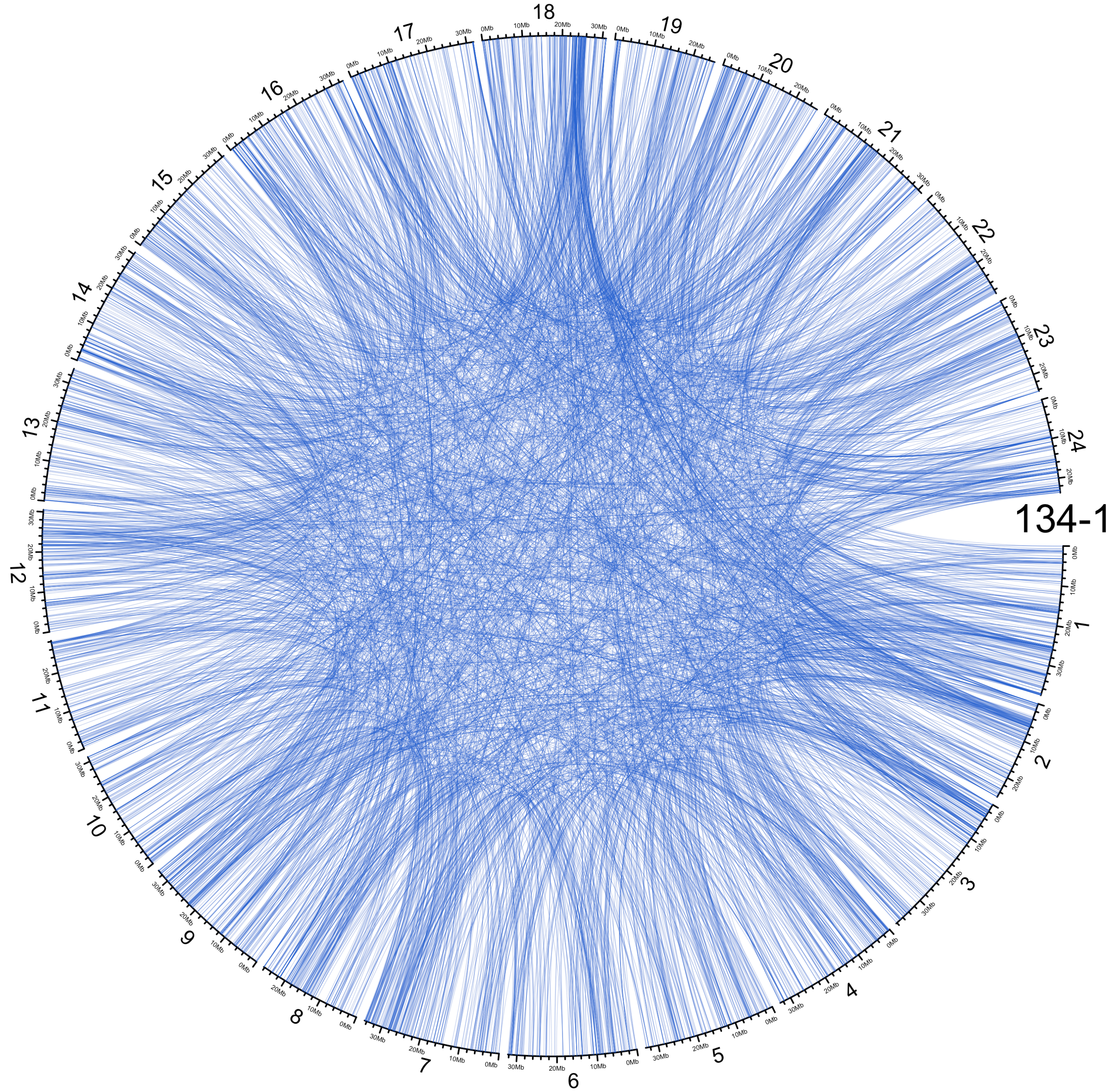
134-1
1
2
3
4
5
6
7
8
9
10
11
12
13
14
15
16
17
18
19
20
21
22
23
24

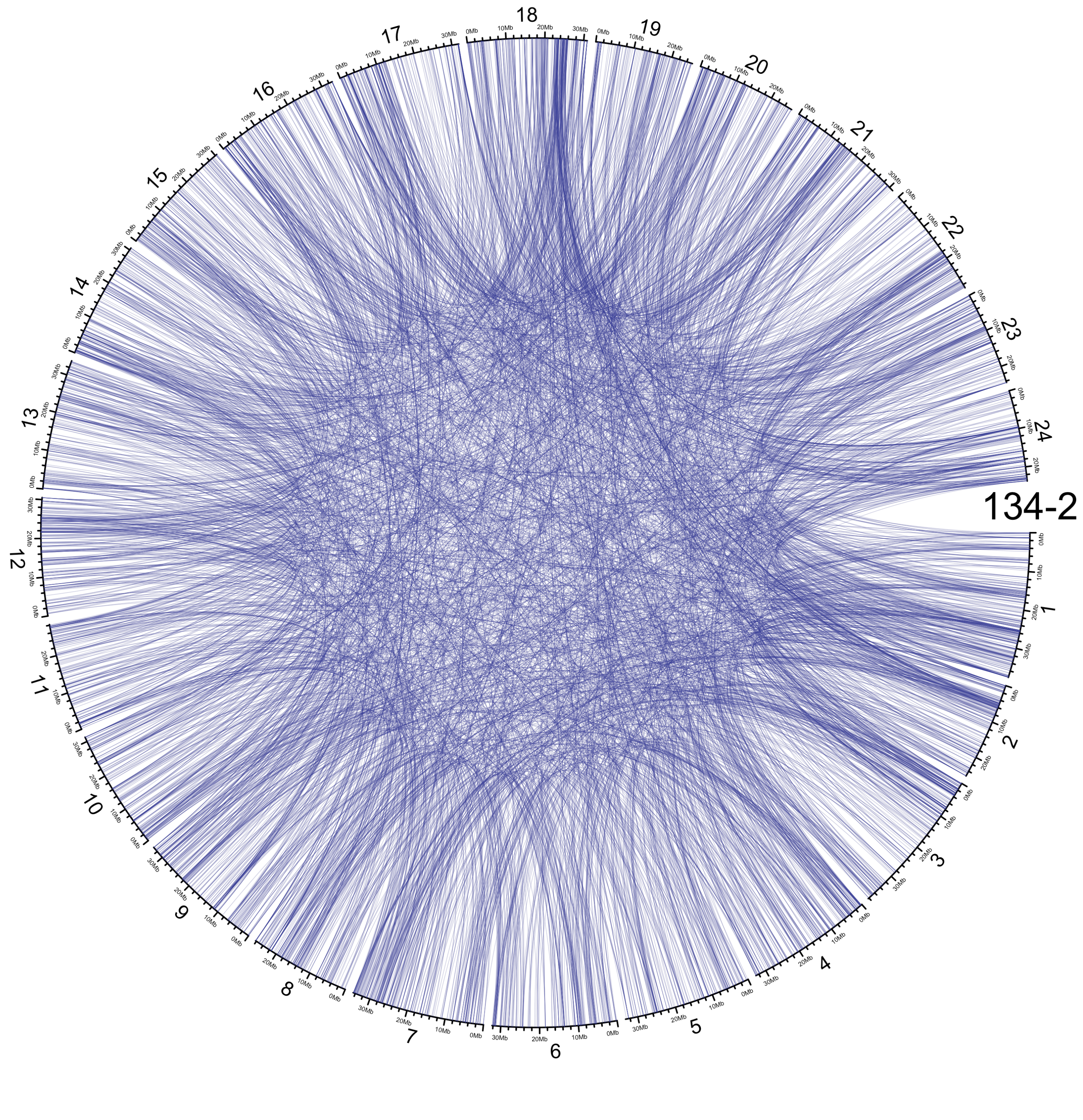
134-2
1
2
3
4
5
6
7
8
9
10
11
12
13
14
15
16
17
18
19
20
21
22
23
24
